# AAV-Delivered RNAi Targeting Mutant LDB3 Prevents and Reverses Myofibrillar Myopathy through Mechanosignaling Restoration

**DOI:** 10.64898/2026.03.28.715031

**Authors:** Pankaj Pathak, Jessica Palmeri, Jessica Hale, Anna Sabu-Kurian, Morteza Peiravi, Danielle A. Springer, Yan Li, Kory R. Johnson, Ami Mankodi

## Abstract

The autosomal dominant p.Ala165Val mutation in LIM Domain Binding Protein 3 (LDB3) causes myofibrillar myopathy marked by Z-disc disruption, accumulation of filamin-C (FLNc) and chaperone proteins, and progressive muscle weakness. We previously showed that this mutation interferes with the LDB3-protein kinase C alpha (PKCα)-FLNc mechanosensing axis and impairs chaperone-assisted selective autophagy (CASA), establishing a gain-of-function mechanism. In this study, we examined whether mutant allele-specific knockdown could reverse the disease or mitigate disease progression *in-vivo*. A single intramuscular-injection of an AAV9-delivered microRNA-based shRNA produced substantial knockdown of mutant *Ldb3* transcripts and protein in *Ldb3^Ala165Val/+^* knock-in mice treated either before or after the onset of pathology. Treatment after disease onset reduced filamin-C and CASA protein aggregates and improved muscle strength, whereas early intervention prevented development of molecular and histological features of myopathy. Phosphoproteomic profiling further showed broad remodeling of dysregulated phosphorylation networks, including restoration of PKCα-responsive sites and normalization of altered sarcomeric and cytoskeletal signaling observed in *Ldb3^Ala165Val/+^* mice. These findings identify disruption of the LDB3-PKCα-FLNc mechanosensing pathway as a central disease driver and suggest that restoring this signaling axis may complement mutant allelespecific RNA interference (RNAi). Overall, our results support RNAi as a promising therapeutic strategy for dominant LDB3-related myofibrillar myopathy.

## Introduction

Rare diseases, increasingly acknowledged as a significant challenge to global public health collectively impact an estimated 300 million individuals worldwide, including approximately 30 million people in the United States, nearly 1 in 10 Americans.^1,2^ Rare diseases are often serious condition with devastating effects on patients and families. Because disease-modifying therapies remain unavailable for most rare diseases, patients living with these individually uncommon, yet collectively prevalent conditions continue to face a substantial and urgent unmet medical need. A majority (∼80%) of rare diseases arise from single-gene defects, providing a strong biological rationale for gene-directed therapeutic interventions.

Gene therapy has proven effective for monogenic disorders caused by loss-of-function mutations through gene replacement strategies. However, dominantly inherited diseases driven by toxic gain-of-function alleles cannot be corrected by gene augmentation alone. In these settings, transcript-targeting approaches such as antisense oligonucleotides (ASOs) and RNA interference provide a rational means to selectively suppress pathogenic transcripts while preserving essential wild-type (WT) protein expression.^3,4^

Although RNA-based therapeutics have advanced rapidly with several siRNA agents now approved, clinical success has largely been restricted to hepatic targets due to the intrinsic liver tropism of systemically delivered nucleic acids.^4^ ASOs have demonstrated efficacy in select neuromuscular disorders, but their short tissue half-life, need for repeated dosing, and variable muscle distribution remain limitations.^5^ On the other hand, adeno-associated virus (AAV)-mediated RNAi enables sustained intramuscular expression from a single administration, supporting durable transcript suppression and improved tissue targeting.^6^ Achieving efficient gene modulation in skeletal muscle therefore remains a key translational objective, for which localized AAV delivery provides a rigorous platform to establish therapeutic potential and inform future systemic strategies.

Myofibrillar myopathy (MFM) is a group of rare but clinically severe monogenetic diseases primarily affecting skeletal muscle with no approved therapy. Patients typically present with progressive muscle weakness and wasting with eventual loss of independent living, respiratory failure, and cardiomyopathy.^7^ MFMs are caused by dominantly inherited mutations in the genes encoding Z-disc proteins, with pathogenic mutations in at least 15 genes have been identified. All MFMs are primarily defined by muscle pathological changes of myofibrillar dissolution associated with disintegration of the Z-disc and aggregation of degraded myofibrillar proteins.^8^ We and others have shown that the disease mechanism is shared among MFMs, involving the impairment of chaperone assisted selective autophagy (CASA), a mechanosensitive protein quality-control pathway essential for the maintenance of the Z-disc integrity in contracting skeletal muscle fibers.^9–12^

One recurrent form of MFM is caused by the p.Ala165Val mutation in LIM domain binding 3 (LDB3; HGNC 15710; rs121908334; NM_001080114.2:c.494 C > T, NP_001073583.1), a PDZ-LIM protein localized to Z-disc.^10^ Previously, we showed that, contrary to prevailing thinking for disease mechanism of protein aggregation in dominant neurodegenerative disorders, the LDB3 p.Ala165Val protein does not seed the sarcomeric protein aggregation through its misfolding or self-aggregation. Rather, it triggers aggregation of filamin C, another MFM-associated Z-disc protein, and its chaperones in the stress-induced CASA pathway at the Z-disc of skeletal muscle fibers in *Ldb3^Ala165Val/+^* knock-in mice.^10^ We also identified new functional roles for LDB3 in recruiting key proteins to strategic sarcomeric Z-disc sites, facilitating mechanosensing, and modulating dynamic signaling pathways mediated by PKCα in striated muscle. The *Ldb3^Ala165Val/+^* knock-in mice reproduce the typical MFM phenotype, including progressive muscle weakness and associated pathology that is seen in the patients carrying this mutation. LDB3 haploinsufficiency is unlikely, as protein levels measured by immunoblotting showed no clear difference in the skeletal muscle between *Ldb3^Ala165Val/+^* knock-in mice and WT littermates, reminiscent of that reported in patients compared with healthy controls.^13^ In addition, heterozygous *Ldb3* mice display no obvious phenotype and have normal survival.^14,15^ Together, these observations support the development of allele-specific silencing as a potential gene therapy strategy in the MFM caused by the LDB3 p.Ala165Val mutation.

To enable mechanistic and therapeutic studies in a robust preclinical setting, we employed an allele specific RNA interference (AS-RNAi) strategy to evaluate the treatment benefit in *Ldb*3*^Ala165Val/+^* mice using a single intramuscular (i.m.) administration of two different AAV9-shRNA-miR constructs delivered either at an early stage (i.e., preceding the obvious clinical and pathological phenotype) or during manifesting progressive stage of disease. In both intervention paradigms, the RNAi intervention resulted in a remarkable rescue of disease phenotypes including improvements in muscle histopathology, contractile function, and molecular features associated with LDB3-related MFM, with therapeutic benefit observed across disease stages. Further phosphoproteomic profiling uncovered the underlying treatment-associated changes in muscle signaling pathways. Together, these findings also validated the important role of CASA pathway and LDB3-PKCα-Filamin C mechanosensing restoration in the RNAi-mediated therapeutic rescue of MFM. Overall, these data establish proof of principle for RNAi-mediated targeting of mutant LDB3 as a therapeutic strategy as well as a prospective treatment for patients with LDB3-MFM, in addition also identifying promising histological and molecular features with potential utility as treatment response biomarkers in a rare neuromuscular disease with substantial unmet medical need.

## RESULTS

### Design and *in-vitro* validation of siRNA targeting mutant *Ldb3* allele

To target the mutant *Ldb3* mRNA for degradation, 19 different siRNAs were designed to span and include the disease-causing C>U (p.Ala165Val) point mutation (Figure 1A). To evaluate the efficacy of candidate siRNAs to preferentially knockdown mutant LDB3, HEK-293 cells were transfected with a plasmid expressing WT or mutant murine LDB3 and five out of the 19 siRNA prioritized based on established sequence features associated with siRNA efficacy and an algorithm (Figure 1A, Table S1).^16–18^ Two siRNAs (si10 and si16) were selected for subsequent *in-vivo* efficacy studies in *Ldb*3*^Ala165Val/+^* mice based on *in-vitro* parallel cell transfection experiments showing strong preferential knockdown effect on the mutant LDB3 protein. Quantification using an antibody recognizing total LDB3 showed a significant suppression of mutant compared with WT-protein (Mean ± SD: mutant LDB3, 0.28 ± 0.04 vs. WT-LDB3, 0.88 ± 0.05, for si10; and mutant LDB3, 0.24 ± 0.02 vs. WT-LDB3, 0.54 ± 0.03, for si16) (Figure 1B and C), which was supported by corresponding *Ldb3* allele expression analyses (Mean ± SD: mutant allele,0.42 ± 0.04 vs. WT-allele, 0.80 ± 0.03, for si10; and mutant allele, 0.41 ± 0.07 vs. WT-allele, 0.53 ± 0.03, for si16) (Figure 1D and E).

**Figure 1.**
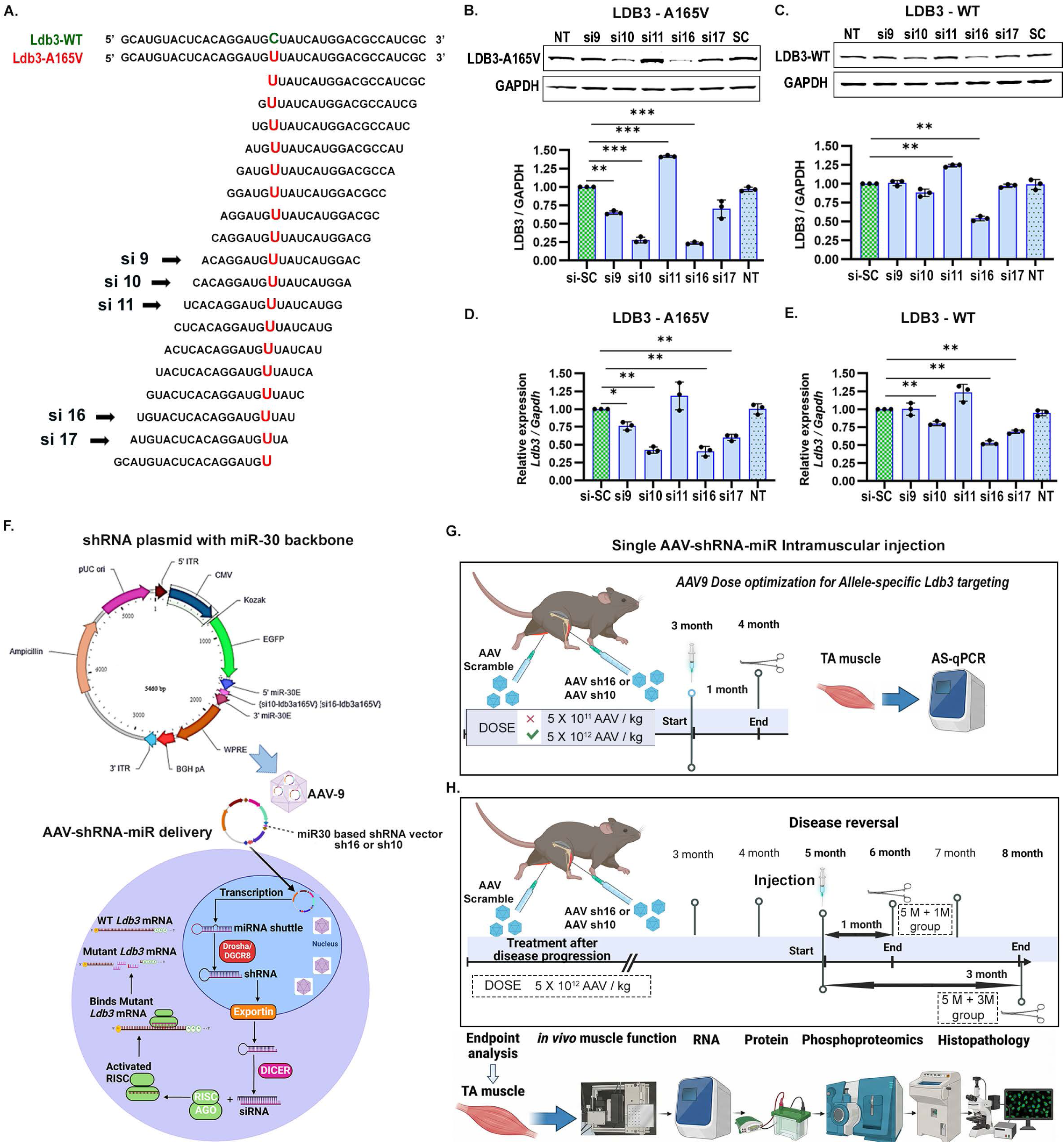
Allele-specific RNAi design, functional validation, and AAV9-mediated delivery targeting *Ldb3-*A165V. (A) WT and A165V mutant *Ldb3* mRNA sequences are shown, highlighting the mutation site (red). Nineteen candidate siRNAs were designed to target this region; the five selected sequences (sense strand), chosen based on computational predictions and prior evidence, are indicated by arrows. siRNAs were numbered according to their position relative to the mismatch with the WT sequence and evaluated in HEK-293 cells expressing either WT or mutant LDB3. (B and C) Immunoblot analysis performed 48 h post-transfection showing effect of the siRNAs on total LDB3 protein levels in mutant- and WT-LDB3 expressing cells (n = 3 per group), including si-SC and NT controls. (D and E) AS-qPCR quantification of mutant and WT *Ldb3* transcripts (n = 3 per group). Transcript levels were normalized to *Gapdh* using the ΔΔCt method and immunoblots were normalized to GAPDH. (F) Schematic of the miR30-based shRNA construct encoding either si10-Ldb3-A165V or si16-Ldb3-A165V (top). The construct was packaged into AAV9 for intramuscular delivery. Following cellular entry, shRNAmiR transcripts are processed through the endogenous microRNA pathway to generate mature shRNAs targeting mutant *Ldb3* mRNA (bottom). (G) Schematic of the dose-optimization strategy for allele-specific *Ldb3* targeting. Three-month-old *Ldb3^Ala165Val/+^* mice received a single intramuscular injection of AAV-shRNAmiR (sh10 or sh16), and TA muscles were analyzed 1 month later. (H) Schematic of the therapeutic disease-reversal paradigm. *Ldb3^Ala165Val/+^* mice were injected intramuscularly with AAV-shRNAmiR (sh10 or sh16) at 5 months of age, and outcomes were evaluated at 1 and 3 months post-injection. All data are presented as mean ± SD of three biological replicates (n = 3) with technical triplicates. Only statistically significant comparisons are shown. *P < 0.05, **P < 0.01 and ***P < 0.001, as determined by Welch’s two-sample t test. si-SC, scrambled siRNA control; NT, no-treatment control. F, G and H created from Biorender.com.

### In-vivo validation of AAV9-shLDB3-mir-30 mediated downregulation of the *Ldb3^A165V^*allele

To selectively knockdown the mutant *Ldb3^A165V^* allele in skeletal muscle, we used an artificial microRNA-based shRNA platform incorporating the miR-30 backbone. shRNAmiR constructs encoding the si10 and si16 sequences validated *in-vitro* were packaged into AAV9 and delivered by i.m. injection to the tibialis anterior (TA) muscle of *Ldb*3*^Ala165Val/+^* mice. Following transduction, shRNAmiR transcripts are processed through endogenous microRNA pathways to generate mature inhibitory RNAs that facilitate degradation of target *Ldb3-A165V* mRNA (Figure 1F). BLAST (Basic Local Alignment Search Tool) analysis confirmed that the shRNA sequences exhibited no significant homology to other murine genes, thereby minimizing the potential for offtarget effects (Table S1). AAV9-shRNA-miR_Scr, containing a shRNA sequence with no known target in mouse mRNA, served as the control.

To determine the effective dose of AAV9_shRNA-miR30, we injected 5×10^11^ or 5×10^12^ AAV/kg body weight into the TA muscle of the three-month-old *Ldb3^Ala165Val/+^* mice. The contralateral TA was injected with a similar dose of AAV9-shRNA-miR_Scr (SC) as control (Figure 1G). One month after the injection of the 5×10^11^ AAV/kg dose of shRNA-miR_10 (sh10) and shRNA-miR_16 (sh16), there was no significant knockdown of the *Ldb3-A165V* mutant allele compared to the WT-*A165A allele* (Mean ± SD: mutant allele, 0.93 ± 0.04 vs. WT allele, 0.96 ± 0.03 for sh10, p = 0.15; mutant allele, 0.86 ± 0.12 vs. WT allele, 1.00 ± 0.10 for sh16, p = 0.39) as measured by an optimized allele-specific RT-qPCR (AS-qPCR) (Figure S1A, B and C). However, at the dose 5×10^12^ vg/kg, a single i.m. injection of sh16 and sh10 led to a significant knockdown of *Ldb3*-*A165V* mutant allele compared to the WT-*A165A allele* Mean ± SD: mutant allele, 0.06 ± 0.01 vs. WT allele, 0.29 ± 0.09 for sh16, p = 0.0002; mutant allele, 0.22 ± 0.06 vs. WT allele, 0.96 ± 0.04 for sh10, p = 0.0002) (Figure S1 D and E) *Ldb3^Ala165Val/+^* mice show progressive muscle weakness accompanied by molecular and histopathological features characteristic of MFM, including a marked decline in peak muscle strength by 6 months of age compared with WT littermates.^10^ To evaluate therapeutic rescue, we administered a single i.m. dose (5×10^12^ vg/kg) of sh10 or sh16 into TA muscle of *Ldb3*^Ala165Val/+^ mice (Figure 1H). We confirmed AAV transduction efficiency across these treatment groups by viral genome quantification and EGFP immunostaining (Figures S1F and S2).

### Intramuscular AAV9-shRNA-miR treatment reverses the MFM disease phenotype in advanced-stage *Ldb3^Ala165Val/+^*mice

Because patients with LDB3-MFM are typically diagnosed after symptom onset, we first evaluated therapeutic efficacy in the advanced (post-pathological) stage of the *Ldb3^Ala165Val/+^* mice. To determine whether knockdown of mutant LDB3 reverses established MFM phenotype, we injected a single 5×10^12^ vg/kg dose of sh10 or sh16 into the TA muscle of five-month-old *Ldb3^Ala165Val/+^* mice, with the contralateral TA muscle receiving a scrambled control. Independent cohorts were analyzed at one (5M-1M) and three months (5M-3M) post-injection to assess histopathological, molecular, and functional outcomes to evaluate therapeutic response.

### Allele-specific silencing of mutant LDB3 following AAV9-shRNA treatment

We observed a significant reduction in the mutant allele expression, with approximately 80% and 90% knockdown in the sh16-treated muscle group 5M-1M (n = 3; p = 0.004) and 5M-3M (n = 6; p = 0.007), respectively, using the AS-qPCR (Figure 2A). Similarly, sh10 (5M-1M) and sh10 (5M-3M) treatment groups showed approximately 90% (p = 0.0001) and 35% (p = 0.001) knockdown of the mutant allele, respectively (n = 3-4 per group) (Figure 2B). In the sh16 treatment groups, the WT-allele was relatively preserved, showing knockdown levels of about 55% and 45% at one- and three-month timepoints, respectively (Figure 2A). Whereas with sh10, the expression of WT-*Ldb3* allele remained largely unaffected, with an approximate knockdown of only 5% observed in the sh10 (5M-1M) and sh10 (5M-3M) groups (Figure 2B). Further, immunoblot analysis revealed a significant reduction in LDB3 protein levels in both sh16 and sh10 treatment groups (antibody specific to mutant protein is unavailable). Specifically, total protein levels decreased by approximately 45% in the sh16 (5M-1M) group [(Mean ± SD: SC (5M-1M), 1.03 ± 0.10 vs. sh16 (5M-1M), 0.56 ± 0.08, p = 0.006)] and over 80% in the sh16 (5M-3M) group [SC (5M-3M), 1.04 ± 0.08 vs. sh16 (5M-3M), 0.16 ± 0.11, p = 0.0008)] (Figure 2C). Similarly, the sh10 treatment groups showed an approximate 40% reduction in LDB3 protein levels for both time points, with mean ± SD values of SC (5M-1M), 1.00 ± 0.02 vs. sh10 (5M-1M), 0.61 ± 0.06 (p = 0.0001) and SC (5M-3M), 1.04 ± 0.01 vs. sh10 (5M-3M), 0.58 ± 0.01 (p = 0.0001) (Figure 2D).

**Figure 2.**
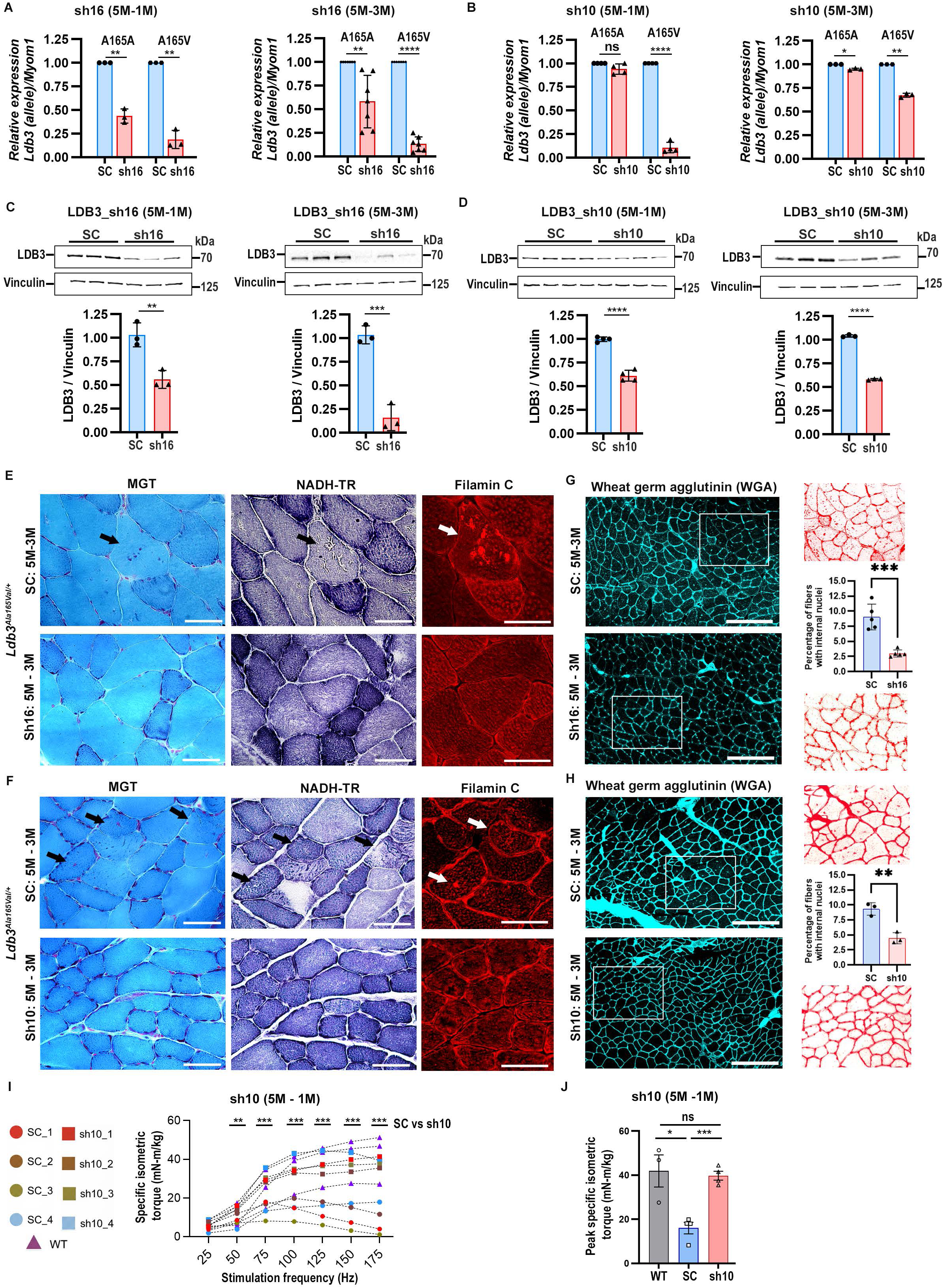
LDB3 knockdown by AAV9-shRNAmiR reverses muscle pathology and functional deficits in advanced-stage *Ldb3^Ala165Val/+^*mice. 5-month-old *Ldb3^Ala165Val/+^*mice received a single intramuscular injection of sh10 or sh16 into the TA muscle; contralateral TA muscles received scramble control (SC). (A and B) AS-qPCR analysis showing relative expression of WT (A165A) and mutant *Ldb3*-A165V transcripts at 1 and 3 months post-injection compared to SC-treated controls in sh16 and sh10 treatment groups. (C and D) Representative immunoblots and densitometric quantification showing total LDB3 protein levels in the sh16 and sh10 treated muscles at 1 and 3 months post-injection compared to SC-treated controls. AS-qPCR and immunoblot data were obtained from 3-7 animals per group and triplicate assays. AS-qPCR data were normalized to *Myom1* using the ΔΔCt method, and immunoblots were normalized to endogenous vinculin protein. (E and F) Adjacent transverse cryosections of frozen TA muscles harvested 3 months posttreatment was stained with modified Gomori trichrome (MGT), NADH-TR, and filamin C. Representative images show sarcoplasmic aggregates and abnormal oxidative enzyme activity (black arrowheads) in SC-treated controls (top panels) compared with sh16- and sh10-treated muscles (bottom panels). Filamin C-positive aggregates are indicated by white arrowheads in SC-treated and absence in sh16 and sh10 treated muscle. (G and H) Representative images of frozen transverse TA muscle cryosections stained with wheat germ agglutinin (WGA; cyan) showing internal nuclei (red dots in magnified insets), together with scatter bar plots quantifying the percentage of fibers containing internal nuclei in sh16- and sh10-treated muscles compared with SC-treated controls. Scale bars: MGT, NADH-TR, and filamin C: 50 µm; WGA: 250 µm. (N = 3-6 TA muscles per group). (I and J) *In-vivo* assessment of muscle contractility in 5-month-old *Ldb3^Ala165Val/+^* mice treated with sh10 for 1 month. (I) Force-frequency relationship curves showing specific isometric torque generated across increasing stimulation frequencies in WT mice, sh10-treated TA muscles, and contralateral SC-treated TA muscles. (J) Quantification of peak specific isometric torque in WT, sh10-treated TA muscles, and contralateral SC-treated TA muscles. In the force-frequency curves, matching colored rectangles and circles denote shRNA-treated TA muscles and the corresponding contralateral SC-treated TA muscles from the same mice, respectively, while purple triangles denote PBS-treated WT TA muscles used as reference controls (n = 3-4 mice per group). Data are presented as mean ± SD (A-F) and mean ± SEM (J). Statistical significance: *P < 0.05, **P < 0.01, ***P < 0.001, and ****P < 0.0001; ns, not significant, as determined by an unpaired, two-tailed Student’s t-test (A, B, I and J) and Welch’s t-test (C and D).

### Reversal of hallmark skeletal muscle pathology of MFM

Microscopic examination of modified Gomori-Trichrome (MGT)-stained TA muscle sections from the Sh10 treated (n = 3) or Sh16 treated (n =6) groups at 5M-3M showed marked reduction or total absence of muscle fibers containing bluish-red granular deposits and rimmed vacuoles, accompanied by normalization of oxidative enzyme activity on nicotinamide adenine dinucleotide–tetrazolium reductase (NADH-TR) staining and reduced accumulation of filamin C on immunostaining in the adjacent serial sections of same muscle fibers. In contrast, as expected, the pathological hallmarks of MFM were readily observed in the scrambled-treated muscle in the opposite hindlimb of the same mice (Figure 2E and F).

Furthermore, treated groups sh16 (5M-3M) and sh10 (5M-3M) showed a marked reduction or absence of myopathic features compared to the contralateral control SC (5M-3M)) group (Figure 2E and F). Notably, compared to the scrambled-treated muscle, the contralateral treated muscle showed a significant decrease in the percentage of fibers with internal nuclei (SC, 5M-3M: 9.03 ± 2.13 vs. sh16, 5M-3M: 3.01 ± 0.59; n = 5; p = 0.0003) and SC, 5M-3M: 9.32 ± 1.04 vs. sh10, 5M-3M: 4.46 ± 0.94; n = 3; p = 0.003), approaching to the levels observed in WT-muscle (Figure 2G and H).^10^

### Restoration of muscle contractile function

In light of the robust suppression of the mutant allele across treatment groups and the pronounced histopathological improvement observed at 5M-3M, we next sought to determine whether mutant LDB3 knockdown translated into measurable functional benefit. To assess early physiological recovery following intervention at an advanced disease stage, *in-vivo* TA muscle contractility was evaluated in a representative cohort treated at five months of age and analyzed one month later (5M-1M) following sh10 treatment.

TA muscle torque was measured by stimulation of the fibular (common peroneal) nerve to assess cumulative isometric force generation at the ankle joint. At one-month post-treatment (5M-1M), TA muscle produced a marked increase in specific isometric torque across stimulation frequencies, generating 4- to 5-fold greater torque from 100 Hz onward compared with contralateral scrambled-treated controls (Figure 2I and S3). Peak specific torque was significantly increased in sh10-treated muscle relative to scramble controls (mean ± SEM: 39.8 ± 2.0 vs. 16.0 ± 2.6; n = 4 per group; p =0.0004) and overlapped with values observed in six-month-old PBS treated WT-TA muscle (41.9 ± 7.3; n = 3), indicating restoration of muscle strength toward the physiological WT range (Figure 2J).

### Reduction of filamin C and CASA chaperone aggregation

Mechanical strain from muscle contraction triggers the unfolding of filamin C, initiating degradation through the CASA pathway, essential for protein homeostasis in skeletal muscle.^19^ In *Ldb3^Ala165Val/+^* mice, sarcoplasmic aggregation of filamin C and its chaperones BCL2-associated athanogene 3 (BAG3) and heat shock protein family A member 8 (HSPA8; HSC70) accumulations were the earliest hallmark of muscle pathology even before LDB3 accumulation.^10^ Because filamin C is important for the disease pathomechanism and is the earliest biomarker of muscle pathology in the *Ldb3^Ala165Val/+^* knock in mice, we examined distribution of filamin C, BAG3, and HSPA8 proteins by immunofluorescence (IF) staining on longitudinal sections of paraformaldehyde (PFA)-perfused TA muscle in the control (SC) and shRNA treatment groups in advanced-stage mice (n = 3-6 per group) (Figure 3A and B). Quantitative analysis showed a significant reduction in the percentage of muscle fibers with filamin C accumulations in the sh10 or sh16 treated TA muscle compared to the contralateral scrambled-treated muscle (n = 3 per group).

**Figure 3.**
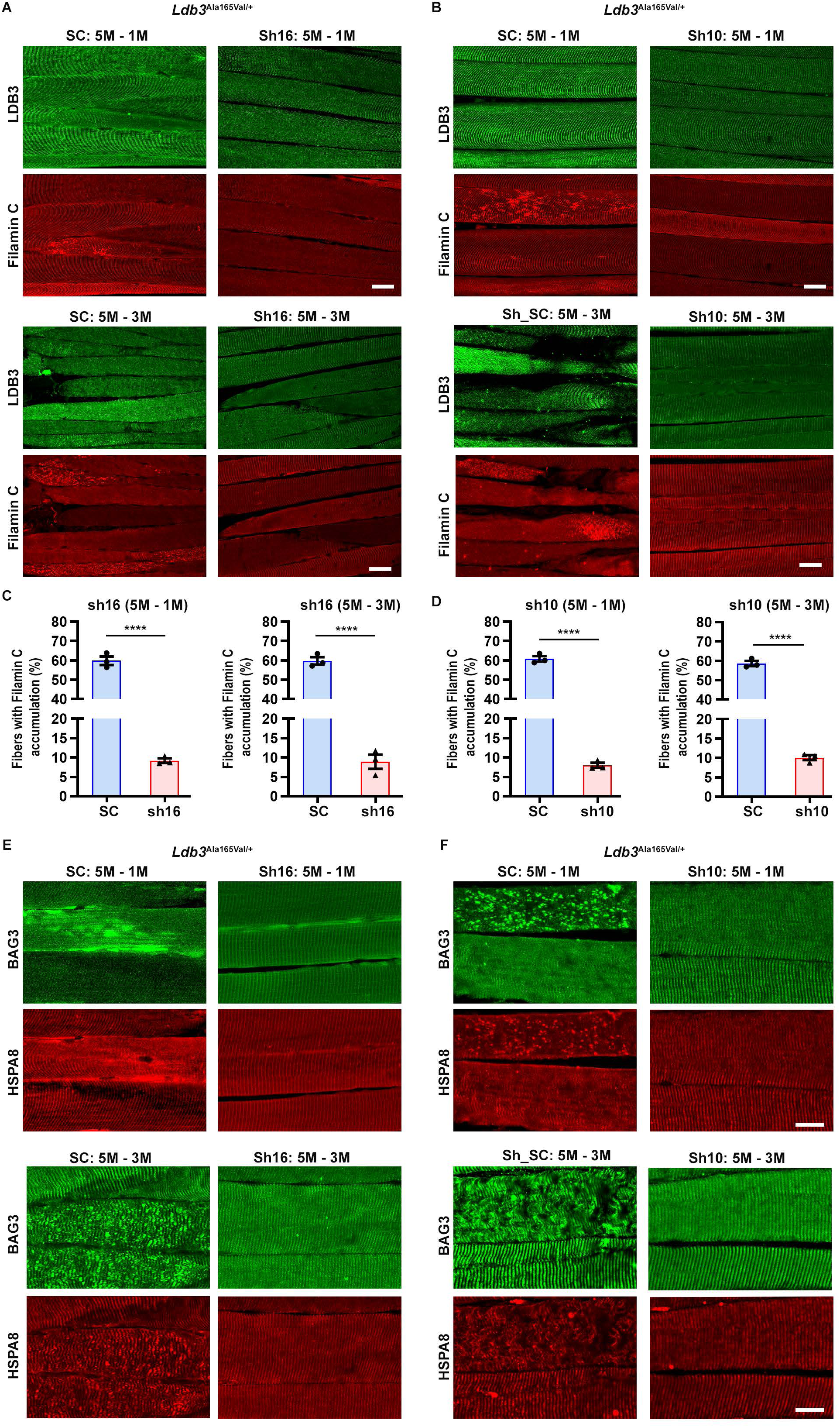
AAV9-shRNAmiR treatment led to clearance of myofibrillar and CASA-associated protein aggregates and restoration of Z-disc architecture in advanced-stage *Ldb3^Ala165Val/+^* mice. Representative IF images of LDB3 and filamin C in PFA perfused longitudinal TA muscle sections from advanced-stage, (5-month-old) *Ldb3^Ala165Val/+^* mice treated with sh-SC, sh16, or sh10 (A and B) for one month (upper panel) or three months (bottom panel) (n = 4-6 per group). (C and D) Quantification of muscle fibers containing sarcoplasmic filamin C aggregates in sh-SC control and shRNA-treated groups. Bar scatter plots represent the percentage of fibers exhibiting filamin C aggregates, quantified from at least five randomly selected areas (> 50 fibers per muscle) from immunostained TA sections (n = 3 per group). (E and F) Representative IF images of BAG3 and HSPA8 in longitudinal TA cryosections from advanced-stage, (5-month-old) *Ldb3^Ala165Val/+^*mice treated with sh-SC, sh16, or sh10 for 1 month (upper panels) or 3 months (lower panels) (n = 4-6 per group). Data are expressed as mean ± SEM, with statistical significance denoted by ****P < 0.0001, determined using an unpaired, two-tailed Student’s t-test. Scale bars represent 50 µm (A and B) and 20 µm (E and F).

There was approximately 85% decrease in the filamin C aggregation at 1 month and 3 months after sh16 treatment [SC (5M-1M), mean ± SEM: 59.84 ± 3.75 vs. sh16 (5M-1M), 9.17 ± 1.02 (p = 0.0001) and SC (5M-3M), 59.79 ± 3.30 vs. sh16 (5M-3M), 8.91 ± 3.17 (p = 0.0001)] (Figure 3C and S4). Similarly, the sh10 treatment led to an 88% and 83% reduction in fibers with filamin C accumulation in 5M-1M and 5M-3M group, respectively (5M-1M), 60.85 ± 2.47 vs. sh10 (5M-1M), 9.98 ± 1.1 (p = 0.0001) and SC (5M-3M), 58.68 ± 2.31 vs. sh10 (5M-3M), 10.07 ± 1.14 (p = 0.0001) (Figure 3D and S4). The reduction in filamin C aggregation was accompanied by reduction in the accumulation of its co-chaperone proteins, BAG3 and HSPA8, in both sh16 and sh10 treated muscle compared to contralateral scrambled-treated TA muscle (Figure 3E and F).

### Restoration of PKC_α_ following mutant LDB3 knockdown

PKCα, the principal conventional PKC isoform in skeletal muscle, localizes to Z-disc through its association with LDB3, where it phosphorylates LDB3 and other force-responsive Z-disc associated proteins.^20^ We previously demonstrated that PKCα protein levels are reduced by approximately 50% in 8-month-old *Ldb3^Ala165Val/+^* mice compared with WT littermates.^10^ To determine whether mutant LDB3 knockdown restores PKCα expression, we performed immunoblot analysis of TA muscles in which the contralateral limb of the same mouse received shRNA or SC treatment, along with TA muscle from age-matched PBS-treated WT littermates (n = 3). Both treatment groups showed a significant increase in PKCα protein levels in the treated TA muscle compared to SC-treated contralateral TA muscle, achieving expression levels comparable to age matched WT muscle. The sh16 (5M-1M) treatment group showed PKCα levels (mean ± SD) of 0.97 ± 0.06 in WT, 0.56 ± 0.06 in SC, and 1.00 ± 0.04 in sh16 (SC:sh16; p = 0.0009), while the sh16 (5M-3M) group showed values of 1.03 ± 0.02 in WT, 0.55 ± 0.02 in SC, and 0.98 ± 0.05 in sh16 (SC:sh16; p = 0.0002) (Figure 4A). Similarly, sh10 treatment significantly increased PKCα levels. In the sh10 (5M-1M) group, levels measured were 1.04 ± 0.02 for WT, 0.58 ± 0.05 for SC, and 1.02 ± 0.01 for sh10 (p = 0.0001). For the sh10 (5M-3M) group, values were 0.98 ± 0.03 for WT, 0.59 ± 0.04 for SC, and 0.85 ± 0.01 for sh10 (SC:sh10; p = 0.0005) (Figure 4B). Collectively, these findings demonstrate that mutant LDB3 silencing restores PKCα protein levels toward the physiological WT range in shRNA treated skeletal muscle.

**Figure 4.**
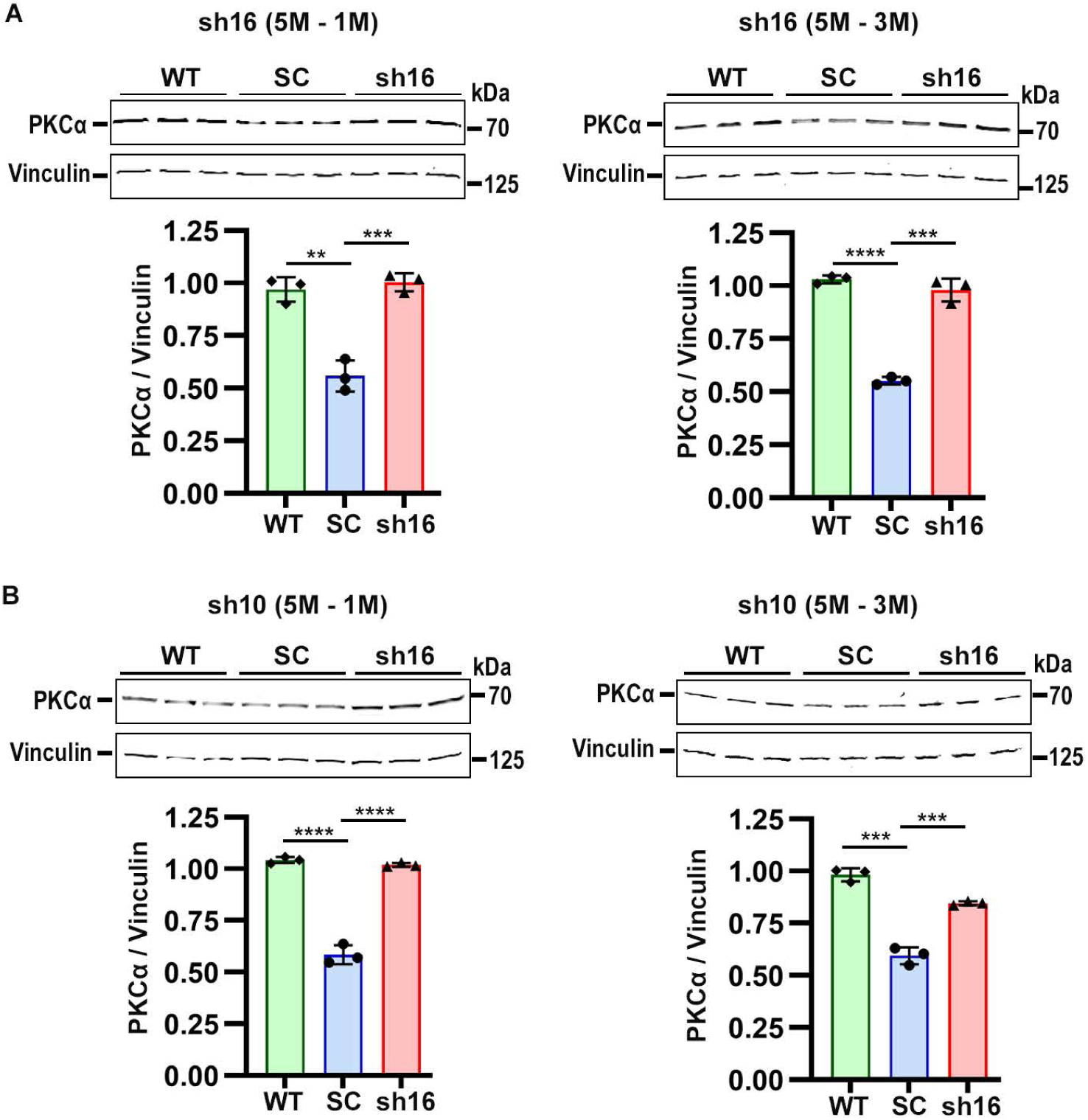
AAV9-shRNAmiR treatment restores PKC*α* levels towards wild-type levels in advanced-stage *Ldb3^Ala165Val/+^* mice. Representative immunoblots showing PKCα protein levels in frozen TA muscle from 5-monthold *Ldb3^Ala165Val/+^* mice treated with AAV9-shRNAmiR constructs and age-matched PBS treated WT controls. (A and B) PKCα protein levels following sh16 (A) or sh10 (B) treatment at 1 and 3 months post-injection. Bar-scatter plots below the blots show densitometric quantification of PKCα normalized to endogenous vinculin (n = 3 per group and triplicate assays). *Ldb3^Ala165Val/+^*mice were administered either scramble control (SC) or shRNAmiR targeting sh16 or sh10. Data are presented as mean ± SD, with statistical significance levels of **P < 0.01, ***P < 0.001, ****P < 0.0001, calculated using a two-tailed unpaired Student’s t-test.

### Early intramuscular AAV9-shRNA-miR treatment prevents MFM-associated proteostatic and signaling abnormalities in *Ldb3^Ala165Val/+^*mice

Early therapeutic interventions offer the potential to prevent or significantly delay the onset of debilitating diseases, thereby improving patient outcomes and quality of life. At 4 months of age, muscle histology in the *Ldb3^Ala165Val/+^* mice appear similar to that in the WT mice on transverse muscle sections. Longitudinal sections of the TA muscle show normal distribution of LDB3 at Z-disc; however, there are visible accumulations of filamin C, BAG3, and HSPA8 at the Z-discs spanning multiple sarcomeres along the length of muscle fibers.^10^ Therefore, we treated 3-month-old *Ldb3^Ala165Val/+^* mice to investigate the potential of early therapy to prevent disease onset and progression.

We administered a single intramuscular dose (5 × 10¹² vg/kg body weight) of either sh10 or sh16 into the TA muscle of *Ldb3^Ala165Val/+^* mice at 3 months of age, with separate groups analyzed at 1 month (3M-3M) and 3 months (3M-3M) post-injection (Figure 5A). We confirmed AAV transduction efficiency in all treatment groups by quantifying viral genomes and assessing EGFP fluorescence (Figures S1E and S2).

**Figure 5.**
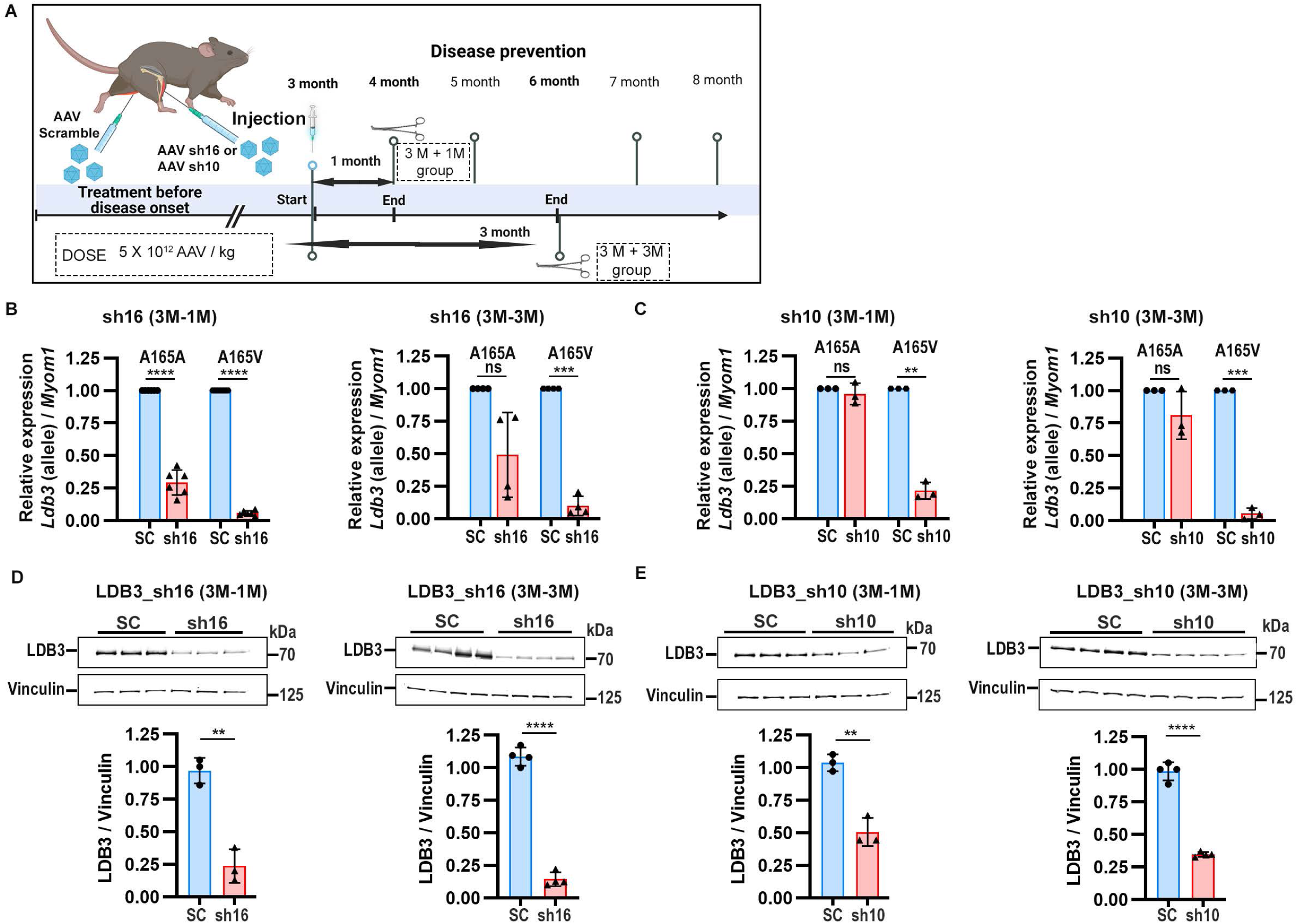
Efficient *Ldb3* knockdown in early-stage (3-month-old) *Ldb3^Ala165Val/+^* mice following AAV9-shRNAmiR treatment. Three-month-old *Ldb3^Ala165Val/+^*mice received a single intramuscular injection of sh10 or sh16 into the TA muscle; contralateral TA muscles received scramble control (SC). (A) Schematic of the early-intervention (disease-prevention) paradigm. ***Ldb3^Ala165Val/+^*** mice were injected intramuscularly with AAV-shRNAmiR (sh10 or sh16) at 3 months of age, and outcomes were evaluated at 1 and 3 months post-injection (created from Biorender.com). (B and C) AS-qPCR analysis showing relative expression of WT (A165A) and mutant *Ldb3*-A165V transcripts at 1 and 3 months post-injection compared to scramble-treated controls (SC) in sh16 and sh10 treatment groups. (D and E) Representative immunoblots and densitometric quantification showing total LDB3 protein levels in the sh16 and sh10 treated muscles at 1 and 3 months post-injection compared to SC-treated controls. AS-qPCR and immunoblot data were obtained from 3-6 animals per group and triplicate assays. AS-qPCR data were normalized to endogenous *Myom1* using the ΔΔCt method, and immunoblots were normalized to endogenous vinculin protein. Data are presented as mean ± SD, with statistical significance defined as **P < 0.01, ***P < 0.001, and ****P < 0.0001; ns, not significant, as determined by unpaired, two-tailed Student’s t-test (B and C) and Welch’s t test (D and E).

### Allele-specific silencing of mutant LDB3

The sh16 treated muscle showed approximately 94% and 90% knockdown of the mutant *Ldb3 A165V* allele at one-month and 3-months after injection, respectively (n = 4-6; p ≤ 0.0001). The sh10 treated muscle showed 78% and 95% knockdown of the mutant one month and 3 months after injection, respectively (n = 3; p < 0.002). We observed 71% and 51% knockdown of WT allele in the sh16 treated muscle at one-month and 3-months, respectively. Whereas, the WT allele remained largely unaffected by sh10 treatment, with 4% and 19% knockdown at one month and 3-months after injection, respectively (Figure 5B and C).

Immunoblot analysis confirmed LDB3 knockdown, demonstrating significant reductions in total LDB3 protein levels in both sh16 and sh10 treatment groups. The sh16 treatment led to approximately 75% and 85% reduction in the total LDB3 protein at one month (mean ± SD: SC, 0.97 ± 0.10 vs. sh16, 0.24 ± 0.13, p = 0.001) and at 3 months (SC, 1.09 ± 0.07 vs. sh16, 0.14 ± 0.05, p < 0.0001), respectively (Figure 5D). Treatment with sh10 led to approximately 50% and 65% reductions in total LDB3 at one month (SC, 1.04 ± 0.06 vs. sh10, 0.51 ± 0.11, p = 0.001) and at 3 months (SC, 0.98 ± 0.07 vs. sh10, 0.35 ± 0.02, p < 0.0001), respectively (Figure 5E).

### Prevention of filamin C and CASA-associated chaperone aggregation

Next, we performed quantitative IF analysis to examine the distribution of filamin C and chaperone proteins in the muscle fibers. Note that aggregation of filamin C co-localized to its chaperones BAG3 and HSPA8 were the earliest hallmark pathological changes observed from 4-months of age the *Ldb3^Ala165Val/+^* knock-in mice, when there were no apparent changes in the LDB3 distribution by immunofluorescence nor in muscle histology.^10^ We found a pronounced reduction or absence of filamin C accumulation in the sh10 or sh16 treated TA muscle as compared to the contralateral scrambled-treated TA muscle. At one month after injection, the sh16-treated mice showed ∼92% decrease in fibers with filamin C accumulations. In the SC group, 56.87 ± 2.76% of fibers contained filamin C accumulations, whereas in the sh16 treated group only 3.86 ± 2.17% fibers contained filamin C accumulations (p = 0.0001). Similarly, at 3 months post-injection the SC group showed 58.79 ± 2.77% of fibers with filamin C accumulations, compared to only 4.94 ± 1.44% fibers with filamin C accumulations in the contralateral sh16 treated TA muscle (p < 0.0001) (Figure 6A, B and S5).

**Figure 6.**
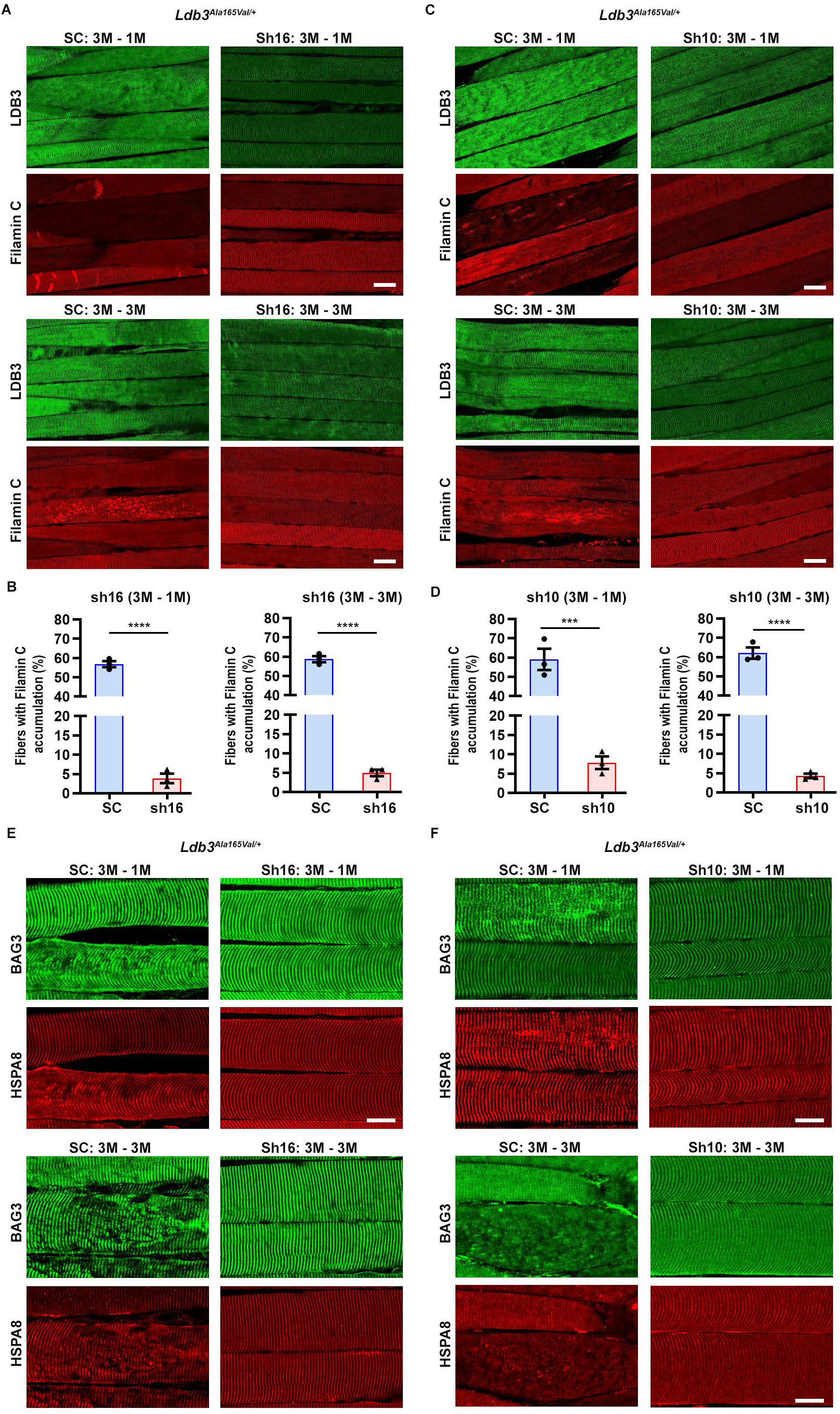
Early intervention with AAV9-shRNAmiR prevents the development of myofibrillar and CASA-associated protein aggregates in *Ldb3^Ala165Val/+^* mice. (A and C) Representative IF images of LDB3 and filamin C in PFA perfused longitudinal TA muscle sections from early-stage (3-month-old) *Ldb3^Ala165Val/+^* mice treated with sh-SC, sh16 (A), or sh10 (C) for one month (upper panel) or three months (lower panel) (n = 4-6 per group). (B and D) Quantification of muscle fibers containing sarcoplasmic filamin C aggregates in sh-SC and shRNA-treated groups corresponding to panels (A) and (C). Bar-scatter plots represent the percentage of fibers exhibiting filamin C aggregates, quantified from at least five randomly selected fields (>50 fibers per muscle) per section (n = 3 per group). (E and F) Representative IF images of BAG3 and HSPA8 in longitudinal TA cryosections from early-stage, (3-month-old) *Ldb3^Ala165Val/+^* mice treated with sh-SC, sh16, or sh10 for 1 month (upper panels) or 3 months (lower panels) (n = 4-6 per group). Data are expressed as mean ± SEM, with statistical significance denoted by **P < 0.01, determined using an unpaired, two-tailed Student’s t-test. Scale bars represent 50 µm (A and C) and 20 µm (E and F).

We observed a similarly robust reduction in filamin C aggregation in sh10-treated muscle compared with contralateral scramble-treated controls, with 87% and 93% decreases in fibers containing filamin C accumulations at 3M-1M and 3M-3M, respectively. At one-month postinjection, only 7.80 ± 2.86% of fibers showed filamin C accumulations, compared with 59.10 ± 9.57% in SC-treated muscle (p = 0.0009). At 3 months after injection, filamin C accumulation was limited to 4.27 ± 1.14% of fibers following sh10 administration, versus 62.12 ± 5.04% in SC controls (p < 0.0001) (Figure 6 C, D and S5). Additionally, IF staining indicated a substantial absence of the accumulation of co-chaperones BAG3 and HSPA8 in both groups sh16 and sh10 treated TA muscle as compared to scrambled-treated TA muscle (Figure 6E and F).

### Retention of normal muscle contractile function

*Ldb3^Ala165Val/+^* mice exhibit progressive muscle weakness, with reduced grip strength beginning at 3 months and significant contractile impairment evident by 6 months of age.^10^ To investigate the treatment efficacy of early-stage interventions, we conducted *in-vivo* muscle contractility assays on the TA muscle from the sh10 treated muscle at one month after injection, and both sh10- and sh16-treated muscle at 4 months after injection, alongside controls (SC) and the age matched PBS treated WT mice for reference.

We found a significant increase in specific isometric torque in the shRNA-treated TA muscle one month after injection compared with the contralateral scrambled-treated muscle across stimulation frequencies. Torque increased approximately 2.5- to 20-fold from 100 Hz onward relative to scrambled-treated controls, reaching levels comparable to those observed in WT TA muscle at corresponding frequencies (Figures 7A and S3). Peak specific isometric torque was significantly higher in sh10-treated muscle compared with SC controls (mean ± SEM): SC, 11.4 ± 1.6; sh10, 31.5 ± 3.6; n = 4 per group; p = 0.002) (Figure 7A). Notably, peak torque values overlapped with those observed in six-month-old WT-TA muscle (28.6 ± 3.4; n = 3), indicating preservation of physiological muscle contractile performance following early RNAi intervention.

**Figure 7.**
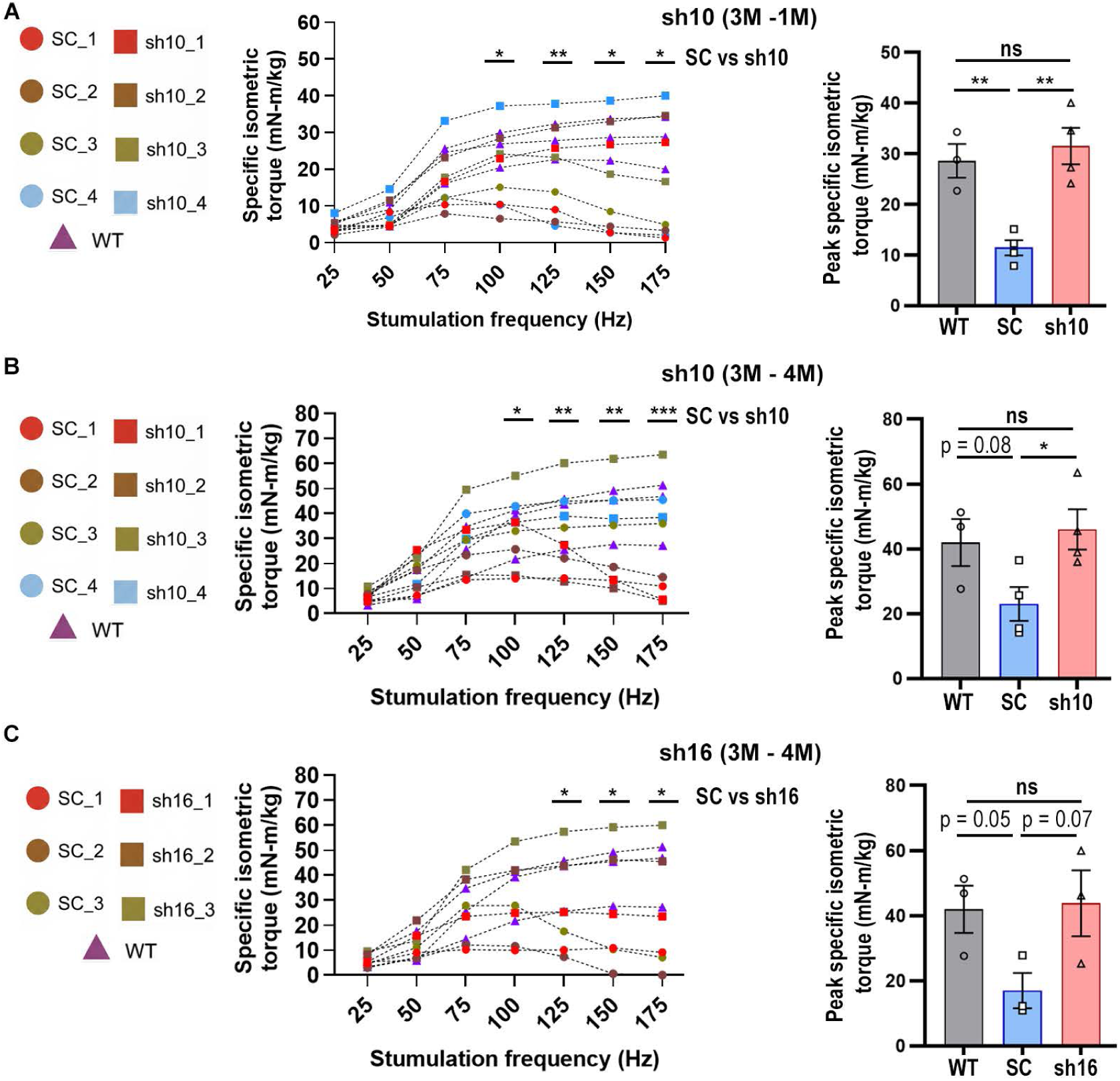
Early AAV9-shRNAmiR treatment preserves muscle contractility toward wild-type levels in *Ldb3^Ala165Val/+^*mice. (A, B and C) In vivo force-frequency relationship curves (left panels) showing specific isometric torque generated across increasing stimulation frequencies in TA muscles from PBS-treated WT mice, shRNA-treated mice, and contralateral SC-treated muscles. shRNA was administered at 3 months of age, with functional assessment performed at 1 month (A; 3M-1M) or 4 months (B and C; 3M-4M) post-injection. Panels represent sh10 treatment (A, B) and sh16 treatment (C). Right panels show corresponding quantification of peak specific isometric torque presented as bar-scatter plots. Early shRNA treatment maintained contractile performance toward wild-type levels and prevented the decline observed in SC-treated muscles. In the force-frequency curves, matching colored rectangles and circles denote shRNA-treated TA muscles and the corresponding contralateral SC-treated TA muscles from the same mice, respectively, while purple triangles denote PBS-treated WT TA muscles used as reference controls (n = 3-4 mice per group). Data are presented as mean ± SEM (A-C). ns: not significant. Six-month-old WT-TA muscle used for the contractility assays are shared between sh10 (5M-1M; Figure 21, J), sh10 and sh16 (3M-4M; panels B and C) and served as references due to their overlapping age range. Statistical significance: *P < 0.05, **P < 0.01 and, ***P < 0.001, determined by an unpaired, two-tailed Student’s t-test.

Notably, in both longer-term treatment groups (3M-4M), both sh10- and sh16-treated TA muscle continued to demonstrate enhanced contractile performance relative to their respective SC controls. Force-frequency analysis showed significantly greater specific isometric torque in both treatment groups from 100 Hz and 125 Hz onwards for sh10 and sh16 respectively compared with SC controls (Figure 7B, C and S3). Across increasing stimulation frequencies, specific isometric torque increased approximately 2- to 20-fold in sh10-treated mice and 2.5- to 18-fold in sh16-treated mice relative to SC controls.

Consistent with these findings, peak specific isometric torque 4 months post-injection remained increased in sh10-treated muscle (mean ± SEM: SC, 23.0 ± 5.2; sh10, 46.0 ± 6.2; n = 4 per group), reaching statistical significance (p = 0.02) (Figure 7B and C). In sh16-treated muscle, peak torque was similarly increased (mean ± SEM: SC, 27.0 ± 6.8; sh16, 43.8 ± 10.1; n = 3 per group), demonstrating a strong trend toward significance (p = 0.07) (Figure 7B). In both treatment groups, peak torque values met or exceeded those observed in six-month-old WT-TA muscle. Overall, increased muscle force production was evident at 1 month and 4 months following a single intramuscular administration of shRNA. Collectively, these findings demonstrate that early AAV9-shRNA-miR intervention effectively prevents contractile dysfunction and preserves near-physiological muscle performance over the four-month study period.

### Early Ldb3-RNAi treatment restores PKC_α_ level and reprograms skeletal muscle phosphoproteomic signaling

LDB3 interacts directly with and is phosphorylated by PKCα, a predominant kinase accounting for approximately 97% of conventional PKCs in skeletal muscle.^20^ LDB3 acts as a signaling scaffold in striated muscle tethering PKCα to the Z-disc, where the kinase phosphorylates proteins that play important role in muscle integrity and function. For example, *in-vitro* studies demonstrate influence of PKCα activity on muscle structure and dynamics by regulating filamin C.^23^ Phosphorylation of filamin C by PKCα inhibits filamin C cleavage and stabilizes interactions of filamin C with its ligands at the Z-disc. Previous work by Pathak et al. (2021) demonstrated a significant downregulation of PKCα (approximately by 50%) as early as 4 months of age in *Ldb3^Ala165Val/+^*mice compared to WT-littermates.^10^ Building upon these findings, subsequent analyses focused on evaluating the impact of early treatments on PKCα protein levels using immunoblot assays. Age-matched, PBS treated WT-TA muscle samples were included as reference to assess the extent of PKCα recovery in treated mice.

Specifically, in the early AAV9-shRNA treatment experiments (i.e., AAV9-shRNA injection at 3 months of age before visible phenotype), we found PKCα levels (mean ± SD) of 0.98 ± 0.07 in the sh16 treated TA muscle compared to 0.61 ± 0.03 in the SC and 1.00 ± 0.04 in WT PBS-treated mice, (n = 3 per group; p = 0.001), respectively. Similarly, at 3 months post injection, PKCα levels (mean ± SD) were 0.97 ± 0.12 in sh16 treated TA muscle, 0.55 ± 0.03 for SC and 1.02 ± 0.01 for WT, and (n = 3 per group; p = 0.004) (Figure 8A).

**Figure 8.**
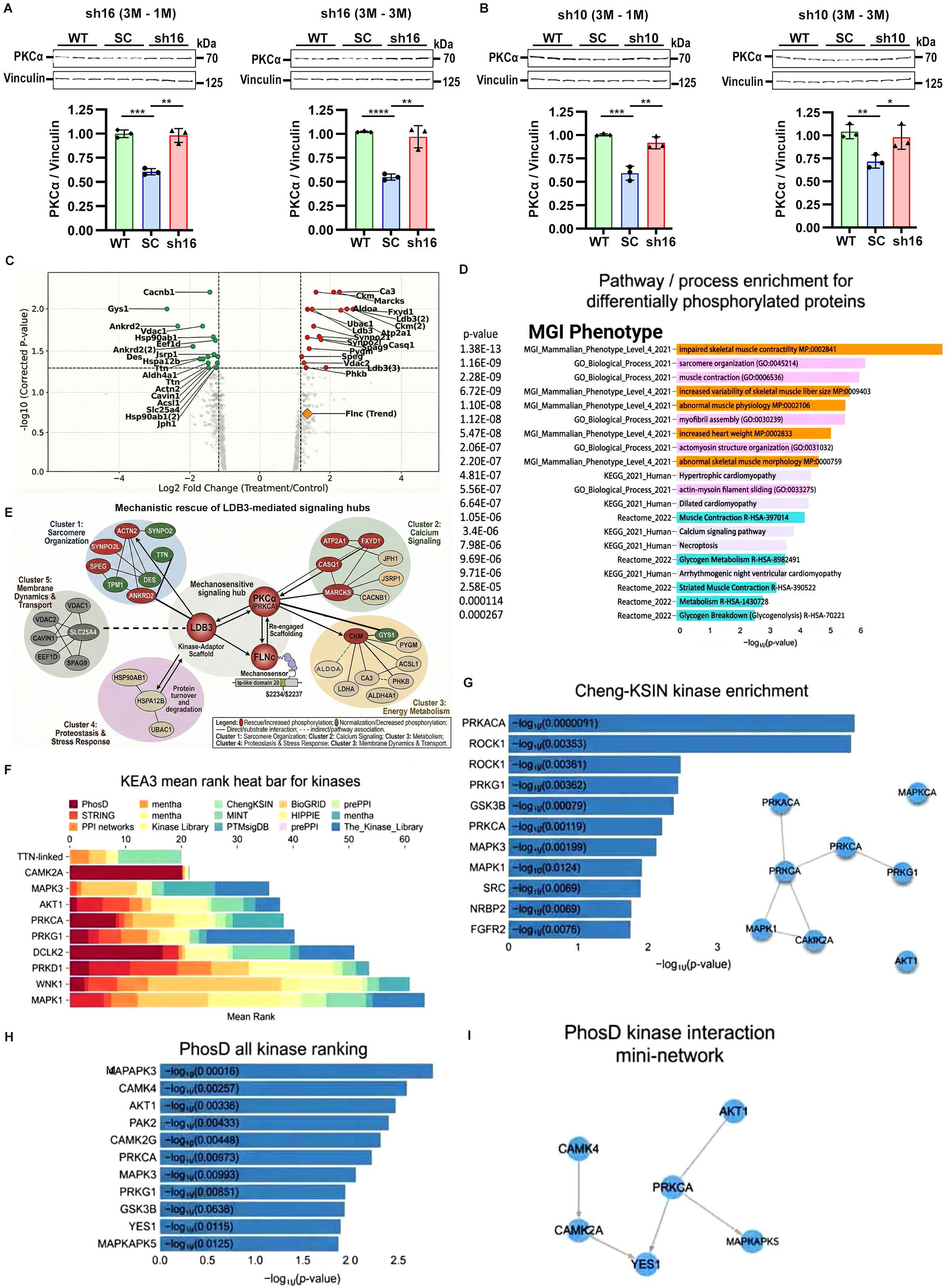
Early AAV9-shRNAmiR treatment normalizes PKCα levels and restores PKC*α*-responsive signaling networks in *Ldb3^Ala165Val/+^* mice. Representative immunoblots showing PKCα protein levels in frozen TA muscle from 3-monthold *Ldb3^Ala165Val/+^* mice treated with AAV9-shRNAmiR constructs and age-matched PBS treated WT controls. (A and B) PKCα expression following sh16 (A) or sh10 (B) treatment at 1 and 3 months post-injection. Bar-scatter plots below the blots show densitometric quantification of PKCα normalized to endogenous vinculin (n = 3 per group and triplicate assays). *Ldb3^Ala165Val/+^* mice were administered either scramble control (SC) or shRNAmiR targeting sh16 or sh10. Data are presented as mean ± SD, with statistical significance levels of *P < 0.05, **P < 0.01, ***P < 0.001, ****P < 0.0001, calculated using a two-tailed unpaired Student’s t-test. Phosphoproteomic profiling was performed to compare shRNA-treated and SC-treated TA muscles (n = 3 per group). (C) Volcano plot of differentially phosphorylated proteins in sh16-treated versus SC-treated TA muscle. Each point represents a phosphoprotein; red points indicate significantly upregulated phosphosites and green points indicate significantly downregulated phosphosites (adjusted P < 0.05), whereas gray points denote non-significant changes. The x axis represents log□ fold change (treatment/control), and the y axis represents −log□□ adjusted P values. A total of 35 phosphoproteins met significance criteria (adjusted P < 0.05). (D) Pathway and process enrichment analysis of differentially phosphorylated proteins. Significantly enriched MGI Phenotype and Gene Ontology (GO) Biological Process terms are shown. Bars represent−log□□ (P value); unadjusted P values are displayed. (E) Protein-protein interaction network constructed from significantly altered phosphoproteins. Nodes represent phosphoproteins; edges indicate curated interactions. PKCα is positioned within the network to visualize connectivity among structural and signaling modules. (F) KEA3 kinase enrichment analysis. Mean rank heat bar showing top-ranked upstream kinases inferred from the phosphoproteomic dataset across multiple kinase-substrate libraries (PhosD, STRING, and Kinase Library). Color scale reflects relative ranking within each database. (G) Cheng-KSIN kinase enrichment analysis. Bar plot and corresponding network visualization of kinases enriched based on motif-derived kinase–substrate interactions. Enrichment is represented as −log(P value). (H) PhosD all-kinase ranking. Top kinases ranked using kinase-substrate co-phosphorylation associations derived from the PhosD library. (I) PhosD kinase interaction mini-network illustrating inferred relationships among PRKCA (PKCα) and associated signaling nodes.

The sh10 treatment likewise resulted in a robust recovery of PKCα protein levels, towards those observed in WT muscle. One month after injection, PKCα levels (mean ± SD) reached 0.92 ± 0.06 in sh10-treated TA muscle, compared with 0.59 ± 0.07 in SC and 1.00 ± 0.01 in WT mice (n = 3 per group; p = 0.005). By three months post-injection, PKCα levels remained elevated in sh10-treated muscle (0.98 ± 0.13) relative to SC (0.72 ± 0.07) and were comparable to WT (1.04 ± 0.08) (n = 3 per group; p = 0.04) (Figure 8B). Thus, consistent with findings in the later-treated advanced-stage disease mice, both sh10 and sh16 treatment led to a robust and statistically significant increase in PKCα protein levels in the skeletal muscle of the *Ldb3^Ala165Val/+^* mice, bringing these levels close to those seen in the WT mice muscle.

### Restoration of PKC_α_-responsive mechanosensing signaling networks

In the context of established therapeutic restoration of PKCα levels in the sh16-treated *Ldb3^Ala165Val/+^* mice, we performed paired phosphoproteomic analyses comparing untreated (SC-treated control) muscle, representing as the pathological reference state, with sh16-treated contralateral muscle to define treatment driven signaling changes. These studies were performed one month after treatment (3M-1M; 4 months of age), when control (SC-treated) mutant mice muscle exhibited early filamin C and CASA accumulation without overt structural degeneration, thereby allowing us to capture primary signaling alterations while minimizing secondary remodeling effects. Using Tandem Mass Tag (TMT) based quantitative proteomics approach, differential phosphorylation was assessed at the phosphosite level using Welch’s two-sample t test with multiple-testing correction. Volcano plot analysis integrating log fold change and −log adjusted P-values identified 35 phosphoproteins with significant phosphorylation differences (adjusted p < 0.05) between scrambled controland LDB3-sh16-treated TA muscle of the same mice (Figure 8C). These proteins spanned structural Z-disc components, contractile regulators, calcium-handling proteins, and metabolic enzymes, indicating coordinated remodeling of mechanosensitive and sarcomeric signaling pathways. Treatment responsiveness was defined at the protein level by the presence of ≥1 significant phosphosite per protein; individual phosphosites were retained and reported separately (Table S2). Notably, quantitative phosphoproteomic analysis identified a treatment-responsive phosphorylation signal within the mechanosensitive Ig like domain 20 region of FLNc following sh16 treatment. This corresponds to approximately 37% increase in the p-FLNC/total FLNC ratio relative to scrambled-treated muscle. While Welch’s testing yielded a borderline p value (0.0598), an equal-variance Student’s t-test reached statistical significance (p = 0.0457) (Figure S1G). Although phospho-specific anti-bodies for specific FLNc serine residues are not commercially available, the increased phosphopeptide intensities, together with restored PKCα protein abundance, support site-specific rescue of FLNc phosphorylation.

In parallel, phosphorylation of LDB3 increased at S179 (2.77-fold, p = 0.0001) and S98 (1.90-fold, p = 0.0029; Table S3), consistent with re-engagement of the PKCα-scaffolding function of LDB3.^21,22^ Together with the findings of restored PKCα levels (Figure 8A and B) and established roles for PKCα-dependent phosphorylation in protecting FLNC from calpain cleavage and CASA-linked turnover,^23^ these data demonstrate that allele-specific RNAi reconstitutes the core LDB3-PKCα-FLNc phosphorylation axis, which leads to the recovery of MFM pathology in mice.

### Ldb3-RNAi normalizes PKC_α_-responsive structural, mechanosensory, and metabolic signaling networks

Beyond FLNc, LDB3-sh16 increased phosphorylation across multiple PKCα-linked nodes curated in Table S3, including canonical PKCα substrates MARCKS (2.12-fold, p = 0.00004) and FXYD1 (2.49-fold, p = 0.0002), as well as metabolic and Ca^2+^-handling proteins such as CKM (S164/T166: 2.68-fold, p = 0.0001) and ATP2A1 (2.32-fold, p = 0.0002). Conversely, several tension/structure-associated proteins showed decreased phosphorylation, including TTN T299/S301 (−1.64-fold, p = 0.0003) and DES S437 (−1.29-fold, p = 0.00008; Table S3), suggesting normalization of pathological strain signaling.^24,25^

To contextualize these phosphosite-level changes, we used Enrichr-KG (a gene set enrichment analysis tool)^26^ to analyze the set of phosphoproteins showing significant phosphorylation differences. Enrichment highlighted core muscle systems, including muscle contraction (p = 2.28E−09) and sarcomere organization (p = 1.16E−09), as well as calcium signaling (p = 3.40E-06) and metabolic pathways (p = 1.14E−04; Figure 8D and Table S4). To integrate the phosphoproteomic alterations within a signaling framework, we constructed a kinase-anchored interaction network from the 35 significant phosphoproteins, placing PKCα as a central node and revealing interconnected modules converging on an LDB3-PKCα-FLNc axis (Figure 8E). A structural sarcomeric cluster included Z-disc components such as ACTN2, TTN, ANKRD2, and SPEG, linked to FLNc and LDB3. A calcium-handling module comprising ATP2A1, CASQ1, and JPH1 aligned with excitation-contraction coupling pathways. A metabolic cluster containing CKM, GYS1, ALDOA, and LDHA connected energy regulation to structural remodeling. A protein quality-control module, including HSP90AB1, HSPA12B, and UBAC1, reflected phosphorylation changes associated with turnover and stress adaptation. In addition, signaling-associated nodes including MARCKS, FXYD1, SPAG9, and PHKB, bridged structural and metabolic modules. Collectively, these findings indicate that restoration of PKCα signaling through LDB3-RNAi reconstitutes the LDB3-PKCα-FLNC mechanosensing hub and promotes coordinated phosphorylation remodeling across structural, calcium, proteostatic, and metabolic networks in skeletal muscle.

### Kinase-level network analysis indicates reactivation of PKC_α_ signaling following LDB3-shRNA treatment

To gain insight into upstream kinase activities that may underline the phosphorylation changes observed in our dataset, we performed kinase enrichment analysis using KEA3.^27^ Candidate regulatory kinases were inferred using an integrated platform that combines motif-based kinase– substrate predictions (Cheng-KSIN) with co-phosphorylation associations (PhosD) derived from curated phosphoproteomic and protein interaction databases. Although KEA3 contains limited representation of muscle-restricted and sarcomeric proteins, PKCα consistently emerged among the highest-ranked kinases across all aggregated scoring methods (STRING, PhosDAll, Kinase Library; Figure 8F). Network mapping further showed that PKCα clustered within a signaling module containing cAMP-dependent protein kinase catalytic subunit alpha (PKACα),^28^ protein kinase cGMP-dependent 1 (cGKI), calcium/calmodulin-dependent protein kinase type II subunit alpha (CAMK2αA),^29^ mitogen-activated protein kinase 3 ( MAPK3; also called extracellular signal-regulated kinase 1 [ERK-1]),^30^ and RAC-alpha serine/threonine-protein kinase (also called AKT protein kinase B alpha [PKBα ])^31,32^ (Figure 8F).

Consistent with the phosphosite-level findings, PKCα showed high mean ranking across KEA3 libraries and significant enrichment within the Cheng-KSIN and PhosD analyses (Figure 8G and 8H). Network mapping further positioned PKCα within a central kinase interaction module (Figure 8I). Together, these analyses identify PKCα as a prominently enriched upstream kinase associated with the phosphorylation changes observed following shRNA treatment, consistent with the TMT-based profiling.

## Discussion

Here, we demonstrate that allele-directed RNA interference targeting the Ldb3 p.Ala165Val mutation unequivocally confers significant benefit *in-vivo*. In a *Ldb3^Ala165Val/+^* knock-in mouse model that faithfully recapitulates the molecular and functional features of the human condition, a single intramuscular administration produced benefit in two distinct therapeutic paradigms; early intervention prior to overt pathology prevented the emergence of structural and functional abnormalities, whereas treatment initiated after disease establishment reversed existing pathology. Structural, molecular, and muscle contractile improvements were detectable as early as one month after treatment and remained evident through four months post-intervention, at which time muscle force production remained comparable to wild-type physiological levels. These findings establish that knockdown of mutant LDB3 expression is sufficient both to halt disease progression and to durably reconstitute disrupted Z-disc mechanosignaling in post-mitotic skeletal muscle. Although male and female mice were not equally represented, prior studies report comparable disease phenotypes across sexes in this model,^10^ and the contralateral treatment design allowed each mouse to serve as its own control, minimizing potential sex-related variability.

Our siRNA design and validation strategy identified candidate sequences capable of preferentially targeting the mutant *Ldb3* transcript. The si10 exhibited strong allelic discrimination, whereas si16 achieved more robust suppression of the mutant allele with moderate selectivity relative to the WT-allele. When incorporated into a microRNA-adapted shRNA scaffold and delivered via AAV9, both sh10 and sh16 constructs produced substantial *in-vivo* knockdown at doses consistent with prior skeletal muscle RNAi studies.^33,34^ In silico specificity analysis indicated minimal homology to other murine transcripts, supporting target selectivity and a low predicted risk of off-target effects. Preservation of WT-LDB3 or only partial reduction thereof is an important design consideration in dominant disorders, where complete ablation of normal protein function may be deleterious. Allele-selective RNA interference has emerged as a promising strategy for dominant genetic diseases.^34,17,35,36^ Although allele specificity remains desirable, our findings, together with evidence from RNAi therapy in dynamin-2–associated centronuclear myopathy study indicate that partial reduction of total gene expression can also be therapeutically beneficial in selected contexts.^37^ Consistent with this paradigm, Ldb3 haploinsufficiency appears well tolerated,^14,15^ and the favorable outcomes observed with sh16 suggest that partial suppression of both mutant and wild-type transcripts did not produce detectable adverse effects in this model.

Having established mutant allele-level knockdown, we next examined whether molecular rescue translated into structural and functional recovery. Mutant LDB3 suppression resulted in marked rescue across both treatment paradigms. In the preventive therapeutic strategy, early RNAi administration blocked or significantly reduced accumulation of filamin C and CASA-associated aggregates and preserved normal sarcomeric organization and contractile performance. In the advanced-disease therapeutic strategy, RNAi significantly eliminated pre-existing filamin C, BAG3, and HSPA8/Hsc70 aggregates and restored muscle force toward WT levels. The ability to reverse established structural and functional decline is particularly relevant clinically, as most patients present after substantial pathology has developed. Filamin C and its associated CASA components therefore serve not only as mediators of disease but also as dynamic pharmacodynamic markers reflecting therapeutic engagement and recovery.

Mechanistic interpretation integrates with prior work defining the LDB3-PKCα-FLNc mechanosensing complex and the role of CASA in force-induced proteostasis.^9,11,12,19^ LDB3 interacts with FLNc through its Ig-like domains 17–21 and recruits CASA chaperones together with PKCα, forming a structural signaling complex that protects FLNc under mechanical strain and limits proteolysis and aggregation.^38,10^ PKCα-dependent phosphorylation within FLNc Ig-like domain 20 and the hinge-2 region inhibits proteolytic cleavage and stabilizes FLNc under load.^38^ The LDB3 p.Ala165Val mutation does not impair ligand binding but disrupts phosphorylation-dependent signaling within this mechanotransduction module.^10^ Disease pathogenesis therefore reflects failure of mechanosensitive stabilization rather than loss of structural association.

A central mechanistic outcome of RNAi treatment was restoration of PKCα abundance prior to overt myofibrillar disorganization in mutant muscle. To delineate the molecular mechanism underlying the therapeutic benefit of allele-directed RNAi, we performed quantitative phosphoproteomic profiling to define target engagement and downstream signaling restoration. Tandem mass tag-based analysis identified 35 significantly altered phosphoproteins following shRNA treatment, consistent with selective remodeling of defined signaling modules rather than global phosphoproteomic disturbance. Coordinated normalization of phosphorylation was observed across structural Z-disc proteins (ACTN2,^21,39^ TTN,^38,40^ DES,^41^ SYNPO2/SYNPO2L,^38^ ANKRD2),^42^ calcium-handling components ATP2A1 (SERCA1), CASQ1, JPH1, JSRP1),^43–44^ and metabolic enzymes (CKM, GYS1, LDHA),^45,46^ indicating systems-level restoration of sarcomeric and excitation-contraction signaling networks. Our unbiased phosphoproteomic analysis also identified an increase in the PKCα-dependent phosphorylation within FLNc mechanosensing Ig-like domain 20, which was previously shown to prevent its proteolytic cleavage and stabilize FLNc under load.^38^ Notably, increased phosphorylation of established PKCα substrates including FLNc, LDB3, MARCKS, and FXYD1 provides direct molecular evidence of re-engagement of PKCα-dependent signaling. Concurrent normalization of tension-sensitive phosphosites on titin and desmin further supports coordinated reactivation of mechanotransduction, calcium handling, and metabolic support pathways central to Z-disc integrity.

Although kinase enrichment algorithms,^27^ incompletely capture muscle-restricted substrates, PKCα consistently ranked among the top predicted upstream regulators associated with the restored phosphoproteome. This convergence reinforces the interpretation that allele-directed LDB3-RNAi corrects a central regulatory defect and re-establishes the kinase-scaffold environment required for sarcomeric resilience under mechanical load.

Importantly, comparison with the published *in-vivo* human skeletal muscle phosphoproteome,^47^ demonstrated that more than two-third of the phosphoproteins altered by RNAi-sh16 in *Ldb3^Ala165Val/+^* muscle is physiologically phosphorylated in human skeletal muscle. Several treatment-responsive phosphosites correspond exactly to conserved human residues. The conservation of these phosphorylation events across species strengthens the translational relevance of the restored PKCα-dependent signaling network and supports the conclusion that RNAi-mediated correction re-establishes physiologically conserved Z-disc mechanosignaling.

Reactivation of PKCα-dependent phosphorylation also highlights a pathway potentially amenable to complementary pharmacologic strategies. Modulation of CASA dynamics or interventions targeting FLNc turnover may synergize with allele-directed RNAi.^19,11^ The microRNA-based shRNA platform leverages endogenous RNA processing to enhance tolerability and reduce toxicity.^48–50^ AAV9 enables efficient skeletal and cardiac muscle transduction and sustained expression from a single administration.^6^ Given the established clinical experience with AAV9 and the availability of muscle-restricted promoters to confine transgene expression, intramuscular delivery represents a rational and informative strategy for evaluating therapeutic efficacy prior to systemic administration. Although challenges related to systemic delivery, immunogenicity, and long-term durability remain, the present data establishes a compelling translational foundation for further development.

Importantly, the implications of this work extend beyond LDB3-associated skeletal myopathy. LDB3 is not restricted to skeletal muscle but is also expressed broadly in the central and peripheral nervous system, including motor cortex neuron, spinal motor neuron, peripheral nerve, and neuromuscular junction.^51,52^ This broader expression pattern suggests that LDB3-mediated mechanosignaling may have functional relevance beyond striated muscle and provides a potential framework for understanding neurogenic manifestations reported in patients with LDB3 and FLNC mutations. MFMs collectively represent mechanistically convergent disorders characterized by Z-disc instability and CASA dysfunction.^9,8^ The demonstration that allele-directed RNA interference can both prevent disease onset and reverse established pathology suggests broader applicability to dominant myofibrillar and toxic gain-of-function disorders.

In summary, the present study functionally validates and extends our prior mechanistic model of LDB3 as a central regulator of Z-disc mechanosensing. Allele-specific RNA interference targeting LDB3-Ala165Val reconstitutes the LDB3-PKCα-FLNc mechanosignaling hub at the Z-disc by restoring PKCα-dependent signaling, stabilizing FLNc, and normalizing phosphorylation networks organized around the skeletal muscle Z-disc. The demonstration of molecular target engagement, selective phosphoproteomic remodeling, and functional rescue across both preventive and reversal paradigms establishes RNAi-mediated silencing of mutant LDB3 as a disease-modifying strategy rather than a symptomatic intervention. Notably, sustained therapeutic benefit following a single intramuscular AAV9 administration underscores the capacity of vector-mediated RNAi to achieve durable transcript suppression in post-mitotic skeletal muscle. Collectively, these findings provide a mechanistically grounded framework for precision RNAi therapy in autosomal dominant myopathies and support broader application of RNAi-based strategies to disorders driven by dysregulated mechanotransduction.

## Materials and Methods

### DNA constructs, design of siRNA and cell transfection

The DNA construct’s blueprint was provided by authors and cloning, and plasmid sequence confirmation were performed by GENEWIZ (Azenta Life Sciences). The full-length coding sequence (LDB3-Ldel10; of mouse *LDB3* - WT and *LDB3* - A165V (chr14:34571772 C > T; GRCm38/ mm10)^53^ were synthesized and inserted into vector pcDNA3.1 (Invitrogen Life Technology). siRNA sequences were designed using Custom siRNA tool (horizon, PerkinElmer). Oligonucleotide sequences of the siRNAs and scramble siRNA used as control (horizon, PerkinElmer) for in-vitro assays are indicated in Figure 1A and Table S1. HEK-293 cells (2 X 10^5^) seeded in 24 well plates were transfected with 0.2 ug of plasmids *LDB3* - WT and *LDB3* - A165V in separate wells using Lipofectamine 2000 as per the manufacturer’s protocol (Invitrogen Life Technology). After completion of 6 hours of incubation (∼ 70% confluency), cells were transfected with a 20 nM of siRNAs [(si9, si10, si11, si16, si17, and si-Scramble (si-SC)] using the Lipofactamine RNAiMax (Invitrogen Life Technologies) according to manufacturer’s instructions. Transfected HEK-293 cells were incubated at 37 ^0^C for 48 h and harvested for RNA and protein for downstream assays.

### RNA Extraction, cDNA Synthesis, and Real-time quantitative PCR (qPCR)

Total RNA was extracted from transfected HEK-293 cell pellets using the RNeasy Plus Mini Kit (Qiagen) according to the manufacturer’s instructions. The knockdown of *Ldb3* mRNA transcript levels mediated by siRNA was assessed using qPCR on a QuantStudio 6 Flex Real-Time PCR System (Applied Biosystems, Life Technologies). For initial assessment of total *Ldb3* expression, a TaqMan assay targeting constitutively spliced exons 2 and 3 of *Ldb3* (NM_001039071.2; Mm01208763_m1) was used, with *Gapdh* (Mm99999915_g1) serving as the reference gene (Life Technologies).

For analysis of muscle tissue, mouse TA muscles frozen in liquid nitrogen were homogenized in Ambion TRIzol Reagent (Life Technologies) with RNase-free 0.5 mm zirconium oxide beads in a Bullet Blender homogenizer (Next Advance, Troy, NY). Total RNA was then isolated by phase separation with 1-Bromo-3-chloropropane and further purified using the RNeasy Mini Kit (Qiagen). Reverse transcription (RT) was performed in a 20-μL reaction volume containing 1 μg of total RNA, using the SuperScript VILO Master Mix RT kit with random hexamers (Thermo Fisher Scientific), following the manufacturer’s protocol.

To evaluate allele-specific knockdown efficiency of *Ldb3*, total RNA was reverse-transcribed into cDNA and analyzed using a custom TaqMan-based qPCR assay. Amplification of the target region was achieved using the forward primer m*Ldb3*_ex6_Forward (5′- ATCCACGCTCAGTACAACACC-3′) and the reverse primer m*Ldb3*_ex8_Reverse (5′- CCGCCAAGTCTTTCACAGG-3′). Allele-specific detection was performed using two custom TaqMan probes: 165A_Probe (5′-CTCACAGGATGcTATCATGG-3′) to detect the wild-type allele and 165V_Probe (5′-CTCACAGGATGtTATCATGGAC-3′) to detect the mutant allele. Each probe was labeled with a distinct fluorophore and quencher, allowing multiplexed detection within the same reaction.

AS-qPCR assays were optimized and validated using cDNA from *Ldb3*-wild-type, *Ldb3*^Ala165Val/Ala165Val^ homozygous, and *Ldb3*^Ala165Val/+^ heterozygous mice to confirm specificity and amplification efficiency (Figure S1A). *Myom1* (TaqMan; Mm00440394_m1, Applied Biosystems, Life Technologies) served as the endogenous control for normalization. Relative allele expression was determined using the ΔΔCt method and normalized to *Myom1*. All primers and probes were validated to confirm optimal amplification efficiency, and assays were performed within the established dynamic range. Linearity was confirmed across four orders of magnitude using a cDNA dilution series. Each experiment included multiple biological and technical replicates (n ≥ 3), with at least three independent RT-qPCR runs performed per sample.

### Immunoblotting

Transfected HEK-293 cell pellets were homogenized in 2X SDS buffer [100 mM Tris-HCl (pH 6.8), 4% SDS, 20% glycerol, 2% β-Mercaptoethanol, 25 mM EDTA (pH 8.0)], and 0.04% Bromophenol Blue supplemented with protease and phosphatase inhibitor tablets (Roche) by sonication on ice. Mouse TA muscle tissues frozen in liquid nitrogen were homogenized in T-PER lysis reagent (Thermo-Fisher) supplemented with protease and phosphatase inhibitors (Roche) using a polytron homogenizer (ThermoScientific).

Cell and tissue lysates were clarified by centrifugation at 10,000 × g for 5 min at 4 °C, and supernatants were collected. Protein concentration was determined using the Bradford assay (Bio-Rad Laboratories). Equal amounts of protein were resolved on Tris–glycine polyacrylamide gels and transferred to nitrocellulose membranes (iBlot3; Life Technologies). Immunoblotting was performed as previously described, and antibody information, including dilutions, has been reported previously.^10^ At least three independent assays were performed for each sample. Multiple biological replicates were used in each assay.

### miR30-shRNA plasmid construct design and AAV production

Custom-designed mammalian miR30-based shRNA AAV vectors incorporating optimized miR-30 sequences to enable efficient shRNA processing and target gene knockdown were generated by VectorBuilder. Recombinant AAV2/9 serotype vectors were designed by the authors and produced using VectorBuilder’s single-stranded AAV backbone containing a CMV promoter, a Kozak sequence upstream of the EGFP reporter transgene, an miR30-based shRNA cassette, the woodchuck hepatitis virus posttranscriptional regulatory element (WPRE), and a bovine growth hormone (bGH) polyadenylation signal. The following constructs were generated:

(1) pAAV[miR30]-CMV>EGFP:{si10-Ldb3A165V}:WPRE; (2) pAAV[miR30]-CMV>EGFP:{si16-Ldb3A165V}:WPRE; and (3) pAAV[miR30]-CMV>EGFP:Scramble_miR30-shRNA:WPRE. The miR30-based shRNA vector constructs and corresponding shRNA sequences are listed in Table S1.

### Mice and *in-vivo* administration of AAV9-shRNA-miRs

The generation and phenotypic characterization of *Ldb3^Ala165Val/+^* knock-in mice have been described previously.^10^ Wild-type C57BL/6N mice were obtained from Charles River Laboratories (Wilmington, MA). Mice were housed in the animal care facility of the National Institute of Neurological Disorders and Stroke (NINDS) under standard conditions, maintained on a 12 h light/dark cycle with unrestricted access to standard chow and water. Both male and female mice were included in all experimental groups, as prior studies demonstrated comparable myofibrillar myopathy phenotypes across sexes in this model.^10^ All procedures were approved by the Institutional Animal Care and Use Committee (IACUC) of NINDS and conducted in accordance with NIH guidelines.

For *in-vivo* studies, *Ldb3^Ala165Val/+^* mice received intramuscular injections of AAV9-shRNA-miR vectors into the TA muscle at defined ages and doses. In all treated mice, one TA muscle received AAV9-shRNA-miR_10 or AAV9-shRNA-miR_16 and served as the treatment arm, while the contralateral TA muscle was injected with AAV9-shRNA-miR_Scr and served as an internal control. For dose-ranging experiments, mice at 3 months of age were injected with either 5 × 10¹¹ or 5 × 10¹² vector genomes (vg)/kg of AAV9-shRNA-miR_10 or AAV9-shRNA-miR_16 (n = 3-6 mice per group), and tissues were collected one month after injection.

For studies in advanced-stage disease, mice were treated at 5 months of age with a single intramuscular injection of 5 × 10¹² vg/kg of either AAV9-shRNA-miR_16 (5M-1M, n = 8; 5M-3M, n = 12) or AAV9-shRNA-miR_10 (5M-1M, n = 11; 5M-3M, n = 9) and analyzed for different assays either one month or three months post-injection. For early-intervention studies, mice were injected at 3 months of age with a single dose of 5 × 10¹² vg/kg of either AAV9-shRNA-miR_16 (3M–1M, n = 15; 3M-3M, n = 13) or AAV9-shRNA-miR_10 (3M-1M, n = 13; 3M-3M, n = 10) and evaluated one month or three months following injection. In addition to mutant cohorts, a total of 18 age-matched WT mice were included as controls across immunoblotting and *in-vivo* muscle contractility assays. An additional cohort of WT mice (n = 6) received intramuscular injections of AAV9-shRNA-miR_10 or AAV9-shRNA-miR_16 and were transcardially perfused for histological and IF analyses to assess muscle immunostaining outcomes. For *in-vivo* muscle function studies, a subset of mice treated at 3 months of age with AAV9-shRNA-miR_10 or AAV9-shRNA-miR_16 was evaluated four months post-injection (3M-4M) for isometric torque measurements, with treated TA muscles compared against contralateral scramble-treated controls.

Intramuscular injections were performed into the lateral compartment of the TA muscle using a 25–27-gauge needle. Mice were anesthetized using either isoflurane (2-5%) or a ketamine/xylazine mixture (30-40 mg/kg ketamine and 2.5-3 mg/kg xylazine; injection volume 110– 150 µL), depending on the experimental session, and depth of anesthesia was confirmed by hindlimb toe pinch prior to injection. A single intramuscular injection of 20 µL was administered into each TA muscle. Following administration, mice were placed on a warming pad and monitored until full recovery and normal ambulation were observed. The contralateral TA muscle received an equivalent dose of AAV9-shRNA-miR_Scr and served as an internal control. Viral doses are reported as vector genomes (vg) per kilogram body weight.

### Digital PCR for viral genome quantification

Genomic DNA was extracted using the QIAamp DNA Mini Kit (Qiagen) and then any residual RNA was digested with RNase (Roche) and purified. Digital PCR (dPCR) was performed to determine the AAV genome copy number per host genome using *Polr2a* as a diploid reference gene. The dPCR reactions were loaded onto the QIAcuity Nanoplate 8.5k 96-well plates and run on a QIAcuity One system (Qiagen, Germany) according to the manufacturer’s instructions.

AAV genome copies were quantified using the dPCR_EGFP assay (Integrated DNA Technologies), which included the following components: forward primer 5′-GCACAAGCTGGAGTACAACTA-3′, reverse primer 5′-TGTTGTGGCGGATCTTGAA-3′, and a FAM-labeled probe 5′-AGCAGAAGAACGGCATCAAGGTGA-3′. Host genome copy number was determined using the *Polr2a* assay (Integrated DNA Technologies), with forward primer 5′-GACTCCTTCACTCACTGTCTTC-3′, reverse primer 5′-TCTTGCTAGGCAGTCCATTATC-3′, and a HEX-labeled probe 5′-ACGAGATGC/ZEN/TGAAAGAGCCAAGGT-3′. All assay components were included in the same reaction mixture. dPCR was performed under the following cycling conditions: 2 min at 95° C, followed by 40 cycles of 15 sec at 95 °C and 30 sec at 64 °C. Following amplification, fluorescence imaging was performed using exposure and gain settings of 500 ms and 6 ms, respectively, in the green (FAM) and yellow (HEX) channels. Negative controls lacking EGFP template and no-template controls (NTCs) were included in each run.

### Histology and Immunofluorescence analysis

At experimental end points, TA muscles were rapidly frozen in dry ice-cooled isopentane. Transverse muscle sections (10 μm thick) were cryosectioned and stained with MGT and NADH-TR for histological analysis. For IF, 10-μm-thick frozen transverse sections were fixed in 4% PFA for 10 min, permeabilized, and blocked with 2.5% normal goat serum and 0.2% Triton X-100 in PBS for 1 h at room temperature. Sections were incubated overnight at 4 °C with primary antibodies against LDB3, filamin C, BAG3, HSPA8, and EGFP (Cell Signaling Technology, ab184601); antibody details and dilutions were as described previously.^10^

Images were acquired using a Leica Stellaris confocal microscope equipped with a 40×/NA 1.3 oil immersion Plan-Apochromat objective lens and processed using Adobe Photoshop Creative Cloud v2017. Identical imaging settings were applied to all images within a given experiment, and comparisons were made between sections stained on the same slide. For longitudinal muscle section analysis, mice were transcardially perfused with 4% PFA in PBS and skeletal muscle tissues were post-fixed overnight at 4° C. Tissues processing, IF staining and image acquisition were performed as previously described.^10^

To assess myonuclear localization, transverse TA muscle sections were stained with wheat germ agglutinin-Alexa Fluor 488 conjugate (WGA; Thermo Fisher Scientific) to label muscle membranes and mounted using ProLong Diamond Antifade Mountant with DAPI (Thermo Fisher Scientific) to visualize nuclei. Images were acquired using a 20×/NA 0.4 objective on a Stellaris confocal microscope and analyzed using Leica Application Suite (LAS) X software. Adjacent fields were digitally stitched with a 10% overlap to generate a composite image of the entire TA cross-section. Myonuclei were quantified in comparable regions of contralateral scramble-control (SC) and treated TA muscles, excluding fibers at the tissue edges. For sh16-treated mice (5M-3M; n = 5 per group), an average of 339 fibers per TA muscle were analyzed. For sh10-treated mice (5M-3M; n = 3 per group), an average of 577 fibers per TA muscle were analyzed. Counting was performed using ImageJ/Fiji software (NIH).

To quantify sarcoplasmic filamin C protein aggregates, confocal microscopy was performed at 40× magnification on longitudinal sections. Five or more randomly selected fields, evenly distributed across each muscle section, were analyzed. The percentage of fibers containing filamin C aggregates was determined by counting ≥50 fibers per muscle section for both control; n=3 and treated; n=3 groups.

### *In-vivo* measurement of isometric torque

The isometric contractile properties of the TA muscle were assessed *in-vivo* using an Aurora Scientific model 1300A dual-mode muscle lever system. Mice were placed in a supine position on a thermostatically controlled platform and anesthetized with inhaled isoflurane (4-5% for induction for less than 1 minute, followed by approximately 2% for maintenance via a nose cone, delivered in 100% O at 1-1.5 L/min).

The knee joint was immobilized, and the foot was secured to a footplate connected to the motor shaft, with the ankle positioned at 90° relative to the tibia. TA muscle contraction (dorsiflexion) was elicited by stimulation of the fibular (common peroneal) nerve using subcutaneous electrodes. A force–frequency protocol was applied by delivering electrical stimulation at frequencies ranging from 25 to 250 Hz in 25-Hz increments, with a 1-min rest period between stimulations. Maximal isometric torque was recorded at full tetanic contraction at each stimulation frequency.

For advanced-stage studies, *Ldb3^Ala165Val/^*^+^ mice treated at 5 months of age were evaluated using a representative cohort, with *in-vivo* muscle contractility measured in sh10-treated TA muscles at the 5M-1M time point. Treated TA muscles were compared with contralateral scramble-control (SC) TA muscles and age-matched WT-TA muscles.

For early-treatment studies, *in-vivo* muscle contractility assays were conducted in sh10-treated mice at the 3M-1M and 3M-4M time points, as well as in sh16-treated mice at the 3M-4M time point. In these early-intervention cohorts, treated TA muscles were analyzed using the same comparative framework, with functional recovery assessed relative to contralateral scrambled-treated TA muscles and age-matched PBS treated WT-TA muscles. Group sizes for functional analyses were n = 3-4 per group (WT, contralateral SC, and sh10- or sh16-treated).

### Global Phosphoproteomic Analysis

#### Experimental Design and Sample Preparation

Paired phosphoproteomic analyses were performed using TA muscles from *Ldb3^Ala165Val/+^* mice treated with AAV9-shRNA-miR_16 (sh16) and the contralateral scrambled-treated control (SC) muscles at the 3M-1M time point. Each animal served as its own internal control, enabling paired comparisons that minimized inter-animal variability. Four biological replicates were analyzed (n= 4 mice), with each mouse contributing one sh16-treated TA muscle and one contralateral SC-TA muscle.

TA muscles were rapidly dissected, snap-frozen in liquid nitrogen, and stored at −80°C until processing. Frozen tissue was homogenized on ice in urea-based lysis buffer (8 M urea, 50 mM Tris-HCl pH 8.0, 75 mM NaCl, 1 mM EDTA) supplemented with protease and phosphatase inhibitor cocktails (Roche). Cytosolic and myofibrillar protein fractions were isolated using standard differential solubilization procedures and processed separately at the experimental level. Protein concentrations were determined using a BCA assay (Thermo Fisher Scientific). Equal amounts of protein (100 µg per sample) from each fraction were reduced, alkylated, and digested overnight at 37° C with sequencing-grade trypsin (1:50 enzyme-to-protein ratio). Peptides were desalted using Waters Oasis HLB cartridges, labeled with TMTpro reagents (Thermo Fisher Scientific), and pooled separately for cytosolic and myofibrillar fractions to preserve compartment-specific information.

#### Phosphopeptide Enrichment and Mass Spectrometry

Phosphopeptides were enriched from pooled samples using a TiO (Titanium dioxide)-based method followed by an IMAC approach. The kits used were the ThermoFisher TiO Phosphopeptide Enrichment Kit and the Fe-NTA Phosphopeptide Enrichment Kit. Enriched phosphopeptide fractions and corresponding non-enriched peptide fractions were analyzed by nanoLC-MS/MS using an Ultimate 3000 UHPLC system coupled to an Orbitrap Fusion Lumos mass spectrometer (Thermo Fisher Scientific). Peptides were separated on a C18 analytical column (75 μm × 25 cm, 2 μm particle size) using a 90-min linear gradient from 3% to 35% acetonitrile in 0.1% formic acid at a flow rate of approximately 300 nL/min. Data were acquired in data-dependent acquisition mode using higher-energy collisional dissociation.

#### Database Searching, Quantification, and Normalization

Raw mass spectrometry data were processed using Proteome Discoverer (v2.4) and searched against the UniProt Mus musculus reference proteome using a target-decoy strategy. Search parameters included up to two missed cleavages, static modifications of N-Ethylmaleimide and TMTpro labeling (N-termini and lysine residues), and dynamic phosphorylation of serine, threonine, and tyrosine residues. Peptide and protein identifications were filtered at a 1% false discovery rate.

Relative quantification was based on TMT reporter ion intensities. Phosphosite abundances were normalized to the corresponding total protein abundance within the same subcellular compartment (cytosolic or myofibrillar), with additional global normalization based on total peptide intensity to correct for sample loading and labeling variability. This compartment-matched normalization ensured that phosphorylation changes reflected relative modulation within structurally and functionally distinct protein pools. Normalized phosphopeptide intensities were converted to log ratios for treated versus control samples and subsequently transformed to linear fold changes for biological interpretation. Processed datasets, including normalized intensities, log ratios, and linear fold changes, were used for downstream visualization, pathway enrichment, and kinase inference analyses.

#### Filtering Criteria, Pathway Enrichment, and Kinase Inference

Cytosolic and myofibrillar phosphoproteomic datasets were analyzed separately at the experimental level and subsequently integrated for systems-level analyses. Because both fractions were derived from the same biological samples, processed in parallel, and normalized to their respective total protein pools, phosphorylation differences reflect internally controlled, paired measurements rather than independent observations.

Differential phosphorylation between sh16-treated and SC-TA muscles was assessed using Welch’s two-sample t test (n = 4 biological replicates per group). For pathway enrichment analysis, treatment-responsive phosphoproteins were defined using a stringent Welch’s t test with multiple-testing correction (adjusted p < 0.05), yielding 35 unique phosphoproteins. Of these, 30 exhibited phosphorylation changes ≥1.25-fold, consistent with biologically meaningful modulation in a TMT-based phosphoproteomic dataset. Pathway enrichment analysis was performed using Enrichr-KG, a knowledge-graph-based extension of the Enrichr platform, with nominal enrichment p values and Benjamini-Hochberg false discovery rate-adjusted q values calculated within each annotation library.^26,54^

For kinase enrichment analysis (KEA3), a biologically anchored feature-selection strategy was applied. Phosphoproteins were selected using a nominal Welch’s t test (p < 0.05) combined with a ≥1.4-fold change threshold, corresponding to the magnitude observed for filamin C (FLNc), a well-established PKCα-regulated substrate. This input set was used for upstream kinase inference using KEA3, which integrates motif-based kinase-substrate predictions and cophosphorylation associations derived from curated phosphoproteomic and protein-protein interaction databases.^27^ Kinase ranking and statistical significance were determined using KEA3’s internal enrichment metrics rather than external hypothesis-testing thresholds.^27^

### Statistical Analysis

Statistical analyses were performed using GraphPad Prism (versions 10 and 10.6.0 [890]; GraphPad Software, Boston, MA). For comparisons between two groups in non-phosphoproteomic experiments, unpaired two-tailed Student’s t-tests and Welch’s t-tests were used. Data are presented as mean ± SEM or mean ± SD, as specified in the text and corresponding figure legends, and individual values are shown in bar-scatter plots where indicated. The statistical tests used, sample sizes (n), number of experimental replicates, and exact P values are provided in the figure legends. For phosphoproteomic datasets, differential phosphorylation between treated and scrambled-treated control samples was assessed using Welch’s two-sample t test. Multiple testing correction was performed where indicated using the Benjamini-Hochberg method. Criteria for defining treatment-responsive phosphoproteins, as well as pathway and kinase enrichment analyses including fold-change thresholds and use of nominal versus adjusted p values-are detailed in the Methods section. Statistical significance was defined as p < 0.05 and is indicated as follows: p < 0.05, p < 0.01, p < 0.001, and p < 0.0001.

## Data Availability

All relevant data are included in the manuscript and Supplemental Information. Any other datasets associated with this study are accessible through the corresponding author upon reasonable request.

## Acknowledgments

This research was supported by the Intramural Research Program of the National Institute of Neurological Disorders and Stroke (NINDS), National Institutes of Health (NIH). The contributions of the NIH authors are considered Works of the United States Government. The findings and conclusions presented in this paper are those of the authors and do not necessarily reflect the views of the NIH or the U.S. Department of Health and Human Services. The authors thank Dr. Carsten G. Bönnemann for serving as interim Principal Investigator and for oversight of this work. We also acknowledge Drs. Vincent Schram and Ling Yi of the NICHD Imaging Core Facility, NIH, for technical assistance with mouse perfusion.

## Author contributions

A.M. and P.P. conceived and designed the study and experiments. P.P., J.P., J.H., A.K., M.P., D.S., and Y.L. performed the experiments. K.J. conducted the formal proteomics data analysis. P.P. and A.M. analyzed and interpreted the data and drafted the manuscript. All authors reviewed and approved the final manuscript.

