## Supplemental figures and tables for "AAV-Delivered RNAi Targeting Mutant LDB3 Prevents and Reverses Myofibrillar Myopathy through Mechanosignaling Restoration"

### Supplemental Information

#### Supplemental figures and legends

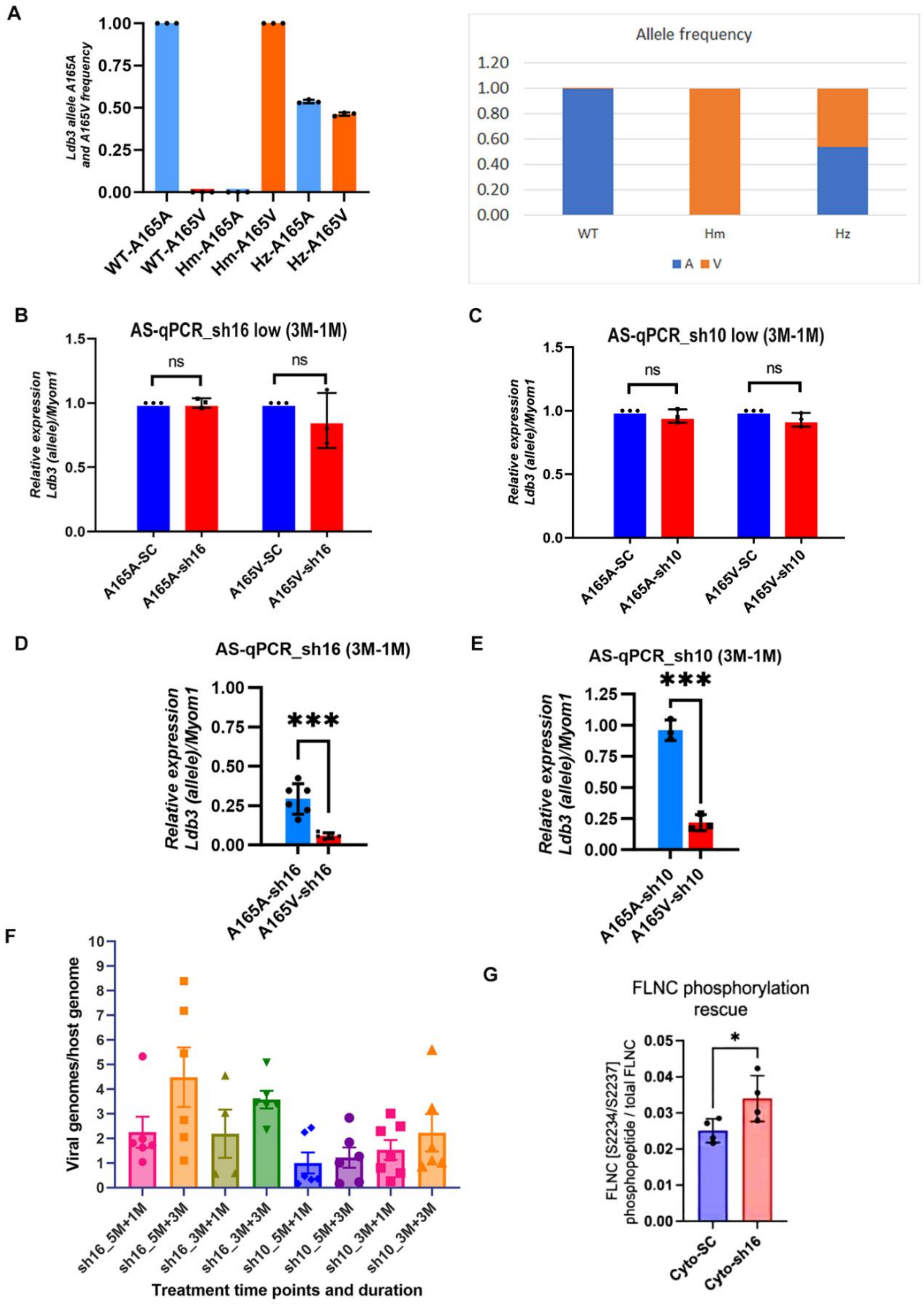

**Figure S1. Allele-specific qPCR optimization, AAV therapeutic dose determination, AAV transduction efficiency, and FLNc phosphorylation following AAV9-shRNAmiR treatment.**

(A) Validation of allele-specific RT-qPCR using cDNA from WT, *Ldb3*$^{Ala165Val/Ala165Val}$ (Homozygous), and *Ldb3*$^{Ala165Val/+}$ (Heterozygous) mice confirms assay specificity for WT-A165A and A165V alleles.

Three-month-old *Ldb3*$^{Ala165Val/+}$ mice received a single intramuscular injection of sh10 or sh16 into the TA muscle; contralateral TA muscles received scramble control (SC). (B and C) AS-qPCR analysis showing relative expression of WT (A165A) and mutant *Ldb3*-A165V transcripts at one-month post-injection following low-dose AAV9-shRNAmiR ($5 \times 10^{11}$ vg/kg) compared with SC-treated controls in sh16 and sh10 treatment groups (n = 3 mice per group). (D and E) AS-qPCR analysis showing relative expression of WT (A165A) and mutant *Ldb3*-A165V transcripts at one-month post-injection following high-dose AAV9-shRNAmiR ($5 \times 10^{12}$ vg/kg) compared with SC-treated controls in sh16 and sh10 treatment groups (n = 3–6 mice per group). (F) AAV vector genome copies per host genome across treatment groups in sh16- and sh10-treated *Ldb3*$^{Ala165Val/+}$ mice at advanced stage (5 months) and early stage (3 months), analyzed at 1 and 3 months post-injection to assess transduction efficiency (n = 4-6 mice per group). (G) Phosphoproteomic fold-change data presented as a bar-scatter plot showing increased phospho-FLNc (S2234/S2237)/total FLNc ratio following sh16 treatment (3-month-old mice, one-month post-injection) compared with SC (n = 4 mice per group). Student's t test: $p = 0.0457$. Data are presented as mean ± SD (A-G). Statistical significance was determined using Welch's t-Test (B and C) and unpaired two-tailed Student's t tests (D, E and G) (*$P < 0.05$ and ***$P < 0.001$), ns = not significant.

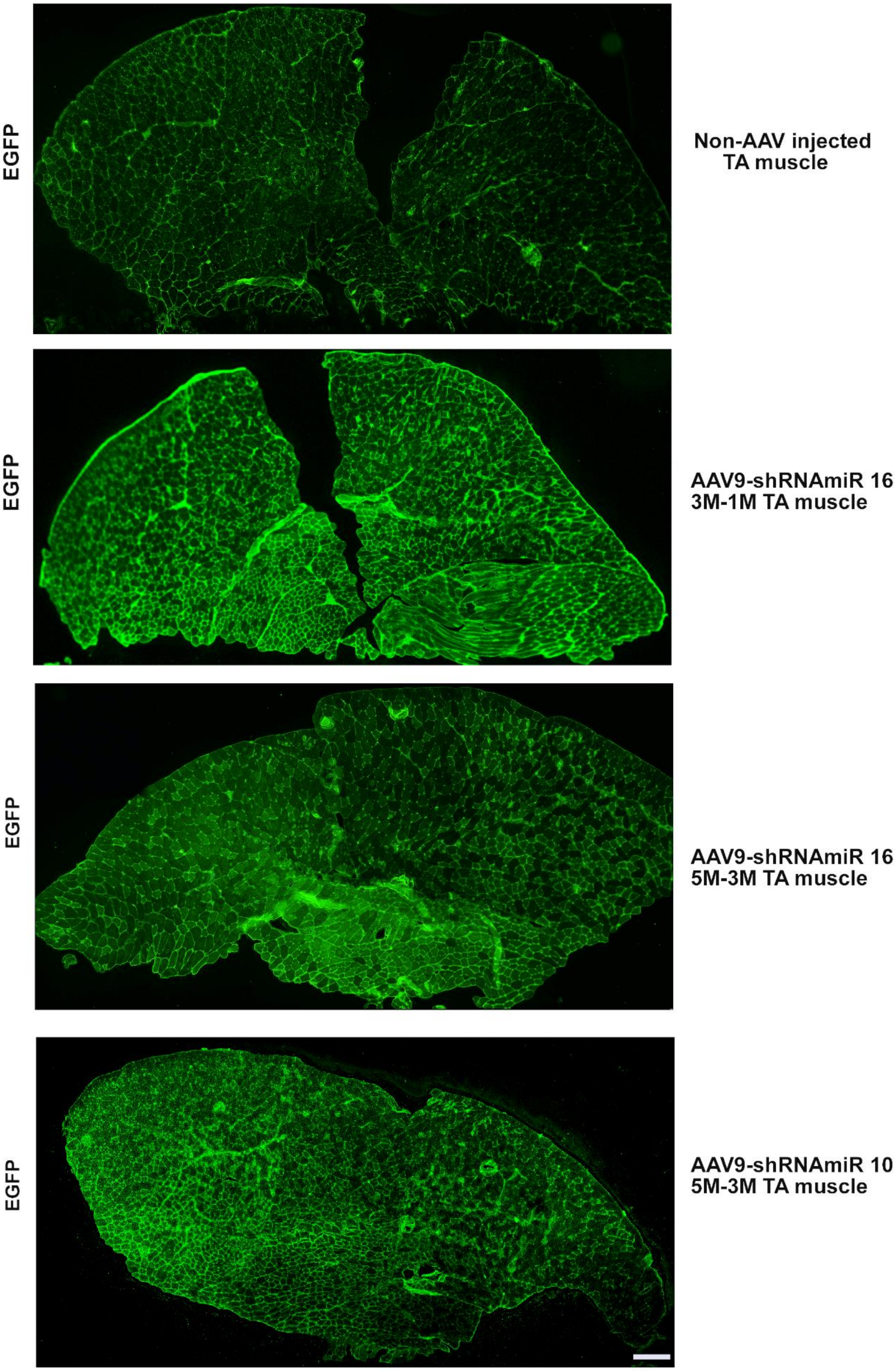
EGFP
Non-AAV injected
TA muscle
EGFP
AAV9-shRNAmiR 16
3M-1M TA muscle
EGFP
AAV9-shRNAmiR 16
5M-3M TA muscle
EGFP
AAV9-shRNAmiR 10
5M-3M TA muscle

**Figure S2: Efficient skeletal muscle transduction following AAV9-shRNAmiR administration.**

Representative IF images of EGFP expression in 8-μm transverse sections of TA muscle from *Ldb3*$^{Ala165Val/+}$ mice following AAV9-shRNAmiR delivery. The top panel shows non-AAV-injected TA muscle stained for EGFP (negative control). Subsequent panels show EGFP expression in TA muscles at 1-month post-injection in 3-month-old mice (3M-1M; age 4 months) and at 3 months post-injection in 5-month-old mice (5M-3M; age 8 months) treated with AAV9-shRNAmiR16 or AAV9-shRNAmiR10. Broad and homogeneous EGFP expression throughout the transverse section demonstrates efficient AAV-mediated skeletal muscle transduction in both early-stage (3M-1M and 3M-3M) and advanced-stage (5M-1M and 5M-3M) treatment paradigms. Scale bar: 250 μm.

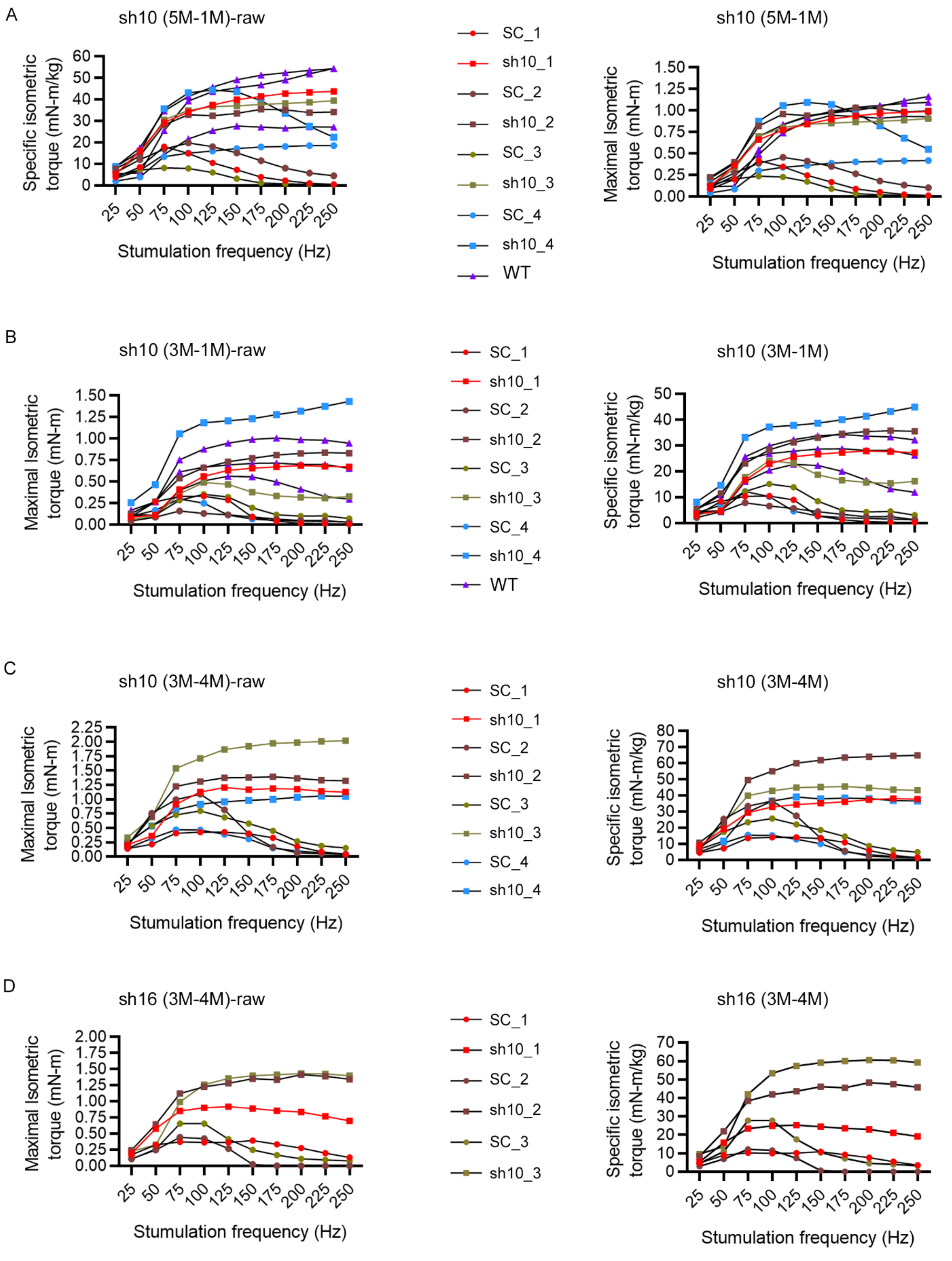

A
sh10 (5M-1M)-raw
Specific isometric torque (mN-m/kg)
Stumulation frequency (Hz)
SC_1
sh10_1
SC_2
sh10_2
SC_3
sh10_3
SC_4
sh10_4
WT
sh10 (5M-1M)
Maximal Isometric torque (mN-m)
Stumulation frequency (Hz)
B
sh10 (3M-1M)-raw
Maximal Isometric torque (mN-m)
Stumulation frequency (Hz)
SC_1
sh10_1
SC_2
sh10_2
SC_3
sh10_3
SC_4
sh10_4
WT
sh10 (3M-1M)
Specific isometric torque (mN-m/kg)
Stumulation frequency (Hz)
C
sh10 (3M-4M)-raw
Maximal Isometric torque (mN-m)
Stumulation frequency (Hz)
SC_1
sh10_1
SC_2
sh10_2
SC_3
sh10_3
SC_4
sh10_4
sh10 (3M-4M)
Specific isometric torque (mN-m/kg)
Stumulation frequency (Hz)
D
sh16 (3M-4M)-raw
Maximal Isometric torque (mN-m)
Stumulation frequency (Hz)
SC_1
sh10_1
SC_2
sh10_2
SC_3
sh10_3
sh16 (3M-4M)
Specific isometric torque (mN-m/kg)
Stumulation frequency (Hz)

**Figure S3. Force frequency relationship data from 25 to 250 Hz for each treatment group in *Ldb3*$^{Ala165Val/+}$ and WT-TA muscle.**

Force-frequency relationships showing raw maximal isometric force (left) and body-weight–normalized specific isometric torque (right) generated by TA muscles in response to increasing stimulation frequencies. (A) sh10 treatment (5M-1M), (B) sh10 treatment (3M-1M), (C) sh10 treatment (3M-4M) and (D) sh16 treatment (3M-4M). Within each panel, shRNAmiR-treated TA muscles are compared with contralateral scramble control (SC)-treated muscles. PBS-treated WT mice are included where indicated (n = 3–4 per group) and serve as a reference for the sh10 and sh16 (3M–4M) treatment groups shown in the main figures.

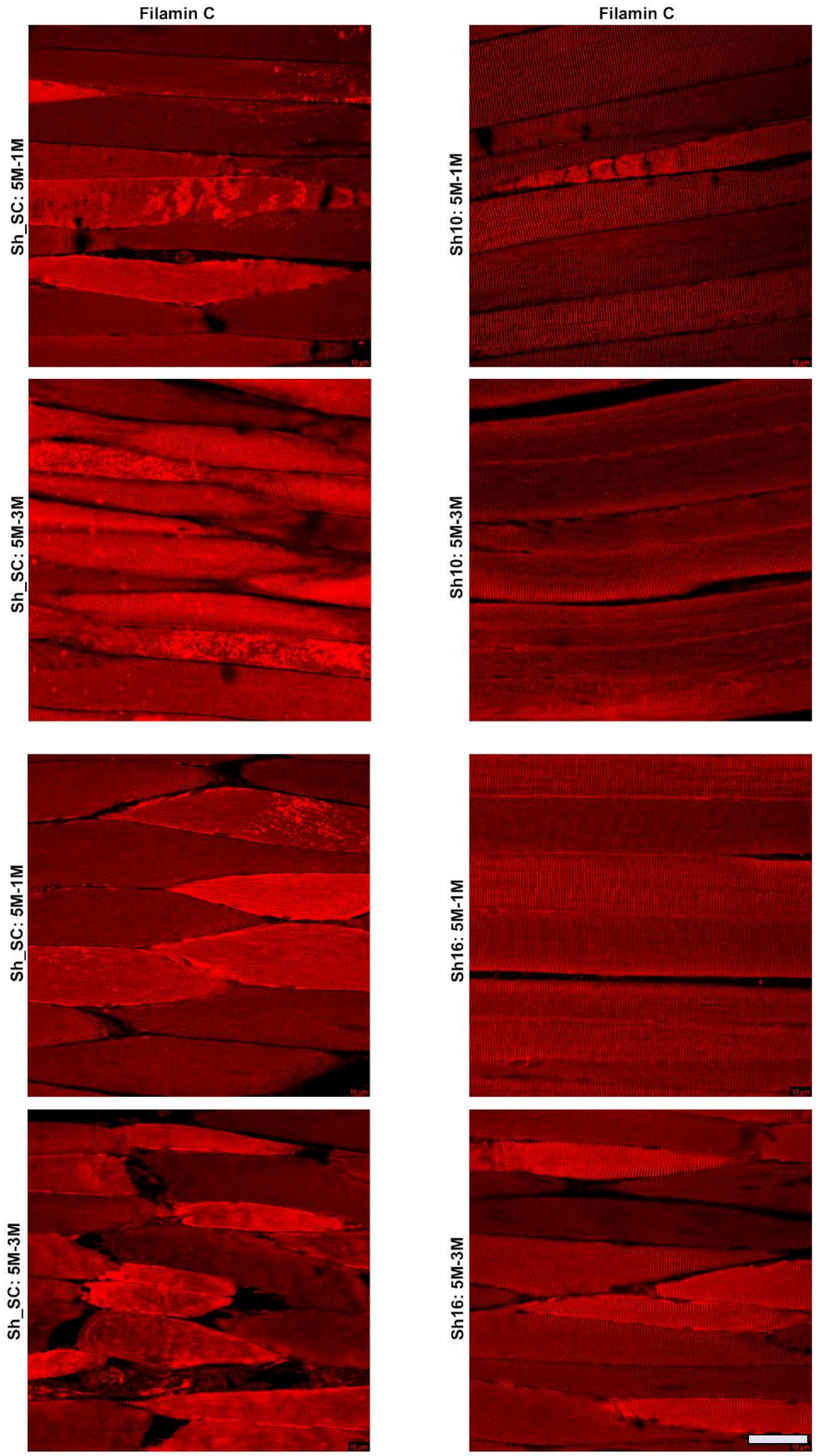

**Figure S4. Therapeutic *Ldb3* suppression reduces filamin C aggregation in advanced-stage *Ldb3*$^{Ala165Val/+}$ mice.**

Representative IF images of filamin C in PFA-perfused longitudinal TA muscle sections from advanced-stage (5-month-old) *Ldb3*$^{Ala165Val/+}$ mice treated with sh-SC, sh10, or sh16 and analyzed at 1 month (5M-1M; age 6 months) or 3 months (5M-3M; age 8 months) post-injection (n = 4-6 per group).

Left panels (sh-SC) show extensive filamin C aggregation with progressive Z-disc disorganization at 5M-1M and marked Z-disc disruption at 5M-3M. Right panels show the corresponding sh10- (upper rows) and sh16-treated (lower rows) muscles at both time points, demonstrating marked reduction of filamin C aggregates and restoration of Z-disc architecture relative to SC-treated controls. Scale bars represent 50 μm.

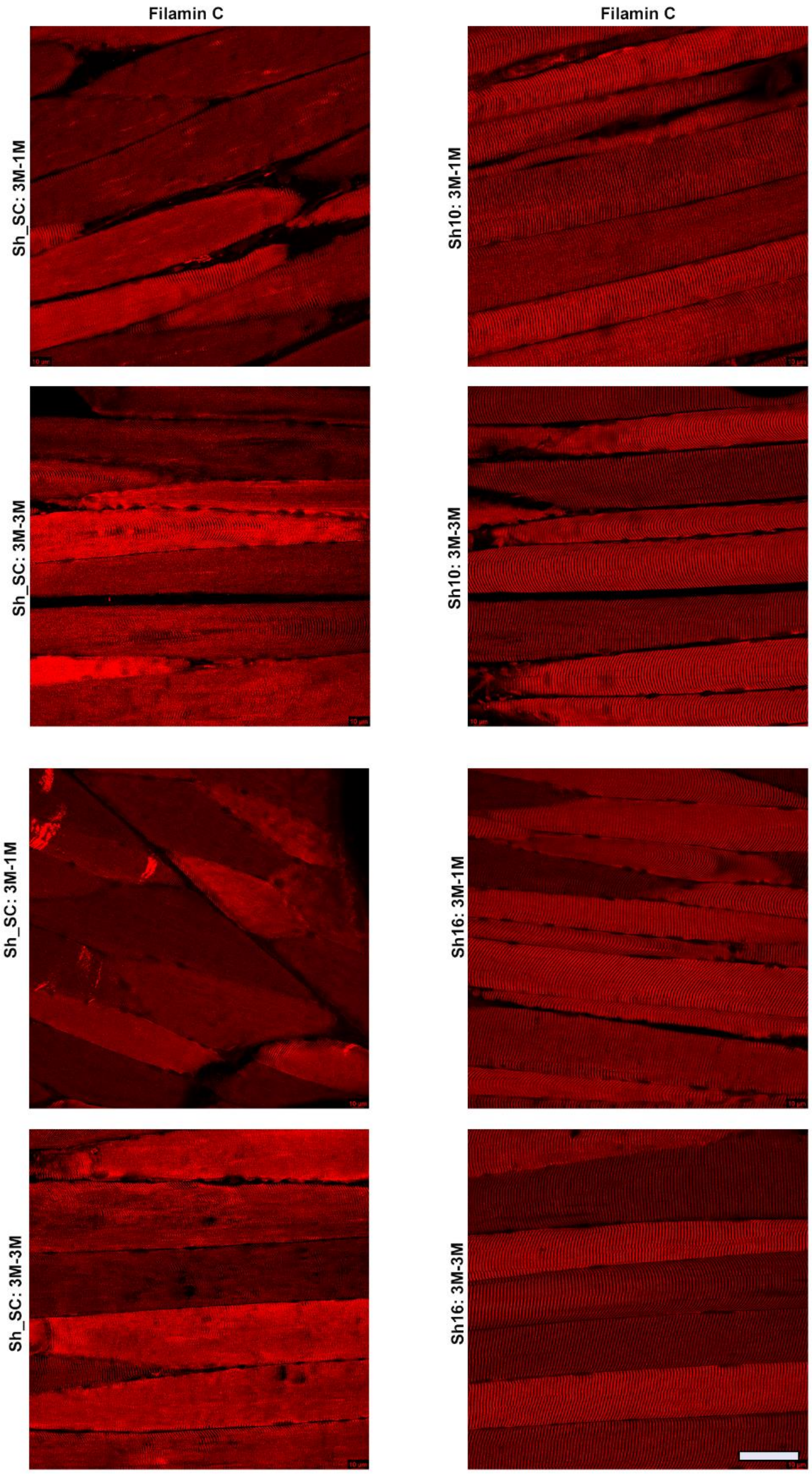

**Figure S5: Early-stage mutant *Ldb3* suppression prevents filamin C aggregation and Z-disc disruption in *Ldb3*$^{Ala165Val/+}$ mice.**

Representative IF images of filamin C in PFA-perfused longitudinal TA muscle sections from early-stage (3-month-old) *Ldb3*$^{Ala165Val/+}$ mice treated with sh-SC, sh10, or sh16 and analyzed at 1 month (3M-1M; age 4 months) or 3 months (3M-3M; age 6 months) post-injection (n = 4-6 per group). Left panels (sh-SC) show the onset of filamin C aggregation at 3M-1M, with aggregates extending across multiple Z-discs, and increased aggregation with evident Z-disc disorganization at 3M-3M. Right panels show the corresponding sh10- (upper rows) and sh16-treated (lower rows) muscles at both time points, in which filamin C aggregation is absent or markedly reduced and Z-disc architecture remains preserved. Scale bars represent 50 μm.

**Supplemental tables**

**Table S1: siRNA design and shRNA sequence BLAST analysis**

**Tab 1.** siRNA scoring based on algorithmic prediction

**Tab 2.** Complete BLAST output for shRNA sequence homology.

**Tab 3.** Top predicted off-target matches identified by shRNA sequence BLAST analysis.

**Tab 4.** Vector maps

**Table S2: TMT-based phosphoproteomic analysis of differential phosphorylation between scrambled control- and LDB3-sh16-treated TA muscle.**

Combined cytosolic and myofibrillar phosphosite-level datasets (Tabs 2 and 3) were used to generate the 35 unique significantly altered phosphoproteins listed in Tab 1 for KEGG pathway enrichment analysis, whereas combined cytosolic and myofibrillar datasets (Tabs 4 and 5) were used for KEA3 kinase enrichment analysis.

**Tab 1.** List of 35 unique significantly altered phosphoproteins with phosphosite details (≥1 significant phosphosite per protein) used for KEGG pathway enrichment analysis.

**Tab 2.** Cytosolic fraction phosphosite-level dataset, including annotated sequences, modification sites, quantitative values ($log_2$ means and linear differences), and Welch's t test uncorrected and corrected p values.

**Tab 3.** Myofibrillar fraction phosphosite-level dataset, including annotated sequences, modification sites, quantitative values ($log_2$ means and linear differences), and Welch's t test uncorrected and corrected p values.

**Tab 4.** Significantly regulated cytosolic phosphoproteins used for KEA3 kinase enrichment analysis.

**Tab 5.** Significantly regulated myofibrillar phosphoproteins used for KEA3 kinase enrichment analysis.

**Table S3.** PKCα-responsive structural, mechanosensory, and metabolic phosphoproteins identified by TMT-based phosphoproteomic analysis of LDB3-sh16–treated TA muscle

**Table S4.** Enrichr-KG analysis of proteins with treatment-responsive phosphosites in sh16-treated *Ldb3*$^{Ala165Val/+}$ TA muscle

**Table S5.** Complete cytosolic fraction phosphosite-level dataset from TMT-based phosphoproteomic analysis of scrambled control– and LDB3-sh16–treated TA muscle.

**Table S6.** Complete myofibrillar fraction phosphosite-level dataset from TMT-based phosphoproteomic analysis of scrambled control– and LDB3-sh16–treated TA muscle.

**siRNA scoring**

| | | GC content (%) | | #'A/U' at position 15-19 | | Absence of Internal repeats [Tm (°C)] | | Position 19 with 'A' | | Position 3 with 'A' | | Position 10 with 'U' | | A base other than 'G' or 'C' at position 19 | | A base other than 'G' at position 13 | | |
|---|---|---|---|---|---|---|---|---|---|---|---|---|---|---|---|---|---|---|
| **siRNA** | **siRNA Sense sequence 5' --> 3'** | **value** | **Points** | **value** | **points** | **value** | **points** | **y/n[a]** | **points** | **y/n[a]** | **points** | **y/n[a]** | **points** | **y/n[a]** | **points** | **y/n[a]** | **points** | **Score** |
| 1. Acatcctgtgagtacatgc | ACA UCC UGU GAG UAC AUG CUU | 47.4 | 1 | 16, 17 | 2 | 47 | 1 | C | 0 | A | 1 | G | 0 | Y | -1 | U | 0 | 3 |
| 2. aAcatcctgtgagtacatg | AAC AUC CUG UGA GUA CAU GUU | 42.1 | 1 | 15,17,18 | 3 | 44 | 1 | G | 0 | C | 0 | U | 1 | Y | -1 | G | -1 | 3 |
| **3.** ataAcatcctgtgagtaca | UAA CAU CCU GUG AGU ACA UUU | 36.8 | 1 | 15,16,18, | 4 | 42 | 1 | U | 0 | A | 1 | G | 0 | N | 0 | A | 0 | 6 |
| **4.** ataAcatcctgtgagtaca | AUA ACA UCC UGU GAG UAC AUU | 33.3 | 1 | 16, 17, 19 | 3 | 42 | 1 | A | 1 | A | 1 | U | 1 | N | 0 | G | 0 | 7 |
| **5.** gataAcatcctgtgagta | GAU AAC AUC CUG UGA GUA CUU | 38.1 | 1 | 15,17,18 | 3 | 44 | 1 | C | 0 | U | 0 | C | 0 | Y | -1 | U | 0 | 3 |
| 6. tgataAcatcctgtgagta | UGA UAA CAU CCU GUG AGU AUU | 38.8 | 1 | 16,18,19 | 3 | 42 | 1 | A | 1 | A | 1 | C | 0 | N | 0 | G | -1 | 5 |
| 7. atgataAcatcctgtgag | AUG AUA ACA UCC UGU GAG UUU | 38.8 | 1 | 15,17,19 | 3 | 42 | 1 | U | 0 | G | 0 | U | 1 | N | 0 | U | 0 | 5 |
| 8. catgataAcatcctgtga | CAU GAU AAC AUC CUG UGA GUU | 42.1 | 1 | 16,18 | 2 | 44 | 1 | G | 0 | U | 0 | A | 0 | Y | -1 | C | 0 | 2 |
| 9. ccatgataAcatcctgtga | CCA UGA UAA CAU CCU GUG AUU | 38.1 | 1 | 15, 17, 17 | 3 | 44 | 1 | A | 1 | A | 1 | C | 0 | N | 0 | C | 0 | 6 |
| 10. tccatgataAcatcctgtg | UCC AUG AUA ACA UCC UGU GUU | 38.1 | 1 | 16, 18 | 2 | 44 | 1 | G | 0 | C | 0 | A | 0 | Y | -1 | U | 0 | 2 |
| 11. gtccatgataAcatcctgt | GUC CAU GAU AAC AUC CUG UUU | 38.1 | 1 | 17, 19 | 2 | 44 | 1 | U | 0 | C | 0 | A | 0 | N | 0 | A | 0 | 3 |
| 12. cgtccatgataAcatcctg | CGU CCA UGA UAA CAU CCU GUU | 47.4 | 1 | 15,18 | 2 | 47 | 1 | G | 0 | U | 0 | U | 1 | Y | -1 | C | 0 | 3 |
| 13. gcgtccatgataAcatcct | GCG UCC AUG AUA ACA UCC UUU | 47.4 | 1 | 15,16,19 | 3 | 47 | 1 | U | 0 | G | 0 | A | 0 | N | 0 | A | 0 | 4 |
| 14. ggcgtccatgataAcatcc | GGC GUC CAU GAU AAC AUC CUU | 52.6 | 1 | 16,17 | 2 | 47 | 1 | C | 0 | C | 0 | G | 0 | Y | -1 | A | 0 | 2 |
| 15. tggcgtccatgataAcatc | UGG CGU CCA UGA UAA CAU CUU | 47.4 | 1 | 15,17,18 | 3 | 47 | 1 | C | 0 | G | 0 | U | 1 | Y | -1 | U | 0 | 4 |
| 16. atggcgtccatgataAcat | AUG GCG UCC AUG AUA ACA UUU | 38.1 | 1 | 15,16,18, | 4 | 44 | 1 | U | 0 | G | 0 | A | 0 | N | 0 | A | 0 | 5 |
| 17. gatggcgtccatgataAca | GAU GGC GUC CAU GAU AAC AUU | 47.4 | 1 | 15,16,17, | 4 | 47 | 1 | A | 1 | G | 0 | C | 0 | N | 0 | G | -1 | 5 |
| 18. cgatggcgtccatgataA | CGA UGG CGU CCA UGA UAA CUU | 52.6 | 1 | 15,16,17, | 4 | 47 | 1 | C | 0 | U | 0 | C | 0 | Y | -1 | U | 0 | 4 |
| 19. gcgatggcgtccatgata | GCG AUG GCG UCC AUG AUA AUU | 52.6 | 1 | 16,17,18, | 4 | 47 | 1 | A | 1 | A | 1 | U | 1 | N | 0 | A | 0 | 8 |

| Criterion | Rule | Scoring |
|---|---|---|
| (GC%) | Optimal GC between 36–53% | 36–53% = 1; otherwise 0 |
| (#A/U 15–19) | Favor A/U-rich seed flank | 1 point per A/U (positions 15–19), max 5 |
| I (Tm) | Avoid strong internal repeats | Tm ≤20 °C = +2; 21–59 °C = +1; ≥60 °C = 0 |
| √ (A19) | Prefer A at position 19 | A at 19 = +1 |
| (A3) | Prefer A at position 3 | A at 3 = +1 |
| I (U10) | Prefer U at position 10 | U at 10 = +1 |
| II (not G/C19) | Penalize G/C at 19 | G or C at 19 = −1 |
| III (not G13) | Penalize G at 13 | G at 13 = −1 |

**Homology BLAST: shRNA Sequences**

shRNA: {si10-ldb3a165V}
Target Sequence: ACTCACAGGATGTTATCATGGA

**Transcripts**

| | | | | | | | | |
|---|---|---|---|---|---|---|---|---|
| PREDICTED: Mus musculus LIM domain binding 3 (Ldb3), transcript variant X4, mRNA | Mus musculus | 36.2 | 36.2 | 100% | 0.21 | 95.45% | 1497 | XM_036519019.5 |
| PREDICTED: Mus musculus LIM domain binding 3 (Ldb3), transcript variant X3, mRNA | Mus musculus | 36.2 | 36.2 | 100% | 0.21 | 95.45% | 4728 | XM_036519018.5 |
| PREDICTED: Mus musculus LIM domain binding 3 (Ldb3), transcript variant X2, mRNA | Mus musculus | 36.2 | 36.2 | 100% | 0.21 | 95.45% | 4929 | XM_036519017.5 |
| Mus musculus LIM domain binding 3 (Ldb3), transcript variant 6, mRNA | Mus musculus | 36.2 | 36.2 | 100% | 0.21 | 95.45% | 1506 | NM_001039075.2 |
| Mus musculus LIM domain binding 3 (Ldb3), transcript variant 5, mRNA | Mus musculus | 36.2 | 36.2 | 100% | 0.21 | 95.45% | 4954 | NM_001039074.2 |
| Mus musculus LIM domain binding 3 (Ldb3), transcript variant 4, mRNA | Mus musculus | 36.2 | 36.2 | 100% | 0.21 | 95.45% | 4737 | NM_001039073.2 |
| Mus musculus LIM domain binding 3 (Ldb3), transcript variant 3, mRNA | Mus musculus | 36.2 | 36.2 | 100% | 0.21 | 95.45% | 4908 | NM_001039072.2 |
| Mus musculus strain C57BL/6J chromosome 14, GRCm39 | Mus musculus | 36.2 | 4722 | 100% | 0.21 | 95.45% | 125139656 | NC_000080.7 |
| Mus musculus strain C57BL/6J chromosome 7, GRCm39 | Mus musculus | 34.2 | 4370 | 100% | 0.82 | 100.00% | 144995196 | NC_000073.7 |
| Mus musculus strain C57BL/6J chromosome 10, GRCm39 | Mus musculus | 34.2 | 3980 | 100% | 0.82 | 95.24% | 130530862 | NC_000076.7 |
| Mus musculus strain C57BL/6J chromosome 17, GRCm39 | Mus musculus | 34.2 | 3630 | 100% | 0.82 | 100.00% | 95294699 | NC_000083.7 |
| Mus musculus strain C57BL/6J chromosome 3, GRCm39 | Mus musculus | 32.2 | 5337 | 100% | 3.3 | 100.00% | 159745316 | NC_000069.7 |
| Mus musculus strain C57BL/6J chromosome 4, GRCm39 | Mus musculus | 32.2 | 5355 | 100% | 3.3 | 100.00% | 156860686 | NC_000070.7 |
| Mus musculus strain C57BL/6J chromosome 9, GRCm39 | Mus musculus | 32.2 | 4906 | 100% | 3.3 | 100.00% | 124359700 | NC_000075.7 |
| Mus musculus strain C57BL/6J chromosome X, GRCm39 | Mus musculus | 32.2 | 5091 | 100% | 3.3 | 95.00% | 169476592 | NC_000086.8 |
| PREDICTED: Mus musculus predicted gene, 38468 (Gm38468), transcript variant X2, ncRNA | Mus musculus | 30.2 | 30.2 | 86% | 13 | 94.74% | 1245 | XR_375892.4 |
| PREDICTED: Mus musculus predicted gene, 38468 (Gm38468), transcript variant X1, ncRNA | Mus musculus | 30.2 | 30.2 | 86% | 13 | 94.74% | 1509 | XR_004941619.1 |
| Mus musculus strain C57BL/6J chromosome 1, GRCm39 | Mus musculus | 30.2 | 7323 | 100% | 13 | 100.00% | 195154279 | NC_000067.7 |
| Mus musculus strain C57BL/6J chromosome 2, GRCm39 | Mus musculus | 30.2 | 7102 | 100% | 13 | 100.00% | 181755017 | NC_000068.8 |
| Mus musculus strain C57BL/6J chromosome 5, GRCm39 | Mus musculus | 30.2 | 5851 | 100% | 13 | 94.74% | 151758149 | NC_000071.7 |
| Mus musculus strain C57BL/6J chromosome 6, GRCm39 | Mus musculus | 30.2 | 5415 | 100% | 13 | 100.00% | 149588044 | NC_000072.7 |
| Mus musculus strain C57BL/6J chromosome 8, GRCm39 | Mus musculus | 30.2 | 5002 | 100% | 13 | 100.00% | 130127694 | NC_000074.7 |
| Mus musculus strain C57BL/6J chromosome 11, GRCm39 | Mus musculus | 30.2 | 4329 | 100% | 13 | 100.00% | 121973369 | NC_000077.7 |
| Mus musculus strain C57BL/6J chromosome 12, GRCm39 | Mus musculus | 30.2 | 4608 | 100% | 13 | 100.00% | 120092757 | NC_000078.7 |
| Mus musculus strain C57BL/6J chromosome 16, GRCm39 | Mus musculus | 30.2 | 3598 | 100% | 13 | 94.74% | 98008968 | NC_000082.7 |
| Mus musculus strain C57BL/6J chromosome 18, GRCm39 | Mus musculus | 30.2 | 3998 | 100% | 13 | 100.00% | 90720763 | NC_000084.7 |
| PREDICTED: Mus musculus thyroid hormone receptor interactor 11 (Trip11), transcript variant X4, mRNA | Mus musculus | 28.2 | 28.2 | 63% | 51 | 100.00% | 11764 | XM_036157159.1 |
| PREDICTED: Mus musculus thyroid hormone receptor interactor 11 (Trip11), transcript variant X3, misc_... | Mus musculus | 28.2 | 28.2 | 63% | 51 | 100.00% | 9708 | XR_004937402.1 |
| PREDICTED: Mus musculus thyroid hormone receptor interactor 11 (Trip11), transcript variant X2, mRNA | Mus musculus | 28.2 | 28.2 | 63% | 51 | 100.00% | 9683 | XM_011243986.3 |
| PREDICTED: Mus musculus predicted gene, 35032 (Gm35032), ncRNA | Mus musculus | 28.2 | 28.2 | 63% | 51 | 100.00% | 49623 | XR_380957.6 |
| PREDICTED: Mus musculus thyroid hormone receptor interactor 11 (Trip11), transcript variant X1, mRNA | Mus musculus | 28.2 | 28.2 | 63% | 51 | 100.00% | 9760 | XM_036515382.4 |

| | | | | | | | | |
|---|---|---|---|---|---|---|---|---|
| PREDICTED: Mus musculus predicted gene, 19802 (Gm19802), mRNA | Mus musculus | 28.2 | 28.2 | 63% | 51 | 100.00% | 3490 | XM_030245429.1 |
| PREDICTED: Mus musculus predicted gene, 51792 (Gm51792), mRNA | Mus musculus | 28.2 | 28.2 | 63% | 51 | 100.00% | 3514 | XM_030245427.1 |
| Mus musculus coiled-coil domain containing 97 (Ccdc97), transcript variant 6, non-coding RNA | Mus musculus | 28.2 | 28.2 | 81% | 51 | 94.44% | 2996 | NR_150796.1 |
| Mus musculus coiled-coil domain containing 97 (Ccdc97), transcript variant 3, mRNA | Mus musculus | 28.2 | 28.2 | 81% | 51 | 94.44% | 3119 | NM_001368382.1 |
| Mus musculus coiled-coil domain containing 97 (Ccdc97), transcript variant 2, mRNA | Mus musculus | 28.2 | 28.2 | 81% | 51 | 94.44% | 3010 | NM_001368381.1 |
| Mus musculus coiled-coil domain containing 97 (Ccdc97), transcript variant 4, mRNA | Mus musculus | 28.2 | 28.2 | 81% | 51 | 94.44% | 3264 | NM_001368383.1 |
| Mus musculus coiled-coil domain containing 97 (Ccdc97), transcript variant 5, mRNA | Mus musculus | 28.2 | 28.2 | 81% | 51 | 94.44% | 3108 | NM_001368384.1 |
| Mus musculus coiled-coil domain containing 97 (Ccdc97), transcript variant 1, mRNA | Mus musculus | 28.2 | 28.2 | 81% | 51 | 94.44% | 2973 | NM_028771.3 |
| Mus musculus ATPase, H+ transporting, lysosomal V1 subunit B2 (Atp6v1b2), mRNA | Mus musculus | 28.2 | 28.2 | 63% | 51 | 100.00% | 2742 | NM_007509.3 |
| Mus musculus thyroid hormone receptor interactor 11 (Trip11), mRNA | Mus musculus | 28.2 | 28.2 | 63% | 51 | 100.00% | 6448 | NM_028446.1 |
| Mus musculus tetratricopeptide repeat domain 30A1 (Ttc30a1), mRNA | Mus musculus | 28.2 | 28.2 | 63% | 51 | 100.00% | 2862 | NM_030188.3 |
| Mus musculus tetratricopeptide repeat domain 30B (Ttc30b), mRNA | Mus musculus | 28.2 | 28.2 | 63% | 51 | 100.00% | 2634 | NM_028236.1 |
| Mus musculus tetratricopeptide repeat domain 30A2 (Ttc30a2), mRNA | Mus musculus | 28.2 | 28.2 | 63% | 51 | 100.00% | 2008 | NM_001081228.1 |
| Mus musculus ras homolog family member J (Rhoj), mRNA | Mus musculus | 28.2 | 28.2 | 63% | 51 | 100.00% | 2358 | NM_023275.2 |
| Mus musculus strain C57BL/6J chromosome 13, GRCm39 | Mus musculus | 28.2 | 4603 | 100% | 51 | 100.00% | 120883175 | NC_000079.7 |
| Mus musculus strain C57BL/6J chromosome 15, GRCm39 | Mus musculus | 28.2 | 3343 | 100% | 51 | 100.00% | 104073951 | NC_000081.7 |
| Mus musculus strain C57BL/6J chromosome 19, GRCm39 | Mus musculus | 28.2 | 2368 | 100% | 51 | 100.00% | 61420004 | NC_000085.7 |
| Mus musculus strain C57BL/6J chromosome Y, GRCm39 | Mus musculus | 28.2 | 3922 | 100% | 51 | 100.00% | 91455967 | NC_000087.8 |
| PREDICTED: Mus musculus early B cell factor 3 (Ebf3), transcript variant X28, mRNA | Mus musculus | 26.3 | 26.3 | 59% | 201 | 100.00% | 4673 | XM_036152653.1 |
| PREDICTED: Mus musculus early B cell factor 3 (Ebf3), transcript variant X26, mRNA | Mus musculus | 26.3 | 26.3 | 59% | 201 | 100.00% | 5874 | XM_036152651.1 |
| PREDICTED: Mus musculus early B cell factor 3 (Ebf3), transcript variant X25, mRNA | Mus musculus | 26.3 | 26.3 | 59% | 201 | 100.00% | 5901 | XM_036152650.1 |
| PREDICTED: Mus musculus predicted gene, 52733 (Gm52733), transcript variant X2, ncRNA | Mus musculus | 26.3 | 26.3 | 77% | 201 | 94.12% | 869 | XR_004942033.1 |
| PREDICTED: Mus musculus period circadian clock 3 (Per3), transcript variant X17, misc_RNA | Mus musculus | 26.3 | 26.3 | 59% | 201 | 100.00% | 5923 | XR_004941849.1 |
| PREDICTED: Mus musculus period circadian clock 3 (Per3), transcript variant X16, misc_RNA | Mus musculus | 26.3 | 26.3 | 59% | 201 | 100.00% | 8191 | XR_004941848.1 |
| PREDICTED: Mus musculus period circadian clock 3 (Per3), transcript variant X14, misc_RNA | Mus musculus | 26.3 | 26.3 | 59% | 201 | 100.00% | 5926 | XR_004941846.1 |
| PREDICTED: Mus musculus period circadian clock 3 (Per3), transcript variant X6, mRNA | Mus musculus | 26.3 | 26.3 | 59% | 201 | 100.00% | 6036 | XM_036153826.1 |
| PREDICTED: Mus musculus period circadian clock 3 (Per3), transcript variant X5, mRNA | Mus musculus | 26.3 | 26.3 | 59% | 201 | 100.00% | 6039 | XM_011250283.4 |
| PREDICTED: Mus musculus Rho GTPase activating protein 21 (Arhgap21), transcript variant X20, mRNA | Mus musculus | 26.3 | 26.3 | 59% | 201 | 100.00% | 18104 | XM_017319263.3 |
| PREDICTED: Mus musculus Rho GTPase activating protein 21 (Arhgap21), transcript variant X19, misc_... | Mus musculus | 26.3 | 26.3 | 59% | 201 | 100.00% | 22243 | XR_004940733.1 |
| PREDICTED: Mus musculus Rho GTPase activating protein 21 (Arhgap21), transcript variant X18, misc_... | Mus musculus | 26.3 | 26.3 | 59% | 201 | 100.00% | 21126 | XR_373946.5 |
| PREDICTED: Mus musculus Rho GTPase activating protein 21 (Arhgap21), transcript variant X17, misc_... | Mus musculus | 26.3 | 26.3 | 59% | 201 | 100.00% | 21129 | XR_373945.5 |
| PREDICTED: Mus musculus Rho GTPase activating protein 21 (Arhgap21), transcript variant X15, misc_... | Mus musculus | 26.3 | 26.3 | 59% | 201 | 100.00% | 22017 | XR_004940732.1 |

| | | | | | | | | |
|---|---|---|---|---|---|---|---|---|
| PREDICTED: Mus musculus Rho GTPase activating protein 21 (Arhgap21), transcript variant X12, mRNA | Mus musculus | 26.3 | 26.3 | 59% | 201 | 100.00% | 7396 | XM_006497545.5 |
| PREDICTED: Mus musculus Rho GTPase activating protein 21 (Arhgap21), transcript variant X11, mRNA | Mus musculus | 26.3 | 26.3 | 59% | 201 | 100.00% | 20821 | XM_030252086.2 |
| PREDICTED: Mus musculus Rho GTPase activating protein 21 (Arhgap21), transcript variant X10, mRNA | Mus musculus | 26.3 | 26.3 | 59% | 201 | 100.00% | 7088 | XM_030252085.2 |
| PREDICTED: Mus musculus Rho GTPase activating protein 21 (Arhgap21), transcript variant X9, mRNA | Mus musculus | 26.3 | 26.3 | 59% | 201 | 100.00% | 7305 | XM_030252084.2 |
| PREDICTED: Mus musculus Rho GTPase activating protein 21 (Arhgap21), transcript variant X8, mRNA | Mus musculus | 26.3 | 26.3 | 59% | 201 | 100.00% | 20824 | XM_006497544.5 |
| PREDICTED: Mus musculus Rho GTPase activating protein 21 (Arhgap21), transcript variant X7, mRNA | Mus musculus | 26.3 | 26.3 | 59% | 201 | 100.00% | 7200 | XM_036162572.1 |
| PREDICTED: Mus musculus Rho GTPase activating protein 21 (Arhgap21), transcript variant X6, mRNA | Mus musculus | 26.3 | 26.3 | 59% | 201 | 100.00% | 7115 | XM_006497543.5 |
| PREDICTED: Mus musculus Rho GTPase activating protein 21 (Arhgap21), transcript variant X4, mRNA | Mus musculus | 26.3 | 26.3 | 59% | 201 | 100.00% | 20851 | XM_006497542.5 |
| PREDICTED: Mus musculus Rho GTPase activating protein 21 (Arhgap21), transcript variant X3, mRNA | Mus musculus | 26.3 | 26.3 | 59% | 201 | 100.00% | 7211 | XM_006497540.5 |
| PREDICTED: Mus musculus Rho GTPase activating protein 21 (Arhgap21), transcript variant X2, mRNA | Mus musculus | 26.3 | 26.3 | 59% | 201 | 100.00% | 7122 | XM_006497539.5 |
| PREDICTED: Mus musculus Rho GTPase activating protein 21 (Arhgap21), transcript variant X1, mRNA | Mus musculus | 26.3 | 26.3 | 59% | 201 | 100.00% | 20854 | XM_006497538.5 |
| PREDICTED: Mus musculus protein kinase, cGMP-dependent, type I (Prkg1), transcript variant X1, mRNA | Mus musculus | 26.3 | 26.3 | 77% | 201 | 94.12% | 6396 | XM_006525768.5 |
| PREDICTED: Mus musculus predicted gene, 31439 (Gm31439), transcript variant X2, ncRNA | Mus musculus | 26.3 | 26.3 | 77% | 201 | 94.12% | 7356 | XR_004939945.1 |
| PREDICTED: Mus musculus zinc finger protein 40 (Zfp40), transcript variant X3, mRNA | Mus musculus | 26.3 | 26.3 | 59% | 201 | 100.00% | 11684 | XM_030249753.2 |
| PREDICTED: Mus musculus N-acetylglucosamine-1-phosphotransferase, gamma subunit (Gnptg), trans... | Mus musculus | 26.3 | 26.3 | 59% | 201 | 100.00% | 2050 | XM_036163463.1 |
| PREDICTED: Mus musculus N-acetylglucosamine-1-phosphotransferase, gamma subunit (Gnptg), trans... | Mus musculus | 26.3 | 26.3 | 59% | 201 | 100.00% | 2047 | XR_004939505.1 |
| PREDICTED: Mus musculus N-acetylglucosamine-1-phosphotransferase, gamma subunit (Gnptg), trans... | Mus musculus | 26.3 | 26.3 | 59% | 201 | 100.00% | 3381 | XM_017317383.3 |
| PREDICTED: Mus musculus interleukin 1 receptor accessory protein (Il1rap), transcript variant X1, mRNA | Mus musculus | 26.3 | 26.3 | 59% | 201 | 100.00% | 9179 | XM_036159791.1 |
| PREDICTED: Mus musculus predicted gene, 41348 (Gm41348), ncRNA | Mus musculus | 26.3 | 26.3 | 59% | 201 | 100.00% | 1749 | XR_875429.3 |
| PREDICTED: Mus musculus SECIS binding protein 2 (Secisbp2), transcript variant X6, mRNA | Mus musculus | 26.3 | 26.3 | 59% | 201 | 100.00% | 3295 | XM_006518966.3 |
| PREDICTED: Mus musculus SECIS binding protein 2 (Secisbp2), transcript variant X5, misc_RNA | Mus musculus | 26.3 | 26.3 | 59% | 201 | 100.00% | 2307 | XR_003950489.2 |
| PREDICTED: Mus musculus SECIS binding protein 2 (Secisbp2), transcript variant X4, misc_RNA | Mus musculus | 26.3 | 26.3 | 59% | 201 | 100.00% | 3095 | XR_003950488.2 |
| PREDICTED: Mus musculus SECIS binding protein 2 (Secisbp2), transcript variant X3, misc_RNA | Mus musculus | 26.3 | 26.3 | 59% | 201 | 100.00% | 2706 | XR_382238.6 |
| PREDICTED: Mus musculus SECIS binding protein 2 (Secisbp2), transcript variant X2, misc_RNA | Mus musculus | 26.3 | 26.3 | 59% | 201 | 100.00% | 3343 | XR_382237.5 |
| PREDICTED: Mus musculus SECIS binding protein 2 (Secisbp2), transcript variant X1, misc_RNA | Mus musculus | 26.3 | 26.3 | 59% | 201 | 100.00% | 3602 | XR_004938074.1 |
| PREDICTED: Mus musculus transcription activation suppressor family member 2 (Tasor2), transcript vari... | Mus musculus | 26.3 | 26.3 | 59% | 201 | 100.00% | 7520 | XM_006515463.3 |
| PREDICTED: Mus musculus transcription activation suppressor family member 2 (Tasor2), transcript vari... | Mus musculus | 26.3 | 26.3 | 59% | 201 | 100.00% | 8137 | XM_006515461.5 |
| PREDICTED: Mus musculus predicted gene, 51809 (Gm51809), ncRNA | Mus musculus | 26.3 | 26.3 | 59% | 201 | 100.00% | 52659 | XR_003949603.2 |
| PREDICTED: Mus musculus XIAP associated factor 1 (Xaf1), transcript variant X5, mRNA | Mus musculus | 26.3 | 26.3 | 59% | 201 | 100.00% | 2292 | XM_036156775.1 |
| PREDICTED: Mus musculus XIAP associated factor 1 (Xaf1), transcript variant X4, mRNA | Mus musculus | 26.3 | 26.3 | 59% | 201 | 100.00% | 2304 | XM_017314514.3 |

**Genomic sequences**

| | | | | | | | | |
|---|---|---|---|---|---|---|---|---|
| Mus musculus strain C57BL/6J chromosome 1 unlocalized genomic scaffold, GRCm39 MMCHR1_RAND | Mus musculus | 28.2 | 52.5 | 68% | 51 | 100.00% | 153518 | NT_166282.1 |
| [illegible] | [illegible] | [illegible] | [illegible] | [illegible] | [illegible] | [illegible] | [illegible] | [illegible] |

shRNA: {si16-ldb3a165V}
Target Sequence: GCATGTACTCACAGGATGTTAT

**Transcripts**

| | | | | | | | | |
|---|---|---|---|---|---|---|---|---|
| PREDICTED: Mus musculus LIM domain binding 3 (Ldb3), transcript variant X4, mRNA | Mus musculus | 36.2 | 36.2 | 81% | 0.21 | 100.00% | 1497 | XM_036519019.5 |
| PREDICTED: Mus musculus LIM domain binding 3 (Ldb3), transcript variant X3, mRNA | Mus musculus | 36.2 | 36.2 | 81% | 0.21 | 100.00% | 4728 | XM_036519018.5 |
| PREDICTED: Mus musculus LIM domain binding 3 (Ldb3), transcript variant X2, mRNA | Mus musculus | 36.2 | 36.2 | 81% | 0.21 | 100.00% | 4929 | XM_036519017.5 |
| Mus musculus LIM domain binding 3 (Ldb3), transcript variant 6, mRNA | Mus musculus | 36.2 | 36.2 | 81% | 0.21 | 100.00% | 1506 | NM_001039075.2 |
| Mus musculus LIM domain binding 3 (Ldb3), transcript variant 5, mRNA | Mus musculus | 36.2 | 36.2 | 81% | 0.21 | 100.00% | 4954 | NM_001039074.2 |
| Mus musculus LIM domain binding 3 (Ldb3), transcript variant 4, mRNA | Mus musculus | 36.2 | 36.2 | 81% | 0.21 | 100.00% | 4737 | NM_001039073.2 |
| Mus musculus LIM domain binding 3 (Ldb3), transcript variant 3, mRNA | Mus musculus | 36.2 | 36.2 | 81% | 0.21 | 100.00% | 4908 | NM_001039072.2 |
| Mus musculus strain C57BL/6J chromosome 14, GRCm39 | Mus musculus | 36.2 | 6841 | 100% | 0.21 | 100.00% | 125139656 | NC_000080.7 |
| Mus musculus strain C57BL/6J chromosome X, GRCm39 | Mus musculus | 34.2 | 7059 | 100% | 0.82 | 100.00% | 169476592 | NC_000086.8 |
| Mus musculus strain C57BL/6J chromosome 5, GRCm39 | Mus musculus | 32.2 | 8039 | 100% | 3.3 | 100.00% | 151758149 | NC_000071.7 |
| Mus musculus strain C57BL/6J chromosome 6, GRCm39 | Mus musculus | 32.2 | 7475 | 100% | 3.3 | 100.00% | 149588044 | NC_000072.7 |
| Mus musculus strain C57BL/6J chromosome 7, GRCm39 | Mus musculus | 32.2 | 6902 | 100% | 3.3 | 100.00% | 144995196 | NC_000073.7 |
| Mus musculus strain C57BL/6J chromosome 8, GRCm39 | Mus musculus | 32.2 | 6483 | 100% | 3.3 | 100.00% | 130127694 | NC_000074.7 |
| Mus musculus strain C57BL/6J chromosome 9, GRCm39 | Mus musculus | 32.2 | 6681 | 100% | 3.3 | 100.00% | 124359700 | NC_000075.7 |
| Mus musculus strain C57BL/6J chromosome 2, GRCm39 | Mus musculus | 32.2 | 11379 | 100% | 3.3 | 100.00% | 181755017 | NC_000068.8 |
| Mus musculus strain C57BL/6J chromosome 3, GRCm39 | Mus musculus | 32.2 | 8110 | 100% | 3.3 | 100.00% | 159745316 | NC_000069.7 |
| Mus musculus strain C57BL/6J chromosome 4, GRCm39 | Mus musculus | 32.2 | 7161 | 100% | 3.3 | 100.00% | 156860686 | NC_000070.7 |
| PREDICTED: Mus musculus thyroid hormone receptor interactor 11 (Trip11), transcript variant X4, mRNA | Mus musculus | 30.2 | 30.2 | 68% | 13 | 100.00% | 11764 | XM_036157159.1 |
| PREDICTED: Mus musculus thyroid hormone receptor interactor 11 (Trip11), transcript variant X3, misc_... | Mus musculus | 30.2 | 30.2 | 68% | 13 | 100.00% | 9708 | XR_004937402.1 |
| PREDICTED: Mus musculus thyroid hormone receptor interactor 11 (Trip11), transcript variant X2, mRNA | Mus musculus | 30.2 | 30.2 | 68% | 13 | 100.00% | 9683 | XM_011243986.3 |
| PREDICTED: Mus musculus thyroid hormone receptor interactor 11 (Trip11), transcript variant X1, mRNA | Mus musculus | 30.2 | 30.2 | 68% | 13 | 100.00% | 9760 | XM_036515382.4 |
| Mus musculus thyroid hormone receptor interactor 11 (Trip11), mRNA | Mus musculus | 30.2 | 30.2 | 68% | 13 | 100.00% | 6448 | NM_028446.1 |
| Mus musculus strain C57BL/6J chromosome 1, GRCm39 | Mus musculus | 30.2 | 11019 | 100% | 13 | 100.00% | 195154279 | NC_000067.7 |
| Mus musculus strain C57BL/6J chromosome 10, GRCm39 | Mus musculus | 30.2 | 6216 | 100% | 13 | 100.00% | 130530862 | NC_000076.7 |
| Mus musculus strain C57BL/6J chromosome 11, GRCm39 | Mus musculus | 30.2 | 5972 | 100% | 13 | 100.00% | 121973369 | NC_000077.7 |
| Mus musculus strain C57BL/6J chromosome 15, GRCm39 | Mus musculus | 30.2 | 5326 | 100% | 13 | 100.00% | 104073951 | NC_000081.7 |
| Mus musculus strain C57BL/6J chromosome 19, GRCm39 | Mus musculus | 30.2 | 3367 | 100% | 13 | 100.00% | 61420004 | NC_000085.7 |
| Mus musculus strain C57BL/6J chromosome 12, GRCm39 | Mus musculus | 30.2 | 6115 | 100% | 13 | 100.00% | 120092757 | NC_000078.7 |
| Mus musculus strain C57BL/6J chromosome 13, GRCm39 | Mus musculus | 30.2 | 6348 | 100% | 13 | 100.00% | 120883175 | NC_000079.7 |

| | | | | | | | | |
|---|---|---|---|---|---|---|---|---|
| Mus musculus strain C57BL/6J chromosome 16, GRCm39 | Mus musculus | 30.2 | 4532 | 100% | 13 | 94.74% | 98008968 | NC_000082.7 |
| Mus musculus strain C57BL/6J chromosome 17, GRCm39 | Mus musculus | 30.2 | 5248 | 100% | 13 | 100.00% | 95294699 | NC_000083.7 |
| PREDICTED: Mus musculus casein kinase 1, gamma 1 (Csnk1g1), transcript variant X13, misc_RNA | Mus musculus | 28.2 | 28.2 | 63% | 51 | 100.00% | 10328 | XR_003947839.2 |
| PREDICTED: Mus musculus casein kinase 1, gamma 1 (Csnk1g1), transcript variant X11, mRNA | Mus musculus | 28.2 | 28.2 | 63% | 51 | 100.00% | 7201 | XM_011242709.3 |
| PREDICTED: Mus musculus casein kinase 1, gamma 1 (Csnk1g1), transcript variant X10, mRNA | Mus musculus | 28.2 | 28.2 | 63% | 51 | 100.00% | 7192 | XM_030244227.2 |
| PREDICTED: Mus musculus casein kinase 1, gamma 1 (Csnk1g1), transcript variant X9, mRNA | Mus musculus | 28.2 | 28.2 | 63% | 51 | 100.00% | 7287 | XM_011242708.3 |
| PREDICTED: Mus musculus casein kinase 1, gamma 1 (Csnk1g1), transcript variant X8, mRNA | Mus musculus | 28.2 | 28.2 | 63% | 51 | 100.00% | 7415 | XM_030244225.2 |
| PREDICTED: Mus musculus casein kinase 1, gamma 1 (Csnk1g1), transcript variant X7, mRNA | Mus musculus | 28.2 | 28.2 | 63% | 51 | 100.00% | 7479 | XM_036154824.1 |
| PREDICTED: Mus musculus casein kinase 1, gamma 1 (Csnk1g1), transcript variant X6, mRNA | Mus musculus | 28.2 | 28.2 | 63% | 51 | 100.00% | 7216 | XM_006511008.4 |
| PREDICTED: Mus musculus casein kinase 1, gamma 1 (Csnk1g1), transcript variant X12, mRNA | Mus musculus | 28.2 | 28.2 | 63% | 51 | 100.00% | 7177 | XM_011242710.2 |
| PREDICTED: Mus musculus casein kinase 1, gamma 1 (Csnk1g1), transcript variant X5, mRNA | Mus musculus | 28.2 | 28.2 | 63% | 51 | 100.00% | 7263 | XM_011242707.2 |
| PREDICTED: Mus musculus casein kinase 1, gamma 1 (Csnk1g1), transcript variant X4, mRNA | Mus musculus | 28.2 | 28.2 | 63% | 51 | 100.00% | 7365 | XM_011242706.2 |
| PREDICTED: Mus musculus casein kinase 1, gamma 1 (Csnk1g1), transcript variant X3, mRNA | Mus musculus | 28.2 | 28.2 | 63% | 51 | 100.00% | 7247 | XM_011242705.2 |
| PREDICTED: Mus musculus casein kinase 1, gamma 1 (Csnk1g1), transcript variant X2, mRNA | Mus musculus | 28.2 | 28.2 | 63% | 51 | 100.00% | 7430 | XM_011242704.2 |
| PREDICTED: Mus musculus casein kinase 1, gamma 1 (Csnk1g1), transcript variant X1, mRNA | Mus musculus | 28.2 | 28.2 | 63% | 51 | 100.00% | 7312 | XM_011242703.2 |
| PREDICTED: Mus musculus predicted gene, 32342 (Gm32342), transcript variant X3, ncRNA | Mus musculus | 28.2 | 28.2 | 63% | 51 | 100.00% | 4690 | XR_003952929.1 |
| PREDICTED: Mus musculus predicted gene, 30837 (Gm30837), transcript variant X2, ncRNA | Mus musculus | 28.2 | 28.2 | 63% | 51 | 100.00% | 457 | XR_376965.2 |
| PREDICTED: Mus musculus predicted gene, 30837 (Gm30837), transcript variant X1, ncRNA | Mus musculus | 28.2 | 28.2 | 63% | 51 | 100.00% | 483 | XR_376966.2 |
| Mus musculus casein kinase 1, gamma 1 (Csnk1g1), mRNA | Mus musculus | 28.2 | 28.2 | 63% | 51 | 100.00% | 7183 | NM_173185.2 |
| Mus musculus strain C57BL/6J chromosome 18, GRCm39 | Mus musculus | 28.2 | 4354 | 100% | 51 | 100.00% | 90720763 | NC_000084.7 |
| PREDICTED: Mus musculus predicted gene, 38468 (Gm38468), transcript variant X2, ncRNA | Mus musculus | 28.2 | 28.2 | 63% | 51 | 100.00% | 1245 | XR_375892.4 |
| PREDICTED: Mus musculus predicted gene, 38468 (Gm38468), transcript variant X1, ncRNA | Mus musculus | 28.2 | 28.2 | 63% | 51 | 100.00% | 1509 | XR_004941619.1 |
| PREDICTED: Mus musculus predicted gene, 30986 (Gm30986), transcript variant X2, ncRNA | Mus musculus | 28.2 | 28.2 | 63% | 51 | 100.00% | 1573 | XR_374993.4 |
| PREDICTED: Mus musculus predicted gene, 30986 (Gm30986), transcript variant X1, ncRNA | Mus musculus | 28.2 | 28.2 | 63% | 51 | 100.00% | 1793 | XR_374992.3 |
| PREDICTED: Mus musculus predicted gene 12462 (Gm12462), transcript variant X2, ncRNA | Mus musculus | 28.2 | 28.2 | 81% | 51 | 94.44% | 2864 | XR_003955388.1 |
| PREDICTED: Mus musculus predicted gene 12462 (Gm12462), transcript variant X1, ncRNA | Mus musculus | 28.2 | 28.2 | 81% | 51 | 94.44% | 3166 | XR_003955387.1 |
| Mus musculus ATPase, H+ transporting, lysosomal V1 subunit B2 (Atp6v1b2), mRNA | Mus musculus | 28.2 | 28.2 | 63% | 51 | 100.00% | 2742 | NM_007509.3 |
| Mus musculus ras homolog family member J (Rhoj), mRNA | Mus musculus | 28.2 | 28.2 | 63% | 51 | 100.00% | 2358 | NM_023275.2 |
| PREDICTED: Mus musculus X-linked Kx blood group related, X-linked (Xkrx), transcript variant X6, mRNA | Mus musculus | 26.3 | 26.3 | 59% | 201 | 100.00% | 2962 | XM_036161993.1 |
| PREDICTED: Mus musculus X-linked Kx blood group related, X-linked (Xkrx), transcript variant X5, mRNA | Mus musculus | 26.3 | 26.3 | 59% | 201 | 100.00% | 4560 | XM_006528570.5 |
| PREDICTED: Mus musculus X-linked Kx blood group related, X-linked (Xkrx), transcript variant X4, mRNA | Mus musculus | 26.3 | 26.3 | 59% | 201 | 100.00% | 4101 | XM_036161992.1 |

| | | | | | | | | |
|---|---|---|---|---|---|---|---|---|
| PREDICTED: Mus musculus X-linked Kx blood group related, X-linked (Xkrx), transcript variant X3, mRNA | Mus musculus | 26.3 | 26.3 | 59% | 201 | 100.00% | 3227 | XM_006528569.5 |
| PREDICTED: Mus musculus X-linked Kx blood group related, X-linked (Xkrx), transcript variant X2, mRNA | Mus musculus | 26.3 | 26.3 | 59% | 201 | 100.00% | 3293 | XM_017318525.3 |
| PREDICTED: Mus musculus X-linked Kx blood group related, X-linked (Xkrx), transcript variant X1, mRNA | Mus musculus | 26.3 | 26.3 | 59% | 201 | 100.00% | 2975 | XM_006528568.5 |
| PREDICTED: Mus musculus predicted gene, 26614 (Gm26614), transcript variant X3, ncRNA | Mus musculus | 26.3 | 26.3 | 59% | 201 | 100.00% | 18638 | XR_004935702.1 |
| PREDICTED: Mus musculus predicted gene, 26614 (Gm26614), transcript variant X2, ncRNA | Mus musculus | 26.3 | 26.3 | 59% | 201 | 100.00% | 17622 | XR_004935701.1 |
| PREDICTED: Mus musculus predicted gene, 26614 (Gm26614), transcript variant X1, ncRNA | Mus musculus | 26.3 | 26.3 | 59% | 201 | 100.00% | 19736 | XR_004935700.1 |
| PREDICTED: Mus musculus Rho guanine nucleotide exchange factor (GEF) 12 (Arhgef12), transcript va... | Mus musculus | 26.3 | 26.3 | 59% | 201 | 100.00% | 18790 | XM_011242598.4 |
| PREDICTED: Mus musculus Rho guanine nucleotide exchange factor (GEF) 12 (Arhgef12), transcript va... | Mus musculus | 26.3 | 26.3 | 59% | 201 | 100.00% | 18783 | XM_030244573.2 |
| PREDICTED: Mus musculus Rho guanine nucleotide exchange factor (GEF) 12 (Arhgef12), transcript va... | Mus musculus | 26.3 | 26.3 | 59% | 201 | 100.00% | 18840 | XM_030244572.2 |
| PREDICTED: Mus musculus Rho guanine nucleotide exchange factor (GEF) 12 (Arhgef12), transcript va... | Mus musculus | 26.3 | 26.3 | 59% | 201 | 100.00% | 18786 | XM_017313567.3 |
| PREDICTED: Mus musculus Rho guanine nucleotide exchange factor (GEF) 12 (Arhgef12), transcript va... | Mus musculus | 26.3 | 26.3 | 59% | 201 | 100.00% | 18843 | XM_030244571.2 |
| PREDICTED: Mus musculus uncharacterized LOC118567615 (LOC118567615), ncRNA | Mus musculus | 26.3 | 26.3 | 59% | 201 | 100.00% | 647 | XR_004935080.1 |
| PREDICTED: Mus musculus zinc finger protein 930 (Zfp930), transcript variant X3, mRNA | Mus musculus | 26.3 | 26.3 | 59% | 201 | 100.00% | 3079 | XM_030243456.2 |
| PREDICTED: Mus musculus zinc finger protein 930 (Zfp930), transcript variant X2, mRNA | Mus musculus | 26.3 | 26.3 | 59% | 201 | 100.00% | 3865 | XM_030243455.2 |
| PREDICTED: Mus musculus zinc finger protein 930 (Zfp930), transcript variant X1, mRNA | Mus musculus | 26.3 | 26.3 | 59% | 201 | 100.00% | 3049 | XM_030243454.2 |
| PREDICTED: Mus musculus early B cell factor 3 (Ebf3), transcript variant X28, mRNA | Mus musculus | 26.3 | 26.3 | 59% | 201 | 100.00% | 4673 | XM_036152653.1 |
| PREDICTED: Mus musculus early B cell factor 3 (Ebf3), transcript variant X26, mRNA | Mus musculus | 26.3 | 26.3 | 59% | 201 | 100.00% | 5874 | XM_036152651.1 |
| PREDICTED: Mus musculus early B cell factor 3 (Ebf3), transcript variant X25, mRNA | Mus musculus | 26.3 | 26.3 | 59% | 201 | 100.00% | 5901 | XM_036152650.1 |
| PREDICTED: Mus musculus zinc finger protein 638 (Zfp638), transcript variant X7, misc_RNA | Mus musculus | 26.3 | 26.3 | 77% | 201 | 94.12% | 7505 | XR_004942919.1 |
| PREDICTED: Mus musculus mitochondrial transcription termination factor 1a (Mterf1a), transcript variant... | Mus musculus | 26.3 | 26.3 | 77% | 201 | 94.12% | 3596 | XM_017320999.3 |
| PREDICTED: Mus musculus mitochondrial transcription termination factor 1a (Mterf1a), transcript variant... | Mus musculus | 26.3 | 26.3 | 77% | 201 | 94.12% | 3628 | XM_017321000.2 |
| PREDICTED: Mus musculus predicted gene, 30019 (Gm30019), transcript variant X2, ncRNA | Mus musculus | 26.3 | 26.3 | 59% | 201 | 100.00% | 1532 | XR_003955285.2 |
| PREDICTED: Mus musculus Rho GTPase activating protein 21 (Arhgap21), transcript variant X20, mRNA | Mus musculus | 26.3 | 26.3 | 59% | 201 | 100.00% | 18104 | XM_017319263.3 |
| PREDICTED: Mus musculus Rho GTPase activating protein 21 (Arhgap21), transcript variant X19, misc_... | Mus musculus | 26.3 | 26.3 | 59% | 201 | 100.00% | 22243 | XR_004940733.1 |
| PREDICTED: Mus musculus Rho GTPase activating protein 21 (Arhgap21), transcript variant X18, misc_... | Mus musculus | 26.3 | 26.3 | 59% | 201 | 100.00% | 21126 | XR_373946.5 |
| PREDICTED: Mus musculus Rho GTPase activating protein 21 (Arhgap21), transcript variant X17, misc_... | Mus musculus | 26.3 | 26.3 | 59% | 201 | 100.00% | 21129 | XR_373945.5 |
| PREDICTED: Mus musculus Rho GTPase activating protein 21 (Arhgap21), transcript variant X15, misc_... | Mus musculus | 26.3 | 26.3 | 59% | 201 | 100.00% | 22017 | XR_004940732.1 |
| PREDICTED: Mus musculus Rho GTPase activating protein 21 (Arhgap21), transcript variant X15, mRNA | Mus musculus | 26.3 | 26.3 | 59% | 201 | 100.00% | 21263 | XM_017319262.3 |
| PREDICTED: Mus musculus Rho GTPase activating protein 21 (Arhgap21), transcript variant X14, mRNA | Mus musculus | 26.3 | 26.3 | 59% | 201 | 100.00% | 21266 | XM_011238986.4 |
| PREDICTED: Mus musculus Rho GTPase activating protein 21 (Arhgap21), transcript variant X11, mRNA | Mus musculus | 26.3 | 26.3 | 59% | 201 | 100.00% | 20821 | XM_030252086.2 |
| PREDICTED: Mus musculus Rho GTPase activating protein 21 (Arhgap21), transcript variant X8, mRNA | Mus musculus | 26.3 | 26.3 | 59% | 201 | 100.00% | 20824 | XM_006497544.5 |

| | | | | | | | | |
|---|---|---|---|---|---|---|---|---|
| PREDICTED: Mus musculus Rho GTPase activating protein 21 (Arhgap21), transcript variant X4, mRNA | Mus musculus | 26.3 | 26.3 | 59% | 201 | 100.00% | 20851 | XM_006497542.5 |
| PREDICTED: Mus musculus Rho GTPase activating protein 21 (Arhgap21), transcript variant X1, mRNA | Mus musculus | 26.3 | 26.3 | 59% | 201 | 100.00% | 20854 | XM_006497538.5 |
| PREDICTED: Mus musculus N-acetylglucosamine-1-phosphotransferase, gamma subunit (Gnptg), trans... | Mus musculus | 26.3 | 26.3 | 59% | 201 | 100.00% | 2050 | XM_036163463.1 |
| PREDICTED: Mus musculus N-acetylglucosamine-1-phosphotransferase, gamma subunit (Gnptg), trans... | Mus musculus | 26.3 | 26.3 | 59% | 201 | 100.00% | 2047 | XR_004939505.1 |
| PREDICTED: Mus musculus N-acetylglucosamine-1-phosphotransferase, gamma subunit (Gnptg), trans... | Mus musculus | 26.3 | 26.3 | 59% | 201 | 100.00% | 3381 | XM_017317383.3 |
| PREDICTED: Mus musculus interleukin 1 receptor accessory protein (Il1rap), transcript variant X1, mRNA | Mus musculus | 26.3 | 26.3 | 59% | 201 | 100.00% | 9179 | XM_036159791.1 |
| PREDICTED: Mus musculus predicted gene, 41348 (Gm41348), ncRNA | Mus musculus | 26.3 | 26.3 | 59% | 201 | 100.00% | 1749 | XR_875429.3 |
| PREDICTED: Mus musculus myosin X (Myo10), transcript variant X1, mRNA | Mus musculus | 26.3 | 26.3 | 77% | 201 | 94.12% | 14651 | XM_030248368.2 |
| PREDICTED: Mus musculus predicted gene, 52128 (Gm52128), ncRNA | Mus musculus | 26.3 | 26.3 | 77% | 201 | 94.12% | 36006 | XR_003951168.2 |

shRNA: Scramble miR30-shRNA
Target Sequence: ACCTAAGGTTAAGTCGCCCTCG

**Transcripts**

| | | | | | | | | |
|---|---|---|---|---|---|---|---|---|
| Mus musculus strain C57BL/6J chromosome X, GRCm39 | Mus musculus | 32.2 | 1017 | 95% | 3.3 | 100.00% | 169476592 | NC_000086.8 |
| Mus musculus strain C57BL/6J chromosome 2, GRCm39 | Mus musculus | 30.2 | 1533 | 100% | 13 | 100.00% | 181755017 | NC_000068.8 |
| Mus musculus strain C57BL/6J chromosome 14, GRCm39 | Mus musculus | 30.2 | 1015 | 100% | 13 | 100.00% | 125139656 | NC_000080.7 |
| PREDICTED: Mus musculus predicted gene 13748 (Gm13748), ncRNA | Mus musculus | 28.2 | 28.2 | 63% | 51 | 100.00% | 11052 | XR_003947836.2 |
| Mus musculus strain C57BL/6J chromosome 1, GRCm39 | Mus musculus | 28.2 | 1504 | 100% | 51 | 100.00% | 195154279 | NC_000067.7 |
| Mus musculus strain C57BL/6J chromosome 4, GRCm39 | Mus musculus | 28.2 | 1426 | 100% | 51 | 100.00% | 156860686 | NC_000070.7 |
| Mus musculus strain C57BL/6J chromosome 5, GRCm39 | Mus musculus | 28.2 | 1504 | 100% | 51 | 100.00% | 151758149 | NC_000071.7 |
| Mus musculus strain C57BL/6J chromosome 6, GRCm39 | Mus musculus | 28.2 | 1078 | 95% | 51 | 100.00% | 149588044 | NC_000072.7 |
| Mus musculus strain C57BL/6J chromosome 7, GRCm39 | Mus musculus | 28.2 | 1033 | 100% | 51 | 100.00% | 144995196 | NC_000073.7 |
| Mus musculus strain C57BL/6J chromosome 8, GRCm39 | Mus musculus | 28.2 | 1084 | 100% | 51 | 100.00% | 130127694 | NC_000074.7 |
| Mus musculus strain C57BL/6J chromosome 11, GRCm39 | Mus musculus | 28.2 | 1065 | 95% | 51 | 100.00% | 121973369 | NC_000077.7 |
| Mus musculus strain C57BL/6J chromosome 12, GRCm39 | Mus musculus | 28.2 | 1092 | 90% | 51 | 100.00% | 120092757 | NC_000078.7 |
| Mus musculus strain C57BL/6J chromosome 13, GRCm39 | Mus musculus | 28.2 | 1219 | 90% | 51 | 94.44% | 120883175 | NC_000079.7 |
| Mus musculus strain C57BL/6J chromosome 15, GRCm39 | Mus musculus | 28.2 | 693 | 95% | 51 | 100.00% | 104073951 | NC_000081.7 |
| Mus musculus strain C57BL/6J chromosome 16, GRCm39 | Mus musculus | 28.2 | 667 | 95% | 51 | 100.00% | 98008968 | NC_000082.7 |
| Mus musculus strain C57BL/6J chromosome 17, GRCm39 | Mus musculus | 28.2 | 780 | 95% | 51 | 100.00% | 95294699 | NC_000083.7 |
| Mus musculus strain C57BL/6J chromosome 19, GRCm39 | Mus musculus | 28.2 | 471 | 72% | 51 | 100.00% | 61420004 | NC_000085.7 |
| PREDICTED: Mus musculus mitoguardin 1 (Miga1), transcript variant X3, mRNA | Mus musculus | 26.3 | 26.3 | 59% | 201 | 100.00% | 13109 | XM_036163015.1 |
| PREDICTED: Mus musculus ankyrin 2, brain (Ank2), transcript variant X43, mRNA | Mus musculus | 26.3 | 26.3 | 59% | 201 | 100.00% | 32626 | XM_036162848.1 |
| PREDICTED: Mus musculus ankyrin 2, brain (Ank2), transcript variant X38, mRNA | Mus musculus | 26.3 | 26.3 | 59% | 201 | 100.00% | 32656 | XM_036162845.1 |
| PREDICTED: Mus musculus ankyrin 2, brain (Ank2), transcript variant X35, mRNA | Mus musculus | 26.3 | 26.3 | 59% | 201 | 100.00% | 32680 | XM_036162842.1 |
| PREDICTED: Mus musculus ankyrin 2, brain (Ank2), transcript variant X34, mRNA | Mus musculus | 26.3 | 26.3 | 59% | 201 | 100.00% | 32680 | XM_036162841.1 |
| PREDICTED: Mus musculus ankyrin 2, brain (Ank2), transcript variant X31, mRNA | Mus musculus | 26.3 | 26.3 | 59% | 201 | 100.00% | 32700 | XM_036162838.1 |
| PREDICTED: Mus musculus ankyrin 2, brain (Ank2), transcript variant X22, mRNA | Mus musculus | 26.3 | 26.3 | 59% | 201 | 100.00% | 32755 | XM_036162833.1 |
| PREDICTED: Mus musculus ankyrin 2, brain (Ank2), transcript variant X19, mRNA | Mus musculus | 26.3 | 26.3 | 59% | 201 | 100.00% | 32779 | XM_036162830.1 |
| PREDICTED: Mus musculus guanosine monophosphate reductase 2 (Gmpr2), transcript variant X2, mRNA | Mus musculus | 26.3 | 26.3 | 59% | 201 | 100.00% | 6265 | XM_011244882.4 |
| PREDICTED: Mus musculus predicted gene, 52013 (Gm52013), transcript variant X2, ncRNA | Mus musculus | 26.3 | 26.3 | 59% | 201 | 100.00% | 26661 | XR_004937626.1 |
| PREDICTED: Mus musculus predicted gene, 52013 (Gm52013), transcript variant X1, ncRNA | Mus musculus | 26.3 | 26.3 | 59% | 201 | 100.00% | 21529 | XR_004937625.1 |
| PREDICTED: Mus musculus mitoguardin 1 (Miga1), transcript variant X4, misc_RNA | Mus musculus | 26.3 | 26.3 | 59% | 201 | 100.00% | 5018 | XR_001783676.2 |

| | | | | | | | | |
|---|---|---|---|---|---|---|---|---|
| PREDICTED: Mus musculus predicted gene, 46219 (Gm46219), ncRNA | Mus musculus | 26.3 | 26.3 | 59% | 201 | 100.00% | 1695 | XR_001779752.2 |
| Mus musculus POU domain, class 4, transcription factor 3 (Pou4f3), mRNA | Mus musculus | 26.3 | 26.3 | 59% | 201 | 100.00% | 1146 | NM_138945.2 |
| Mus musculus mitoguardin 1 (Miga1), transcript variant 2, mRNA | Mus musculus | 26.3 | 26.3 | 59% | 201 | 100.00% | 5163 | NM_001162375.1 |
| Mus musculus mitoguardin 1 (Miga1), transcript variant 1, mRNA | Mus musculus | 26.3 | 26.3 | 59% | 201 | 100.00% | 5355 | NM_174868.4 |
| Mus musculus strain C57BL/6J chromosome 3, GRCm39 | Mus musculus | 26.3 | 1286 | 95% | 201 | 100.00% | 159745316 | NC_000069.7 |
| Mus musculus strain C57BL/6J chromosome 9, GRCm39 | Mus musculus | 26.3 | 1031 | 100% | 201 | 100.00% | 124359700 | NC_000075.7 |
| Mus musculus strain C57BL/6J chromosome 10, GRCm39 | Mus musculus | 26.3 | 1349 | 100% | 201 | 100.00% | 130530862 | NC_000076.7 |
| Mus musculus strain C57BL/6J chromosome 18, GRCm39 | Mus musculus | 26.3 | 616 | 100% | 201 | 100.00% | 90720763 | NC_000084.7 |
| PREDICTED: Mus musculus RIKEN cDNA 4931406C07 gene (4931406C07Rik), transcript variant X3, m... | Mus musculus | 24.3 | 24.3 | 54% | 794 | 100.00% | 2586 | XM_030244585.2 |
| PREDICTED: Mus musculus RIKEN cDNA 4931406C07 gene (4931406C07Rik), transcript variant X2, m... | Mus musculus | 24.3 | 24.3 | 54% | 794 | 100.00% | 2749 | XM_036155283.1 |
| PREDICTED: Mus musculus RIKEN cDNA 4931406C07 gene (4931406C07Rik), transcript variant X1, m... | Mus musculus | 24.3 | 24.3 | 54% | 794 | 100.00% | 2637 | XM_006510596.3 |
| PREDICTED: Mus musculus enhancer of mRNA decapping 3 (Edc3), transcript variant X3, mRNA | Mus musculus | 24.3 | 24.3 | 54% | 794 | 100.00% | 5644 | XM_011242760.4 |
| PREDICTED: Mus musculus enhancer of mRNA decapping 3 (Edc3), transcript variant X1, mRNA | Mus musculus | 24.3 | 24.3 | 54% | 794 | 100.00% | 4352 | XM_036155089.1 |
| PREDICTED: Mus musculus predicted gene, 32643 (Gm32643), transcript variant X15, ncRNA | Mus musculus | 24.3 | 24.3 | 54% | 794 | 100.00% | 14103 | XR_003947629.2 |
| PREDICTED: Mus musculus predicted gene, 32643 (Gm32643), transcript variant X14, ncRNA | Mus musculus | 24.3 | 24.3 | 54% | 794 | 100.00% | 14062 | XR_003947628.2 |
| PREDICTED: Mus musculus predicted gene, 32643 (Gm32643), transcript variant X13, ncRNA | Mus musculus | 24.3 | 24.3 | 54% | 794 | 100.00% | 14157 | XR_003947627.2 |
| PREDICTED: Mus musculus predicted gene, 32643 (Gm32643), transcript variant X12, ncRNA | Mus musculus | 24.3 | 24.3 | 54% | 794 | 100.00% | 13981 | XR_003947626.2 |
| PREDICTED: Mus musculus predicted gene, 32643 (Gm32643), transcript variant X11, ncRNA | Mus musculus | 24.3 | 24.3 | 54% | 794 | 100.00% | 14008 | XR_003947625.2 |
| PREDICTED: Mus musculus predicted gene, 32643 (Gm32643), transcript variant X10, ncRNA | Mus musculus | 24.3 | 24.3 | 54% | 794 | 100.00% | 13934 | XR_879390.4 |
| PREDICTED: Mus musculus predicted gene, 32643 (Gm32643), transcript variant X9, ncRNA | Mus musculus | 24.3 | 24.3 | 54% | 794 | 100.00% | 14401 | XR_004935174.1 |
| PREDICTED: Mus musculus predicted gene, 32643 (Gm32643), transcript variant X8, ncRNA | Mus musculus | 24.3 | 24.3 | 54% | 794 | 100.00% | 14251 | XR_004935173.1 |
| PREDICTED: Mus musculus predicted gene, 32643 (Gm32643), transcript variant X7, ncRNA | Mus musculus | 24.3 | 24.3 | 54% | 794 | 100.00% | 14056 | XR_879387.4 |
| PREDICTED: Mus musculus predicted gene, 32643 (Gm32643), transcript variant X6, ncRNA | Mus musculus | 24.3 | 24.3 | 54% | 794 | 100.00% | 14103 | XR_388237.5 |
| PREDICTED: Mus musculus predicted gene, 32643 (Gm32643), transcript variant X5, ncRNA | Mus musculus | 24.3 | 24.3 | 54% | 794 | 100.00% | 13953 | XR_879386.4 |
| PREDICTED: Mus musculus predicted gene, 32643 (Gm32643), transcript variant X4, ncRNA | Mus musculus | 24.3 | 24.3 | 54% | 794 | 100.00% | 14419 | XR_004935172.1 |
| PREDICTED: Mus musculus predicted gene, 32643 (Gm32643), transcript variant X3, ncRNA | Mus musculus | 24.3 | 24.3 | 54% | 794 | 100.00% | 14269 | XR_004935171.1 |
| PREDICTED: Mus musculus predicted gene, 32643 (Gm32643), transcript variant X2, ncRNA | Mus musculus | 24.3 | 24.3 | 54% | 794 | 100.00% | 13906 | XR_879384.4 |
| PREDICTED: Mus musculus predicted gene, 32643 (Gm32643), transcript variant X1, ncRNA | Mus musculus | 24.3 | 24.3 | 54% | 794 | 100.00% | 13886 | XR_879385.4 |
| PREDICTED: Mus musculus complement component 5a receptor 2 (C5ar2), transcript variant X9, mRNA | Mus musculus | 24.3 | 24.3 | 54% | 794 | 100.00% | 2185 | XM_006540056.3 |
| PREDICTED: Mus musculus complement component 5a receptor 2 (C5ar2), transcript variant X8, mRNA | Mus musculus | 24.3 | 24.3 | 54% | 794 | 100.00% | 2542 | XM_006540055.5 |
| PREDICTED: Mus musculus complement component 5a receptor 2 (C5ar2), transcript variant X7, mRNA | Mus musculus | 24.3 | 24.3 | 54% | 794 | 100.00% | 2514 | XM_006540054.5 |

| | | | | | | | | |
|---|---|---|---|---|---|---|---|---|
| PREDICTED: Mus musculus complement component 5a receptor 2 (C5ar2), transcript variant X1, misc_... | Mus musculus | 24.3 | 24.3 | 54% | 794 | 100.00% | 2192 | XR_881730.4 |
| PREDICTED: Mus musculus potassium voltage-gated channel, Shal-related family, member 2 (Kcnd2), tr... | Mus musculus | 24.3 | 24.3 | 54% | 794 | 100.00% | 95035 | XM_036165884.1 |
| PREDICTED: Mus musculus predicted gene, 52764 (Gm52764), transcript variant X1, ncRNA | Mus musculus | 24.3 | 24.3 | 54% | 794 | 100.00% | 7721 | XR_004942721.1 |
| PREDICTED: Mus musculus predicted gene, 19417 (Gm19417), ncRNA | Mus musculus | 24.3 | 24.3 | 54% | 794 | 100.00% | 26255 | XR_376437.5 |
| PREDICTED: Mus musculus predicted gene 11802 (Gm11802), transcript variant X7, ncRNA | Mus musculus | 24.3 | 24.3 | 54% | 794 | 100.00% | 10637 | XR_881093.4 |
| PREDICTED: Mus musculus predicted gene 11802 (Gm11802), transcript variant X6, ncRNA | Mus musculus | 24.3 | 24.3 | 54% | 794 | 100.00% | 10733 | XR_881092.4 |
| PREDICTED: Mus musculus predicted gene 11802 (Gm11802), transcript variant X3, ncRNA | Mus musculus | 24.3 | 24.3 | 54% | 794 | 100.00% | 10797 | XR_881089.4 |
| PREDICTED: Mus musculus predicted gene 11802 (Gm11802), transcript variant X2, ncRNA | Mus musculus | 24.3 | 24.3 | 54% | 794 | 100.00% | 10785 | XR_881088.4 |
| PREDICTED: Mus musculus predicted gene 11802 (Gm11802), transcript variant X1, ncRNA | Mus musculus | 24.3 | 24.3 | 54% | 794 | 100.00% | 10661 | XR_881090.4 |
| PREDICTED: Mus musculus alkB homolog 3, alpha-ketoglutarate-dependent dioxygenase (Alkbh3), trans... | Mus musculus | 24.3 | 24.3 | 54% | 794 | 100.00% | 11519 | XM_006500125.4 |
| PREDICTED: Mus musculus predicted gene, 30620 (Gm30620), transcript variant X6, ncRNA | Mus musculus | 24.3 | 24.3 | 54% | 794 | 100.00% | 3760 | XR_004939998.1 |
| PREDICTED: Mus musculus predicted gene, 30620 (Gm30620), transcript variant X4, ncRNA | Mus musculus | 24.3 | 24.3 | 54% | 794 | 100.00% | 4531 | XR_004939997.1 |
| PREDICTED: Mus musculus predicted gene, 30620 (Gm30620), transcript variant X1, ncRNA | Mus musculus | 24.3 | 24.3 | 54% | 794 | 100.00% | 4534 | XR_004939996.1 |
| PREDICTED: Mus musculus protocadherin 9 (Pcdh9), transcript variant X2, misc_RNA | Mus musculus | 24.3 | 24.3 | 54% | 794 | 100.00% | 148183 | XR_003950833.2 |
| PREDICTED: Mus musculus prospero homeobox 2 (Prox2), transcript variant X4, mRNA | Mus musculus | 24.3 | 24.3 | 54% | 794 | 100.00% | 3910 | XM_017315213.3 |
| PREDICTED: Mus musculus prospero homeobox 2 (Prox2), transcript variant X2, mRNA | Mus musculus | 24.3 | 24.3 | 54% | 794 | 100.00% | 8282 | XM_030246952.2 |
| PREDICTED: Mus musculus prospero homeobox 2 (Prox2), transcript variant X1, mRNA | Mus musculus | 24.3 | 24.3 | 54% | 794 | 100.00% | 4350 | XM_006516278.5 |
| PREDICTED: Mus musculus myosin phosphatase Rho interacting protein (Mprip), transcript variant X2, m... | Mus musculus | 24.3 | 24.3 | 54% | 794 | 100.00% | 16540 | XR_003949455.2 |
| PREDICTED: Mus musculus vascular endothelial zinc finger 1 (Vezf1), transcript variant X9, mRNA | Mus musculus | 24.3 | 24.3 | 54% | 794 | 100.00% | 7355 | XM_030245929.2 |
| PREDICTED: Mus musculus vascular endothelial zinc finger 1 (Vezf1), transcript variant X8, mRNA | Mus musculus | 24.3 | 24.3 | 54% | 794 | 100.00% | 6041 | XM_030245928.2 |
| PREDICTED: Mus musculus vascular endothelial zinc finger 1 (Vezf1), transcript variant X7, mRNA | Mus musculus | 24.3 | 24.3 | 54% | 794 | 100.00% | 7158 | XM_030245927.2 |
| PREDICTED: Mus musculus vascular endothelial zinc finger 1 (Vezf1), transcript variant X6, mRNA | Mus musculus | 24.3 | 24.3 | 54% | 794 | 100.00% | 7190 | XM_011248969.3 |
| PREDICTED: Mus musculus vascular endothelial zinc finger 1 (Vezf1), transcript variant X5, mRNA | Mus musculus | 24.3 | 24.3 | 54% | 794 | 100.00% | 7708 | XM_036156629.1 |
| PREDICTED: Mus musculus vascular endothelial zinc finger 1 (Vezf1), transcript variant X4, mRNA | Mus musculus | 24.3 | 24.3 | 54% | 794 | 100.00% | 7726 | XM_036156628.1 |
| PREDICTED: Mus musculus vascular endothelial zinc finger 1 (Vezf1), transcript variant X3, mRNA | Mus musculus | 24.3 | 24.3 | 54% | 794 | 100.00% | 4951 | XM_011248968.4 |
| PREDICTED: Mus musculus vascular endothelial zinc finger 1 (Vezf1), transcript variant X2, mRNA | Mus musculus | 24.3 | 24.3 | 54% | 794 | 100.00% | 5644 | XM_036156627.1 |
| PREDICTED: Mus musculus vascular endothelial zinc finger 1 (Vezf1), transcript variant X1, mRNA | Mus musculus | 24.3 | 24.3 | 54% | 794 | 100.00% | 5662 | XM_036156626.1 |
| PREDICTED: Mus musculus predicted gene, 51801 (Gm51801), transcript variant X2, ncRNA | Mus musculus | 24.3 | 24.3 | 54% | 794 | 100.00% | 7561 | XR_003949003.2 |
| PREDICTED: Mus musculus predicted gene, 51801 (Gm51801), transcript variant X1, ncRNA | Mus musculus | 24.3 | 24.3 | 54% | 794 | 100.00% | 6458 | XR_004936324.1 |
| PREDICTED: Mus musculus NUAK family, SNF1-like kinase, 1 (Nuak1), transcript variant X2, mRNA | Mus musculus | 24.3 | 24.3 | 54% | 794 | 100.00% | 5206 | XM_011243603.4 |
| PREDICTED: Mus musculus NUAK family, SNF1-like kinase, 1 (Nuak1), transcript variant X1, mRNA | Mus musculus | 24.3 | 24.3 | 54% | 794 | 100.00% | 11216 | XM_036156104.1 |

| | | | | | | | | |
|---|---|---|---|---|---|---|---|---|
| PREDICTED: Mus musculus TBC1 domain family, member 30 (Tbc1d30), transcript variant X1, mRNA | Mus musculus | 24.3 | 24.3 | 54% | 794 | 100.00% | 5447 | XM_030245334.2 |
| PREDICTED: Mus musculus LEM domain containing 3 (Lemd3), transcript variant X1, misc_RNA | Mus musculus | 24.3 | 24.3 | 54% | 794 | 100.00% | 4917 | XR_001779544.3 |
| PREDICTED: Mus musculus TAFA chemokine like family member 2 (Tafa2), transcript variant X5, mRNA | Mus musculus | 24.3 | 24.3 | 54% | 794 | 100.00% | 18141 | XM_036155794.1 |
| PREDICTED: Mus musculus TAFA chemokine like family member 2 (Tafa2), transcript variant X4, mRNA | Mus musculus | 24.3 | 24.3 | 54% | 794 | 100.00% | 17894 | XM_030245097.2 |
| PREDICTED: Mus musculus TAFA chemokine like family member 2 (Tafa2), transcript variant X3, mRNA | Mus musculus | 24.3 | 24.3 | 54% | 794 | 100.00% | 18249 | XM_036155793.1 |

| Selected shRNA and Target Sequences | | | |
|---|---|---|---|
| **shRNA** | **Target Sequence in Mouse** | **Potential Off-Target mRNA (Mice)** | **Sequence Homology (%)** |
| | | Mitoguardin 1 (Miga1) | 59 |
| shSCRNAmiR | ACCTAAGGTTAAGTCGCCCTCG | Ankyrin 2, brain (Ank2) | 59 |
| | | POU domain, class 4, transcription factor 3 (Pou4f3) | 59 |
| | | Coiled-coil domain containing 97 (Ccdc97) | 81 |
| sh10RNAmiR | ACTCACAGGATGTTATCATGGA | Thyroid hormone receptor interactor 11 (Trip11) | 63 |
| | | ATPase, H+ transporting, lysosomal V1 subunit B2 (Atp6v1b2) | 63 |
| | | Thyroid hormone receptor interactor 11 (Trip11) | 68 |
| sh16RNAmiR | GCATGTACTCACAGGATGTTAT | Casein Kinase 1, gamma 1 (Csnk1g1) | 63 |
| | | ATPase, H+ transporting, lysosomal V1 subunit B2 (Atp6v1b2) | 63 |

Therapeutic shRNAs were designed in a miR30 backbone for allele-specific silencing of the mutant *Ldb3* transcript, with in silico screening revealing only partial homology (≤68%) to unrelated mouse transcripts and no high-risk off-target matches.
A scramble shRNA with comparable homology did not produce molecular or functional effects, and two independent shRNAs yielded concordant outcomes, supporting target-specific silencing.

pAAV[miR30]-CMV>EGFP:{si10-Ldb3A165V}:WPRE

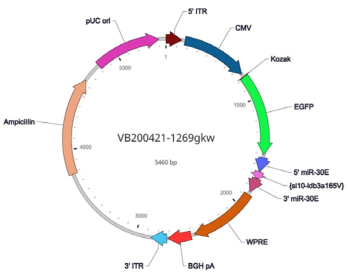

pAAV[miR30]-CMV>EGFP:{si16-ldb3a165V}:WPRE

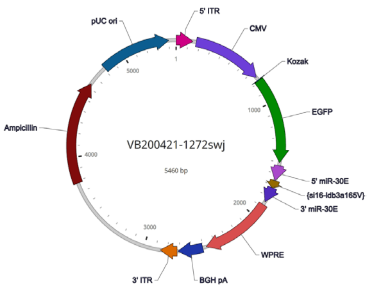

pAAV[miR30]-CMV>EGFP:Scramble_miR30-shRNA:WPRE

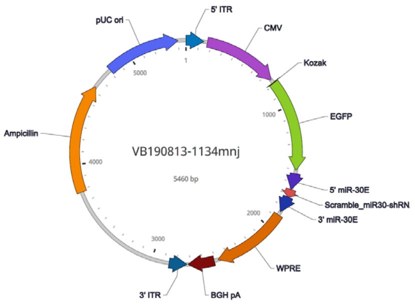

| S. No. | Protein | Modifications (all sites) |
|---|---|---|
| 1 | SLC25A4 | Y191, Y195, T197 |
| 2 | ACTN2 | S840 |
| 3 | ANKRD2 | S36 |
| 4 | CASQ1 | Y57 |
| 5 | CA3 | S48, Y51 |
| 6 | CAVIN1 | S38, T40, S42 |
| 7 | CKM | S164, T166, S372, T322, T327, S337 |
| 8 | SPAG9 | T365 |
| 9 | ALDH4A1 | T541, S542 |
| 10 | DES | S437 |
| 11 | EEF1D | S162 |
| 12 | FLNC | S2234, S2237 |
| 13 | ALDOA | S36, T37, S39 |
| 14 | PYGM | S514 |
| 15 | GYS1 | S652, S653, S657 |
| 16 | HSP90AB1 | S226, S255 |
| 17 | HSPA12B | S276 |
| 18 | JSRP1 | S223, S225, S228, S232 |
| 19 | JPH1 | T460, T461, S469, S473, S475, S480 |
| 20 | LDB3 | S98, S179 |
| 21 | ACSL1 | S423, S424 |
| 22 | LDHA | T322 |
| 23 | MARCKS | S138, S141, T143 |
| 24 | FXYD1 | S82, S83 |
| 25 | PHKB | S692, S693 |
| 26 | ATP2A1 | S547 |
| 27 | SPEG | S542, S545, T546, S2322, S2323, S2325, S2327, S2333 |
| 28 | SYNPO2L | T702 |
| 29 | SYNPO2 | S895, S899 |
| 30 | TTN | S264, T266, S269, T299, S301 |
| 31 | TPM1 | S174 |
| 32 | UBAC1 | S98 |
| 33 | VDAC1 | S115, S117 |
| 34 | VDAC2 | T114, S116 |
| 35 | CACNB1 | T418 |

| CYTO_pho | CYTO_tota | Annotated | Modificati | Confidenc | # Proteins | # PSMs | Master Pro | Positions i | Master Pro | Modificati | # Missed C | m/z [Da] ( | Theo. MH+ | DeltaM [pp | XCorr (by S | Log2_Mea | Log2_Mea | Linear_Dif | Welch_Tte | **Welch_Ttest_Corrected_Pvalue** |
|---|---|---|---|---|---|---|---|---|---|---|---|---|---|---|---|---|---|---|---|---|
| CYTO_pho | CYTO_tota | [K].DIKHD | 1xPhosph | High | 1 | 9 | P16015 | P16015 [3 | Carbonic a | P16015 1x | 1 | 823.678 | 3291.69 | 0.54 | 6.53 | 0.00388 | 0.00888 | 2.28671 | 2.94E-05 | 0.00642 |
| CYTO_pho | CYTO_tota | [K].LEKGQ | 1xPhosph | High | 1 | 8 | P07310 | P07310 [3 | Creatine k | P07310 1x | 1 | 894.485 | 2681.44 | 0.3 | 4.34 | 0.00085 | 0.00138 | 1.63628 | 3.46E-05 | 0.00642 |
| CYTO_pho | CYTO_tota | [K].AEDGA | 1xPhosph | High | 1 | 10 | P26645 | P26645 [1 | Myristoyla | P26645 1x | 0 | 711.351 | 2132.04 | -0.45 | 5.43 | 0.3704 | 0.7839 | 2.11639 | 3.90E-05 | 0.00642 |
| CYTO_pho | CYTO_tota | [K].ATHPP | 1xPhosph | High | 1 | 5 | Q8R3Z5 | Q8R3Z5 [4 | Voltage-de | Q8R3Z5 1 | 0 | 695.027 | 2083.07 | -0.13 | 4.37 | 0.03769 | 0.0259 | -1.45534 | 4.34E-05 | 0.00642 |
| CYTO_pho | CYTO_tota | [K].ASSEG | 1xPhosph | High | 1 | 6 | Q9JKS4 | Q9JKS4 [1 | LIM domai | Q9JKS4 1x | 0 | 631.659 | 1892.96 | 0.28 | 5.28 | 0.01157 | 0.03207 | 2.77109 | 0.00014 | 0.01007 |
| CYTO_pho | CYTO_tota | [R].HSSPH | 1xPhosph | High | 1 | 9 | Q9Z1E4 | Q9Z1E4 [6 | Glycogen [ | Q9Z1E4 1 | 0 | 683.294 | 2047.87 | -0.55 | 4.27 | 0.03094 | 0.01141 | -2.71246 | 0.00014 | 0.01007 |
| CYTO_pho | CYTO_tota | [K].LSVEA | 1xPhosph | High | 1 | 19 | P07310 | P07310 [1 | Creatine k | P07310 1x | 1 | 672.13 | 2685.5 | -2.19 | 5.26 | 0.0025 | 0.00671 | 2.6834 | 0.00014 | 0.01007 |
| CYTO_pho | CYTO_tota | [K].GILAAD | 1xPhosph | High | 1 | 59 | P05064 | P05064 [2 | Fructose-b | P05064 1x | 1 | 545.051 | 2177.18 | -0.2 | 4.5 | 0.02979 | 0.04531 | 1.52096 | 0.00014 | 0.01007 |
| CYTO_pho | CYTO_tota | [R].TGEPD | 1xPhosph | High | 1 | 12 | Q9Z239 | Q9Z239 [7 | Phosphole | Q9Z239 1 | 1 | 732.004 | 2194 | -0.31 | 5.42 | 2.14629 | 5.3525 | 2.49384 | 0.00015 | 0.01007 |
| CYTO_pho | CYTO_tota | [K].RVPSP | 1xPhosph | High | 1 | 4 | Q8VDI7 | Q8VDI7 [9 | Ubiquitin- | Q8VDI7 1x | 1 | 527.984 | 1581.94 | -0.46 | 3.16 | 0.33048 | 0.45704 | 1.38294 | 0.00017 | 0.01011 |
| CYTO_pho | CYTO_tota | [K].IMSVIK | 1xPhosph | High | 1 | 2 | Q8R429 | Q8R429 [5 | Sarcoplas | Q8R429 1 | 1 | 694.375 | 2081.11 | 0.4 | 3.22 | 0.00058 | 0.00136 | 2.32018 | 0.00019 | 0.01035 |
| CYTO_pho | CYTO_tota | [R].AQSPT | 2xPhosph | High | 1 | 5 | Q91YE8 | Q91YE8 [8 | Synaptopo | Q91YE8 2 | 0 | 713.357 | 2138.06 | -0.85 | 3.49 | 1.03749 | 1.61731 | 1.55887 | 0.00034 | 0.01622 |
| CYTO_pho | CYTO_tota | [R].SAPGK | 1xPhosph | High | 1 | 2 | Q9WV06 | Q9WV06 [ | Ankyrin re | Q9WV06 1 | 1 | 866.442 | 2597.31 | 0.59 | 1.54 | 0.02331 | 0.00971 | -2.3996 | 0.00037 | 0.01622 |
| CYTO_pho | CYTO_tota | [R].EPGEP | 2xPhosph | High | 1 | 2 | Q62407 | Q62407 [5 | Striated m | Q62407 2 | 0 | 602.554 | 2407.2 | -0.36 | 2.41 | 0.1145 | 0.09583 | -1.19486 | 0.00042 | 0.01622 |
| CYTO_pho | CYTO_tota | [K].LTFDS | 1xPhosph | High | 1 | 9 | Q60932 | Q60932 [1 | Voltage-de | Q60932 1 | 0 | 1045.03 | 2089.05 | 1.34 | 4.36 | 0.02691 | 0.01613 | -1.66874 | 0.00043 | 0.01622 |
| CYTO_pho | CYTO_tota | [K].QSHSP | 1xPhosph | High | 1 | 8 | Q9ET80 | Q9ET80 [4 | Junctophil | Q9ET80 1x | 0 | 727.7 | 2181.08 | 0.36 | 5.39 | 0.07118 | 0.06055 | -1.17554 | 0.00044 | 0.01622 |
| CYTO_pho | CYTO_tota | [R].EKEISD | 1xPhosph | High | 1 | 4 | P11499 | P11499 [2 | Heat shoc | P11499 1x | 1 | 848.424 | 2543.26 | -0.13 | 3.35 | 0.00718 | 0.00526 | -1.36383 | 0.00062 | 0.02121 |
| CYTO_pho | CYTO_tota | [K].NYKNV | 1xPhosph | High | 1 | 3 | O09165 | O09165 [5 | Calseques | O09165 1 | 1 | 800.941 | 1600.87 | 0.05 | 1.6 | 0.0023 | 0.00317 | 1.37887 | 0.00066 | 0.02121 |
| CYTO_pho | CYTO_tota | [K].GATPA | 1xPhosph | High | 1 | 8 | P57776 | P57776 [1 | Elongation | P57776 1x | 2 | 838.596 | 4188.94 | 1.38 | 6.2 | 0.15271 | 0.11503 | -1.32756 | 0.00068 | 0.02121 |
| CYTO_pho | CYTO_tota | [K].GSSTP | 1xPhosph | High | 1 | 2 | Q58A65 | Q58A65 [3 | C-Jun-ami | Q58A65 1 | 1 | 737.745 | 2211.22 | -0.22 | 2.85 | 0.41429 | 0.72968 | 1.76127 | 0.00076 | 0.0226 |
| CYTO_pho | CYTO_tota | [R].SAPGK | 1xPhosph | High | 1 | 2 | Q9WV06 | Q9WV06 [ | Ankyrin re | Q9WV06 1 | 2 | 689.109 | 2753.41 | 0.59 | 2.31 | 0.02331 | 0.01196 | -1.94981 | 0.00097 | 0.0274 |
| CYTO_pho | CYTO_tota | [R].IGEDY | 1xPhosph | High | 1 | 4 | Q9WUB3 | Q9WUB3 | Glycogen | Q9WUB3 | 1 | 785.081 | 2353.23 | -0.49 | 3.8 | 0.00083 | 0.00119 | 1.4326 | 0.00106 | 0.02861 |
| CYTO_pho | CYTO_tota | [R].GGSIS | 1xPhosph | High | 1 | 46 | Q3MI48 | Q3MI48 [2 | Junctional | Q3MI48 1x | 0 | 711.001 | 2130.99 | -0.02 | 6.09 | 0.51815 | 0.39041 | -1.3272 | 0.00133 | 0.03412 |
| CYTO_pho | CYTO_tota | [R].SIDSS | 1xPhosph | High | 1 | 2 | Q9CZJ2 | Q9CZJ2 [2 | Heat shoc | Q9CZJ2 1x | 0 | 598.288 | 1195.57 | 0.7 | 1.85 | 0.28375 | 0.23102 | -1.22826 | 0.0015 | 0.03631 |
| CYTO_pho | CYTO_tota | [R].WTSP | 1xPhosph | High | 1 | 14 | Q8CHT0 | Q8CHT0 [ | Delta-1-py | Q8CHT0 1 | 0 | 549.644 | 1646.92 | 0.78 | 2.61 | 0.32183 | 0.21367 | -1.50621 | 0.00153 | 0.03631 |
| CYTO_pho | CYTO_tota | [K].SADTL | 1xPhosph | High | 1 | 6 | P06151 | P06151 [3 | L-lactate d | P06151 1x | 0 | 903.984 | 1806.96 | -2.38 | 2.99 | 0.00177 | 0.00207 | 1.16848 | 0.00176 | 0.04016 |
| CYTO_pho | CYTO_tota | [R].ATEEP | 1xPhosph | High | 1 | 21 | O54724 | O54724 [3 | Caveolae- | O54724 1 | 0 | 741.374 | 2222.11 | -0.09 | 5.2 | 0.12476 | 0.08362 | -1.49197 | 0.0021 | 0.04279 |
| CYTO_pho | CYTO_tota | [K].IQSSL | 1xPhosph | High | 1 | 4 | P41216 | P41216 [4 | Long-chai | P41216 1x | 0 | 739.417 | 1477.83 | -0.42 | 1.57 | 0.00619 | 0.00509 | -1.21764 | 0.00211 | 0.04279 |
| CYTO_pho | CYTO_tota | [K].LTFDT | 1xPhosph | High | 1 | 4 | Q60930 | Q60930 [1 | Voltage-de | Q60930 1 | 1 | 638.102 | 2549.38 | 0.67 | 3.82 | 0.01124 | 0.01438 | 1.27928 | 0.00215 | 0.04279 |
| CYTO_pho | CYTO_tota | [R].AAYFG | 1xPhosph | High | 1 | 15 | P48962 | P48962 [1 | ADP/ATP t | P48962 1x | 0 | 947.486 | 1893.96 | 0.25 | 3.73 | 0.01424 | 0.01145 | -1.24316 | 0.00217 | 0.04279 |
| CYTO_pho | CYTO_tota | [K].GTTPP | 2xPhosph | High | 1 | 2 | Q9ET80 | Q9ET80 [4 | Junctophil | Q9ET80 2x | 1 | 528.013 | 2109.03 | 0.54 | 2.32 | 0.076 | 0.05987 | -1.26942 | 0.00275 | 0.04915 |
| CYTO_pho | CYTO_tota | [K].GILAAD | 1xPhosph | High | 1 | 191 | P05064 | P05064 [2 | Fructose-b | P05064 1x | 0 | 674.365 | 2021.08 | 0 | 5.13 | 0.04583 | 0.03983 | -1.15067 | 0.00275 | 0.04915 |
| CYTO_pho | CYTO_tota | [K].SKRPI | 1xPhosph | High | 1 | 3 | Q9JKS4 | Q9JKS4 [8 | LIM domai | Q9JKS4 1x | 1 | 678.996 | 3390.95 | -0.09 | 4.59 | 0.01165 | 0.02208 | 1.8954 | 0.00289 | 0.04915 |
| CYTO_pho | CYTO_tota | [K].IEDVG | 1xPhosph | High | 1 | 20 | P11499 | P11499 [2 | Heat shoc | P11499 1x | 0 | 1092 | 2182.99 | -0.39 | 4.2 | 0.03791 | 0.02568 | -1.4762 | 0.0029 | 0.04915 |
| CYTO_pho | CYTO_tota | [R].QSSTA | 1xPhosph | High | 1 | 6 | Q7TSH2 | Q7TSH2 [6 | Phosphory | Q7TSH2 1 | 0 | 1028.84 | 3084.52 | -0.01 | 5.84 | 0.02055 | 0.02738 | 1.33244 | 0.00291 | 0.04915 |
| CYTO_pho | CYTO_tota | [R].GTGG | 1xPhosph | High | 1 | 11 | P07310 | P07310 [3 | Creatine k | P07310 1x | 0 | 793.051 | 2377.14 | 0.71 | 7.11 | 0.00051 | 0.00045 | -1.13997 | 0.00368 | 0.05881 |
| CYTO_pho | CYTO_tota | [R].LGSFG | 1xPhosph | High | 1 | 11 | Q8VHX6 | Q8VHX6 [ | Filamin-C | Q8VHX6 1 | 0 | 661.346 | 1321.68 | 0.32 | 2.76 | 0.02556 | 0.03509 | 1.37309 | 0.05975 | 0.18141 |

| MYD_phospho_index | MYD_total_index | Annotated Sequence | Modifications | Confidence (by Search Engine): Sequest HT | # Proteins | # PSMs | Master Protein Accessions | Positions in Master Proteins | Master Protein Descriptions | Modifications in Master Proteins (all Sites) | # Missed Cleavages | m/z [Da] (by Search Engine): Sequest HT | Theo. MH+ [Da] | DeltaM [ppm] (by Search Engine): Sequest HT | XCorr (by Search Engine): Sequest HT | Log2_Mean_M_SC | Log2_Mean_M_TP | Linear Difference M_TP_vs_M_SC | Welch_Test_Uncorrected_Pvalue | Welch_Test_Corrected_Pvalue |
|---|---|---|---|---|---|---|---|---|---|---|---|---|---|---|---|---|---|---|---|---|
| MYD_phospho_1536 | MYD_total_25 | [K].SKRPIPISTTAPPIQSPLPVIPHQK.[D] | 1xPhospho [S16]; 2xTMTpro [K2; K25]; 1xTMTpro [N-Term] | High | 1 | 8 | Q9JKS4 | Q9JKS4 [83-107] | LIM domain-binding protein 3 OS=Mus musculus OX=10090 GN=Ldb3 PE=1 SV=1 | Q9JKS4 1xPhospho [S98] | 1 | 739.83263 | 3695.15935 | -6.84 | 7.99 | 0.008798067 | 0.013269286 | 1.508204778 | 1.19E-05 | 0.010119554 |
| MYD_phospho_1351 | MYD_total_71 | [K].TLPQVSPKTPRPMAPK.[T] | 1xPhospho [T9]; 1xOxidation [M13]; 2xTMTpro [K8; K16]; 1xTMTpro [N-Term] | High | 1 | 6 | Q8BWB1 | Q8BWB1 [694-709] | Synaptopodin 2-like protein OS=Mus musculus OX=10090 GN=Synpo2l PE=1 SV=1 | Q8BWB1 1xPhospho [T702] | 1 | 675.13611 | 2697.52307 | -0.17 | 5.98 | 0.016465902 | 0.028326162 | 1.7203933 | 4.88E-05 | 0.020772792 |
| MYD_phospho_813 | MYD_total_20 | [R].ETSPEQPGSEVHTK.[K] | 1xPhospho [S9]; 1xTMTpro [K14]; 1xTMTpro [N-Term] | High | 1 | 3 | P31001 | P31001 [429-442] | Desmin OS=Mus musculus OX=10090 GN=Des PE=1 SV=3 | P31001 1xPhospho [S437] | 1 | 569.0408 | 2273.14196 | -0.21 | 3.54 | 0.00284765 | 0.003199486 | -1.284689164 | 8.20E-05 | 0.033360384 |
| MYD_phospho_1179 | MYD_total_91 | [R].VPTVPPRPGSSLSGSIENLESEAVFEAK.[E] | 3xPhospho [S13; S]; 1xTMTpro [K28]; 1xTMTpro [N-Term] | High | 1 | 10 | Q62407 | Q62407 [2313-2340] | Striated muscle-specific serine/threonine-protein kinase OS=Mus musculus OX=10090 GN=Speg PE=1 SV=2 | Q62407 3xPhospho [S2322; S2323; S2325; S2327; S2333] | 0 | 937.20624 | 3745.80238 | 0.2 | 4.91 | 0.025542299 | 0.030877748 | 1.208886804 | 0.000169671 | 0.036097426 |
| MYD_phospho_1485 | MYD_total_13 | [R].ILASDKPYILAEELR.[R] | 1xPhospho [S4]; 1xTMTpro [K6]; 1xTMTpro [N-Term] | High | 1 | 3 | Q9JI91 | Q9JI91 [837-851] | Alpha-actinin-3 OS=Mus musculus OX=10090 GN=Actn3 PE=1 SV=2 | Q9JI91 1xPhospho [S840] | 0 | 807.42174 | 2419.24933 | 0.55 | 4.71 | 0.002746363 | 0.001588309 | -1.729033699 | 0.000245403 | 0.038662835 |
| MYD_phospho_438 | MYD_total_1 | [R].APTPSPVR.[S] | 2xPhospho [T3; S5]; 1xTMTpro [N-Term] | High | 1 | 7 | A2ASS6 | A2ASS6 [297-304] | Titin OS=Mus musculus OX=10090 GN=Ttn PE=1 SV=1 | A2ASS6 2xPhospho [T299; S301] | 0 | 430.20551 | 1288.60229 | -0.25 | 2.41 | 0.000274345 | 0.000167274 | -1.640098052 | 0.000285084 | 0.038662835 |
| MYD_phospho_859 | MYD_total_8 | [R].KLVIIESDLER.[A] | 1xPhospho [S7]; 1xTMTpro [K1]; 1xTMTpro [N-Term] | High | 1 | 5 | P58771 | P58771 [168-178] | Tropomyosin alpha-1 chain OS=Mus musculus OX=10090 GN=Tpm1 PE=1 SV=1 | P58771 1xPhospho [S174] | 1 | 668.38595 | 2003.14336 | -0.03 | 3.45 | 0.003883196 | 0.003416858 | -1.136188968 | 0.000319036 | 0.038662835 |
| MYD_phospho_500 | MYD_total_1 | [R].SPTPPSIAAK.[A] | 1xPhospho [T/S]; 1xTMTpro [K10]; 1xTMTpro [N-Term] | High | 1 | 49 | A2ASS6 | A2ASS6 [264-273] | Titin OS=Mus musculus OX=10090 GN=Ttn PE=1 SV=1 | A2ASS6 1xPhospho [S264; T266; S269] | 0 | 828.96419 | 1656.92174 | -0.38 | 3.16 | 0.000799358 | 0.000632563 | -1.263533363 | 0.000511316 | 0.054380557 |

| MYO_phospho | MYO_total_inde | Annotated Seq | Modifications | Confidence (by | # Proteins | # PSMs | Master Protein | Positions in Ma | Master Protein | Modifications i | # Missed Cleav | m/z [Da] (by Se | Theo. MH+ [Da | DeltaM [ppm] ( | XCorr (by Searc | Log2_Mean_M | Log2_Mean_M | **Linear_Differe** | **Welch_Ttest_** | Welch_Ttest_Corrected_Pvalue |
|---|---|---|---|---|---|---|---|---|---|---|---|---|---|---|---|---|---|---|---|---|
| MYO_phospho | MYO_total_25 | [K].ASSEGAQG | 1xPhospho [S]; | High | 1 | 9 | Q9JKS4 | Q9JKS4 [169-1 | LIM domain-bir | Q9JKS4 1xPhos | 0 | 631.65855 | 1892.96104 | 0.03 | 4.3 | 0.004043853 | 0.011625322 | 2.874813631 | 0.001154884 | 0.060894164 |
| MYO_phospho | MYO_total_1 | [K].KTPSPIEAE | 1xPhospho [S4 | High | 1 | 9 | A2ASS6 | A2ASS6 [12868 | Titin OS=Mus m | A2ASS6 1xPhos | 1 | 605.99984 | 1815.98613 | -0.64 | 2.7 | 0.000638476 | 0.00118998 | 1.863781945 | 0.00303239 | 0.066233234 |
| MYO_phospho | MYO_total_1 | [K].KLSDHSVE | 1xPhospho [S3 | High | 1 | 23 | A2ASS6 | A2ASS6 [6879- | Titin OS=Mus m | A2ASS6 1xPhos | 1 | 548.0592 | 2189.21474 | 0.11 | 4.84 | 0.001017701 | 0.001755352 | 1.724820878 | 0.001982884 | 0.06490132 |
| MYO_phospho | MYO_total_71 | [K].TLPQVSPKT | 1xPhospho [T9 | High | 1 | 6 | Q8BWB1 | Q8BWB1 [694- | Synaptopodin 2 | Q8BWB1 1xPh | 1 | 675.13611 | 2697.52307 | -0.17 | 5.86 | 0.016465902 | 0.028326162 | 1.7202922 | 4.88E-05 | 0.020772792 |
| MYO_phospho | MYO_total_22 | [K].SSKLIMELT | 1xPhospho [S2 | High | 1 | 6 | Q9JK37 | Q9JK37 [15-27 | Myozenin-1 OS | Q9JK37 1xPhos | 1 | 685.03876 | 2053.10085 | 0.42 | 3.78 | 0.018069124 | 0.029703891 | 1.643903185 | 0.015636026 | 0.150513801 |
| MYO_phospho | MYO_total_14 | [R].TTSTLQREE | 1xPhospho [T1 | High | 1 | 9 | Q62234 | Q62234 [207-2 | Myomesin-1 OS | Q62234 1xPho | 1 | 565.29471 | 2258.15611 | 0.4 | 3.33 | 0.008203568 | 0.013063426 | 1.59240773 | 0.022039386 | 0.174715138 |
| MYO_phospho | MYO_total_22 | [K].SSKLIMELT | 1xPhospho [S2 | High | 1 | 1 | Q9JK37 | Q9JK37 [15-27 | Myozenin-1 OS | Q9JK37 1xPhos | 1 | 679.70687 | 2037.10593 | 0.06 | 2.94 | 0.004672305 | 0.007273757 | 1.556781142 | 0.033261404 | 0.196565657 |
| MYO_phospho | MYO_total_13 | [K].MLDAEDIVN | 1xPhospho [T1 | High | 1 | 14 | Q9JI91 | Q9JI91 [228-24 | Alpha-actinin-2 | Q9JI91 1xPhos | 0 | 637.57316 | 2547.26574 | 1.99 | 5.28 | 0.005047245 | 0.007727192 | 1.530972243 | 0.002749349 | 0.066233234 |
| MYO_phospho | MYO_total_25 | [K].AAQSQLSQ | 1xPhospho [S7 | High | 1 | 5 | Q9JKS4 | Q9JKS4 [38-68 | LIM domain-bir | Q9JKS4 1xPhos | 0 | 990.49921 | 3958.97129 | 0.93 | 7.6 | 0.004086563 | 0.006194518 | 1.515825882 | 0.007622666 | 0.104627235 |
| MYO_phospho | MYO_total_25 | [K].SKRPIPISTT | 1xPhospho [S1 | High | 1 | 8 | Q9JKS4 | Q9JKS4 [83-10 | LIM domain-bir | Q9JKS4 1xPhos | 1 | 739.83263 | 3695.15935 | -6.84 | 7.99 | 0.008798067 | 0.013269286 | 1.508204778 | 1.19E-05 | 0.010119554 |
| MYO_phospho | MYO_total_29 | [R].SSSRAEAN | 1xPhospho [S]; | High | 1 | 49 | Q9JIF9 | Q9JIF9 [229-24 | Myotilin OS=Mu | Q9JIF9 1xPhos | 1 | 813.06686 | 2437.18518 | 0.35 | 5.55 | 0.122860619 | 0.184503401 | 1.501729375 | 0.004390436 | 0.079715601 |
| MYO_phospho | MYO_total_31 | [R].SVTSPISDF | 2xPhospho [T/Y | High | 1 | 24 | Q91YE8 | Q91YE8 [600-6 | Synaptopodin-2 | Q91YE8 2xPho | 0 | 1088.84692 | 3264.52264 | 1.1 | 6.91 | 0.018601119 | 0.026625135 | 1.431372742 | 0.048461906 | 0.233853685 |
| MYO_phospho | MYO_total_314 | [R].EMEKSFDE | 1xPhospho [S5 | High | 1 | 7 | Q8K4L3 | Q8K4L3 [662-6 | Supervillin OS= | Q8K4L3 1xPho | 1 | 647.08369 | 2585.31386 | -0.36 | 2.68 | 0.287062067 | 0.406387061 | 1.415676637 | 0.002879203 | 0.066233234 |
| MYO_phospho | MYO_total_71 | [R].GPARPSLP | 2xPhospho [Y/S | High | 1 | 13 | Q8BWB1 | Q8BWB1 [158- | Synaptopodin 2 | Q8BWB1 2xPh | 0 | 1194.93237 | 3582.7778 | 1.33 | 4.66 | 0.045047634 | 0.063288177 | 1.404916793 | 0.010094129 | 0.122722339 |
| MYO_phospho | MYO_total_29 | [R].SRSSSRAEA | 2xPhospho [S]; | High | 1 | 11 | Q9JIF9 | Q9JIF9 [227-24 | Myotilin OS=Mu | Q9JIF9 2xPhos | 2 | 690.8269 | 2760.28465 | 0.41 | 5.95 | 0.013228251 | 0.018441639 | 1.394110202 | 0.001561411 | 0.06490132 |
| MYO_phospho | MYO_total_31 | [R].MQSSVSES | 2xPhospho [S]; | High | 1 | 20 | Q91YE8 | Q91YE8 [538-5 | Synaptopodin-2 | Q91YE8 2xPho | 0 | 686.27423 | 2056.80774 | 0.19 | 3.78 | 0.018372874 | 0.025600155 | 1.39336691 | 0.006622042 | 0.102461043 |
| MYO_phospho | MYO_total_314 | [R].KFSLKEFGE | 1xPhospho [S3 | High | 1 | 2 | Q8K4L3 | Q8K4L3 [1009- | Supervillin OS= | Q8K4L3 1xPho | 2 | 1084.24479 | 3250.71768 | 0.66 | 4.86 | 0.803489833 | 1.118783019 | 1.392404698 | 0.022836493 | 0.178292255 |
| MYO_phospho | MYO_total_91 | [R].LSALGRSP | 1xPhospho [S7 | High | 1 | 6 | Q62407 | Q62407 [388-3 | Striated muscle | Q62407 1xPho | 1 | 447.58365 | 1340.73707 | -0.49 | 3.24 | 0.099147504 | 0.136342102 | 1.375144065 | 0.008196647 | 0.108989787 |
| MYO_phospho | MYO_total_39 | [R].KISQTAQTY | 1xPhospho [S3 | High | 1 | 13 | E9PZQ0 | E9PZQ0 [2842- | Ryanodine rece | E9PZQ0 1xPho | 1 | 524.78125 | 2096.10329 | -0.06 | 2.65 | 0.098270937 | 0.134299535 | 1.366625151 | 0.022173014 | 0.174715138 |
| MYO_phospho | MYO_total_25 | [K].RPIPISTTAP | 1xPhospho [S1 | High | 1 | 13 | Q9JKS4 | Q9JKS4 [85-10 | LIM domain-bir | Q9JKS4 1xPhos | 0 | 794.71134 | 3175.82521 | -0.53 | 7.71 | 0.037058399 | 0.050123591 | 1.352556839 | 0.008777825 | 0.113180742 |
| MYO_phospho | MYO_total_1 | [R].APTPSPVR | 4xPhospho [T3 | High | 1 | 43 | A2ASS6 | A2ASS6 [297-3 | Titin OS=Mus m | A2ASS6 4xPhos | 1 | 701.63025 | 2102.87987 | -1.75 | 5.93 | 0.001120502 | 0.000826565 | -1.35561171 | 0.001727867 | 0.06490132 |
| MYO_phospho | MYO_total_20 | [R].APSYGAGE | 1xPhospho [S/Y | High | 1 | 20 | P31001 | P31001 [79-10 | Desmin OS=Mu | P31001 1xPhos | 0 | 1047.18194 | 3139.53083 | 0.14 | 8.48 | 0.003046851 | 0.002247037 | -1.35594176 | 0.011099374 | 0.124283785 |
| MYO_phospho | MYO_total_1 | [R].IELSPSMEA | 1xPhospho [S6 | High | 1 | 4 | A2ASS6 | A2ASS6 [2075- | Titin OS=Mus m | A2ASS6 1xPhos | 0 | 630.66978 | 1889.99392 | 0.46 | 3.14 | 0.000626389 | 0.000461051 | -1.35861164 | 0.015741161 | 0.150513801 |
| MYO_phospho | MYO_total_8 | [K].LEKSIDDLE | 1xPhospho [S4 | High | 1 | 2 | P58771 | P58771 [249-2 | Tropomyosin al | P58771 1xPhos | 1 | 866.4466 | 2597.3243 | 0.37 | 5.06 | 0.001467617 | 0.001078444 | -1.36086543 | 0.026869066 | 0.195432267 |
| MYO_phospho | MYO_total_1 | [R].SLGDMSDE | 1xPhospho [S]; | High | 1 | 5 | A2ASS6 | A2ASS6 [33875 | Titin OS=Mus m | A2ASS6 1xPhos | 0 | 1019.51051 | 3056.51046 | 2.13 | 8.37 | 9.23E-05 | 6.76E-05 | -1.36481556 | 0.029818363 | 0.196565657 |
| MYO_phospho | MYO_total_1 | [R].QQSPSPIR | 3xPhospho [S5 | High | 1 | 20 | A2ASS6 | A2ASS6 [279-2 | Titin OS=Mus m | A2ASS6 3xPhos | 1 | 739.66943 | 2216.99391 | -0.08 | 5.12 | 0.000603697 | 0.000439186 | -1.37458112 | 0.001216452 | 0.060894164 |
| MYO_phospho | MYO_total_369 | [R].LGSRPSIQE | 2xPhospho [S3 | High | 1 | 2 | O70373 | O70373 [203-2 | Xin actin-bindi | O70373 2xPho | 0 | 758.7121 | 2274.12154 | 0.09 | 3.27 | 0.383862203 | 0.278822254 | -1.37672728 | 0.038851946 | 0.216098077 |
| MYO_phospho | MYO_total_1 | [R].APTPSPVR. | 1xPhospho [T/S | High | 1 | 5 | A2ASS6 | A2ASS6 [297-3 | Titin OS=Mus m | A2ASS6 1xPhos | 0 | 403.54973 | 1208.63595 | -1.09 | 2.21 | 0.0001184 | 8.59E-05 | -1.37782745 | 0.001018069 | 0.060894164 |
| MYO_phospho | MYO_total_1 | [R].TASPHFTVS | 1xPhospho [T/S | High | 1 | 14 | A2ASS6 | A2ASS6 [812-8 | Titin OS=Mus m | A2ASS6 1xPhos | 0 | 588.31747 | 1762.93846 | -0.34 | 4.3 | 0.000248307 | 0.000179797 | -1.38103931 | 0.034969718 | 0.201305386 |
| MYO_phospho | MYO_total_13 | [R].ISSSNPYST | 1xPhospho [S/Y | High | 1 | 12 | Q9JI91 | Q9JI91 [593-60 | Alpha-actinin-2 | Q9JI91 1xPhos | 0 | 734.01117 | 2200.0164 | 1.16 | 3.84 | 0.005772927 | 0.004160499 | -1.38755626 | 0.005436143 | 0.094411375 |
| MYO_phospho | MYO_total_1 | [R].TPSPDYDL | 1xPhospho [S/T | High | 1 | 18 | A2ASS6 | A2ASS6 [33859 | Titin OS=Mus m | A2ASS6 1xPhos | 0 | 646.29368 | 1936.86893 | -1.26 | 3.74 | 0.001228876 | 0.000882471 | -1.39253969 | 0.001037637 | 0.060894164 |
| MYO_phospho | MYO_total_1 | [R].YSSPPAHV | 1xPhospho [S/Y | High | 1 | 143 | A2ASS6 | A2ASS6 [33931 | Titin OS=Mus m | A2ASS6 1xPhos | 0 | 558.63544 | 1673.89078 | 0.59 | 2.62 | 0.002692094 | 0.001928542 | -1.39592244 | 0.018013937 | 0.159686046 |
| MYO_phospho | MYO_total_40 | [R].LRLSPSPTS | 2xPhospho [S4 | High | 1 | 3 | P48678 | P48678 [387-3 | Prelamin-A/C O | P48678 2xPhos | 1 | 569.28339 | 1705.83587 | -0.15 | 2.96 | 0.016169474 | 0.011525076 | -1.40298194 | 0.01102291 | 0.124283785 |
| MYO_phospho | MYO_total_1 | [R].RTPSPDYD | 1xPhospho [S/T | High | 1 | 24 | A2ASS6 | A2ASS6 [33858 | Titin OS=Mus m | A2ASS6 1xPhos | 1 | 698.32735 | 2092.97004 | -1.22 | 5.24 | 0.001060172 | 0.000754796 | -1.40458203 | 0.002657034 | 0.066233234 |
| MYO_phospho | MYO_total_3 | [K].TEGGATVT | 1xPhospho [S/T | High | 1 | 5 | Q5SX39 | Q5SX39 [64-87 | Myosin-4 OS=M | Q5SX39 1xPho | 2 | 727.97614 | 3635.84844 | 0.87 | 2.8 | 0.000386792 | 0.000275277 | -1.40510294 | 0.041164892 | 0.216324716 |
| MYO_phospho | MYO_total_1 | [R].LSEYKSELD | 1xPhospho [S6 | High | 1 | 1 | A2ASS6 | A2ASS6 [34004 | Titin OS=Mus m | A2ASS6 1xPhos | 1 | 867.78248 | 2601.33393 | -0.41 | 2.35 | 0.000125657 | 8.91E-05 | -1.41005608 | 0.00412356 | 0.077981102 |
| MYO_phospho | MYO_total_1 | [R].QQSPSPIR | 2xPhospho [S]; | High | 1 | 14 | A2ASS6 | A2ASS6 [279-2 | Titin OS=Mus m | A2ASS6 2xPhos | 1 | 535.01233 | 2137.02758 | -0.05 | 2.83 | 0.000189181 | 0.000130259 | -1.45234235 | 0.002468773 | 0.066233234 |
| MYO_phospho | MYO_total_419 | [R].KDLFANTVL | 1xPhospho [T7 | High | 1 | 2 | Q8BFZ3 | Q8BFZ3 [292-3 | Beta-actin-like | Q8BFZ3 1xPho | 1 | 905.106 | 2713.32275 | -7.1 | 2.62 | 1.142847039 | 0.786579333 | -1.45293296 | 0.014768153 | 0.146136026 |
| MYO_phospho | MYO_total_1 | [R].APTPSPVR | 3xPhospho [T3 | High | 1 | 10 | A2ASS6 | A2ASS6 [297-3 | Titin OS=Mus m | A2ASS6 3xPhos | 1 | 674.97614 | 2022.91354 | 0.16 | 5.98 | 0.000358667 | 0.000243698 | -1.4717652 | 0.007058504 | 0.103710431 |
| MYO_phospho | MYO_total_3 | [K].TEGGATVT | 1xPhospho [T/S | High | 1 | 10 | Q5SX39 | Q5SX39 [64-84 | Myosin-4 OS=M | Q5SX39 1xPho | 1 | 975.82369 | 2925.45606 | 0.16 | 3.25 | 0.000391762 | 0.00026226 | -1.49379277 | 0.021900741 | 0.174715138 |
| MYO_phospho | MYO_total_1 | [R].RRTPSPDY | 2xPhospho [T3 | High | 1 | 42 | A2ASS6 | A2ASS6 [33857 | Titin OS=Mus m | A2ASS6 2xPhos | 2 | 583.01501 | 2329.03748 | 0.32 | 5.86 | 0.001337188 | 0.000886165 | -1.5089597 | 0.001324734 | 0.06263049 |
| MYO_phospho | MYO_total_71 | [R].SPSPGELQ | 1xPhospho [S]; | High | 1 | 13 | Q8BWB1 | Q8BWB1 [92-1 | Synaptopodin 2 | Q8BWB1 1xPh | 0 | 980.27041 | 3918.05846 | 0.34 | 9.48 | 0.022065017 | 0.014503705 | -1.52133655 | 0.013906709 | 0.1443245 |
| MYO_phospho | MYO_total_9 | [R].VGEPSMSA | 1xPhospho [S]; | High | 1 | 19 | O88990 | O88990 [314-3 | Alpha-actinin-3 | O88990 1xPho | 0 | 536.90747 | 1608.70821 | -0.22 | 3.57 | 0.008266958 | 0.00538819 | -1.53427382 | 0.015558259 | 0.150513801 |
| MYO_phospho | MYO_total_31 | [R].TAKPFLGSN | 1xPhospho [S/T | High | 1 | 21 | Q91YE8 | Q91YE8 [580-5 | Synaptopodin-2 | Q91YE8 1xPho | 0 | 840.08767 | 2518.24846 | 0 | 5.54 | 0.0334343 | 0.021082767 | -1.5858592 | 0.048233727 | 0.233853685 |
| MYO_phospho | MYO_total_1 | [K].YTFESDKGL | 1xPhospho [T/Y | High | 1 | 4 | A2ASS6 | A2ASS6 [33527 | Titin OS=Mus m | A2ASS6 1xPhos | 1 | 950.46882 | 3798.79583 | 15.17 | 3.23 | 0.000127212 | 8.00E-05 | -1.5909368 | 0.01161687 | 0.126526191 |
| MYO_phospho | MYO_total_1 | [R].RRTPSPDY | 1xPhospho [T/S | High | 1 | 13 | A2ASS6 | A2ASS6 [33857 | Titin OS=Mus m | A2ASS6 1xPhos | 2 | 563.02308 | 2249.07115 | -0.29 | 5.33 | 0.000351988 | 0.000216944 | -1.62248134 | 0.002533957 | 0.066233234 |
| MYO_phospho | MYO_total_71 | [R].SPPDSEAY | 1xPhospho [Y/S | High | 1 | 26 | Q8BWB1 | Q8BWB1 [127- | Synaptopodin 2 | Q8BWB1 1xPh | 0 | 875.65643 | 3499.60339 | 0.15 | 6.1 | 0.061004101 | 0.037318212 | -1.63470053 | 0.010671508 | 0.122722339 |
| MYO_phospho | MYO_total_1 | [R].APTPSPVR. | 2xPhospho [T3 | High | 1 | 7 | A2ASS6 | A2ASS6 [297-3 | Titin OS=Mus m | A2ASS6 2xPhos | 0 | 430.20551 | 1288.60229 | -0.25 | 2.41 | 0.000274345 | 0.000167274 | -1.64009805 | 0.000285084 | 0.038662825 |
| MYO_phospho | MYO_total_445 | [R].SSPPKDSE | 1xPhospho [S]; | High | 1 | 6 | Q9D2N4 | Q9D2N4 [362- | Dystrobrevin al | Q9D2N4 1xPh | 1 | 711.6958 | 2133.07205 | 0.38 | 2.92 | 1.188806435 | 0.722717462 | -1.64491174 | 0.016613971 | 0.15367923 |
| MYO_phospho | MYO_total_326 | [R].YHGHSMS | 2xPhospho [Y/S | High | 1 | 4 | P35486 | P35486 [289-3 | Pyruvate dehyd | P35486 2xPhos | 0 | 691.6134 | 2072.82577 | -0.06 | 3.54 | 0.324827696 | 0.195538129 | -1.66119875 | 0.029679374 | 0.196565657 |
| MYO_phospho | MYO_total_10 | [R].RAGAEGSS | 1xPhospho [S]; | High | 1 | 2 | P97457 | P97457 [9-31] | Myosin regulat | P97457 1xPhos | 1 | 993.12909 | 2977.37222 | 0.17 | 5.58 | 0.002774514 | 0.001657033 | -1.67438627 | 0.032363779 | 0.196565657 |
| MYO_phospho | MYO_total_326 | [R].YHGHSMS | 1xPhospho [S/Y | High | 1 | 11 | P35486 | P35486 [289-3 | Pyruvate dehyd | P35486 1xPhos | 0 | 664.95814 | 1992.85944 | 0.21 | 2.15 | 0.324827696 | 0.191531907 | -1.6959456 | 0.031186125 | 0.196565657 |
| MYO_phospho | MYO_total_13 | [R].ILASDKPYIL | 1xPhospho [S4 | High | 1 | 2 | Q9JI91 | Q9JI91 [837-85 | Alpha-actinin-2 | Q9JI91 1xPhos | 0 | 807.12174 | 2419.34933 | 0.55 | 4.71 | 0.002746363 | 0.001589309 | -1.7280237 | 0.000245403 | 0.038662825 |
| MYO_phospho | MYO_total_1 | [R].VKSPETVKS | 1xPhospho [S9 | High | 1 | 7 | A2ASS6 | A2ASS6 [34462 | Titin OS=Mus m | A2ASS6 1xPhos | 2 | 548.82719 | 2192.28718 | -0.12 | 5.84 | 0.000382913 | 0.000202077 | -1.89488456 | 0.034643787 | 0.201305386 |
| MYO_phospho | MYO_total_1 | [K].RVKSPEPVT | 1xPhospho [S4 | High | 1 | 18 | A2ASS6 | A2ASS6 [34473 | Titin OS=Mus m | A2ASS6 1xPhos | 2 | 538.30479 | 2150.19786 | -0.25 | 3.46 | 0.000700094 | 0.000355003 | -1.97207907 | 0.025885656 | 0.193234155 |
| MYO_phospho | MYO_total_69 | [R].TLSDYNIQK | 1xPhospho [S1 | High | 4 | 2 | P62984 | P62984 [55-72 | Ubiquitin-60S r | P62984 1xPhos | 1 | 705.38942 | 2818.53596 | -0.04 | 5.94 | 0.071455557 | 0.016381672 | -4.36192096 | 0.007504935 | 0.104627235 |

TABLE S3. PKCα-regulated or mechanotransduction-relevant phosphoproteins altered by RNAi-sh16

| Protein | Fraction | Mouse Phosphosite(s) Detected | Fold Δ (TR/SC) | Direction | Welch_Ttest_ Uncorrected_ p-value | Welch_Ttest_ Corrected_p-value | PKCα Substrate? | Role in Muscle & Mechanotransduction | Relationship to LDB3–PKCα–FLNC axis | References |
|---|---|---|---|---|---|---|---|---|---|---|
| **Filamin-C (FLNC)** | CYTO | **S2234, S2237** | **1.37** | ↑ | 0.059753789* | 0.18* | **Yes** (PKCα prevents calpain cleavage of FLNC) | Main **Z-disc mechanosensor**; stabilizes sarcomeres | Direct substrate of PKCα; interacts physically with LDB3. | [1–3] |
| **LDB3 (Cypher/ZASP)** | CYTO | S179 | **2.77** | ↑ | 0.00014 | 0.01 | **PKCα-binding scaffold** | Z-disc anchoring protein; recruits PKCα and FLNC | Central hub of LDB3–PKCα–FLNC signaling | [2,4,5] |
| LDB3 | CYTO | S98 | **1.9** | ↑ | 0.00289 | 0.049 | Scaffold (LIM domain) | Z-disc anchoring protein; recruits PKCα and FLNC | ↑ phosphorylation indicates restored PKCα positioning | [2,4,5] |
| **LDB3 (MYO fraction)** | MYO | S98 | **1.51** | ↑ | 0.00001 | 0.01 | Scaffold (LIM domain) | Myofibrillar scaffold state | Shows compartment-specific PKCα rescue | [2] |
| **FXYD1 (Phospholemman)** | CYTO | S82/S83 | **2.49** | ↑ | 0.00015 | 0.01 | **Yes (canonical PKCα substrate)** | Regulates Na /K$^{+}$ pump; modulates $Ca^{2}$ loading | PKCα normalization restores membrane excitability linked to Z-disc tension | [6–7] |
| **MARCKS** | CYTO | S138/S141/T143 | **2.12** | ↑ | 0.00004 | 0.006 | **Yes (classic PKC substrate)** | Actin–membrane tethering | Reorganization of membrane–cytoskeleton on tension after PKCα rescue | [8] |
| **CKM (Creatine kinase M-type)** | CYTO | S164 | **2.68** | ↑ | 0.00014 | 0.01 | **Yes** | Maintains ATP buffering near myofibrils | Local metabolic rescue required for FLNC stability | [9,10] |
| CKM | CYTO | S372 | **1.64** | ↑ | 0.00003 | 0.006 | Yes | Maintains ATP buffering near myofibrils | Reflects PKCα activation | [9,10] |
| **ATP2A1 (SERCA1)** | CYTO | S547 | **2.32** | ↑ | 0.00019 | 0.01 | **PKC-regulated** (via PLN/SERCA) | $Ca^{2}$ reuptake into SR | Improved $Ca^{2}$ cycling reduces FLNC mechanical strain | [11] |
| **CASQ1 (Calsequestrin-1)** | CYTO | Y57 | **1.38** | ↑ | 0.00066 | 0.021 | PKC-regulated indirectly | SR $Ca^{2}$ buffering | Helps re-establish FLNC-compatible $Ca^{2}$ environment | [12] |
| **SPEG** | MYO | S2322–2333 cluster | **1.21** | ↑ | 0.00017 | 0.036 | PKC-linked remodeling kinase | Myofibrillar remodeling; Z-disc/I-band maintenance | Activated only when Z-disc tension normalizes with restored FLNC | [13] |
| **Synaptopodin-2 / Synpo2l** | MYO | T702 | **1.72** | ↑ | 0.00005 | 0.021 | PKC-responsive | Actin–filamin crosslinking | Mechanotransduction partner of FLNC | [1] |
| **Desmin** | MYO | S437 | **−1.29** | ↓ | 0.00008 | 0.023 | PKC-modulated | IF network linking Z-discs | ↓ phosphorylation = reduced cytoskeletal stress after rescue | [14] |
| **Titin (TTN)** | MYO | T299/S301 | **−1.64** | ↓ | 0.00029 | 0.038 | **PKCα phosphorylates titin's Z-disc/I-band regions** | Passive elasticity; mechanotransducer | ↓ indicates resolution of titin overstretch stress | [1,15] |
| Titin | MYO | S264/T266/S269 | **−1.26** | ↓ | 0.00080 | 0.054 | Yes | Passive elasticity; mechanotransducer | ↓ phosphorylation matches restored Z-disc integrity | [1,15] |
| **α-Actinin-2 (ACTN2)** | MYO | S840 | **−1.73** | ↓ | 0.00025 | 0.039 | Not PKCα substrate, but PKC-modulated | Crosslinks actin to Z-disc; binds LDB3 & FLNC | ↓ = reduced stress remodeling at the Z-disc | [4,16] |
| **Tropomyosin 1 (TPM1)** | MYO | S174 | **−1.14** | ↓ | 0.00388 | 0.038 | Regulated by PKC in some contexts | Thin filament regulatory protein | ↓ = normalization of thin filament tension | [17] |
| **GYS1 (Glycogen synthase 1)** | CYTO | S652–S657 | **−2.71** | ↓ | 0.00014 | 0.01 | **Yes** | Glycogen metabolism | ↓ phosphorylation indicates metabolic normalization | [18] |

*For FLNC S2234/S2237, Welch's t-test yielded p = 0.0598; equal-variance t-test yielded p = 0.0457.

? PKCα Substrate assignments are based on published experimental evidence or curated kinase/phosphosite databases (PhosphoSitePlus, UniProt, KEA3). Indirect terms ('PKC-responsive', PKC-modulated') indicate pathway-level association rather than confirmed kinase - substrate activity.

**Table S4.** Enrichr-KG results for proteins with treatment-responsive phosphosites in sh16–treated *Ldb3*$^{\mathrm{Ala165Val/+}}$ mice muscle

| Library | Enriched term | p value | BH-FDR q value | z score | combined score | Overlap genes |
|---|---|---|---|---|---|---|
| MGI_Mammalian_Phenotype_Level_4_2021 | impaired skeletal muscle contractility MP:0002841 | 1.38e-13 | 3.475e-11 | 185 | 5467 | CACNB1;DES;CKM;ATP2A1;CASQ1;JPH1;TTN |
| GO_Biological_Process_2021 | sarcomere organization (GO:0045214) | 1.16e-09 | 4.481e-07 | 144 | 2973 | SYNPO2L;ACTN2;TPM1;CASQ1;TTN |
| GO_Biological_Process_2021 | muscle contraction (GO:0006936) | 2.28e-09 | 4.481e-07 | 40.7 | 809.1 | DES;ACTN2;FXYD1;TPM1;ANKRD2;ALDOA;TTN |
| MGI_Mammalian_Phenotype_Level_4_2021 | increased variability of skeletal muscle fiber size MP:0009403 | 6.72e-09 | 8.466e-07 | 97.7 | 1839 | DES;LDB3;PYGM;FLNC;TTN |
| MGI_Mammalian_Phenotype_Level_4_2021 | abnormal muscle physiology MP:0002106 | 1.1e-08 | 9.203e-07 | 48.4 | 886.9 | CACNB1;JSRP1;CKM;ATP2A1;CASQ1;TTN |
| GO_Biological_Process_2021 | myofibril assembly (GO:0030239) | 1.12e-08 | 1.464e-06 | 87.4 | 1600 | SYNPO2L;ACTN2;TPM1;CASQ1;TTN |
| MGI_Mammalian_Phenotype_Level_4_2021 | increased heart weight MP:0002833 | 5.47e-08 | 3.445e-06 | 25.1 | 419.5 | GYS1;LDHA;DES;FXYD1;SPEG;LDB3;SLC25A4 |
| GO_Biological_Process_2021 | actomyosin structure organization (GO:0031032) | 2.06e-07 | 2.023e-05 | 46.7 | 719 | SYNPO2L;ACTN2;TPM1;CASQ1;TTN |
| MGI_Mammalian_Phenotype_Level_4_2021 | abnormal skeletal muscle morphology MP:0000759 | 2.2e-07 | 1.108e-05 | 46 | 705.9 | DES;LDB3;ANKRD2;FLNC;TTN |
| KEGG_2021_Human | Hypertrophic cardiomyopathy | 4.81e-07 | 2.259e-05 | 39 | 567 | CACNB1;DES;TPM1;ATP2A1;TTN |
| GO_Biological_Process_2021 | actin-myosin filament sliding (GO:0033275) | 5.56e-07 | 3.183e-05 | 75.6 | 1089 | DES;ACTN2;TPM1;TTN |
| KEGG_2021_Human | Dilated cardiomyopathy | 6.64e-07 | 2.259e-05 | 36.4 | 517.8 | CACNB1;DES;TPM1;ATP2A1;TTN |
| Reactome_2022 | Muscle Contraction R-HSA-397014 | 1.05e-06 | 0.0001577 | 21.5 | 296.4 | CACNB1;DES;ACTN2;FXYD1;TPM1;ATP2A1 |
| KEGG_2021_Human | Calcium signaling pathway | 3.4e-06 | 7.711e-05 | 17.4 | 219.7 | VDAC2;PHKB;VDAC1;ATP2A1;CASQ1;SLC25A4 |
| KEGG_2021_Human | Necroptosis | 7.98e-06 | 0.0001321 | 21.4 | 251.7 | HSP90AB1;VDAC2;VDAC1;PYGM;SLC25A4 |
| Reactome_2022 | Glycogen Metabolism R-HSA-8982491 | 9.69e-06 | 0.0007267 | 89 | 1028 | GYS1;PHKB;PYGM |
| KEGG_2021_Human | Arrhythmogenic right ventricular cardiomyopathy | 9.71e-06 | 0.0001321 | 35.2 | 405.8 | CACNB1;DES;ACTN2;ATP2A1 |
| Reactome_2022 | Striated Muscle Contraction R-HSA-390522 | 2.58e-05 | 0.001292 | 62.3 | 658.1 | DES;ACTN2;TPM1 |
| Reactome_2022 | Metabolism R-HSA-1430728 | 0.000114 | 0.004259 | 4.59 | 41.71 | ALDH4A1;GYS1;MARCKS;LDHA;HSP90AB1;CKM;CA3;ACSL1;VDAC2;PHKB;VDAC1;PYGM |
| Reactome_2022 | Glycogen Breakdown (Glycogenolysis) R-HSA-70221 | 0.000267 | 0.008015 | 101 | 829.1 | PHKB;PYGM |

Enrichr-KG was run using 35 significantly altered protein derived from TMT-based phosphoproteomics (phosphosite abundances normalized to the corresponding total protein). This table reports the complete Enrichr output as exported (all returned terms). p values are nominal enrichment p values and BH-FDR q values are Benjamini–Hochberg adjusted within each library.
